# A framework to detect positive selection using variant effect predictions reveals widespread adaptive evolution of human neurons

**DOI:** 10.1101/2025.09.11.675696

**Authors:** Alexander L. Starr, Michael E. Palmer, Jian Gao, Colin G. Nichols, Hunter B. Fraser

**Affiliations:** Department of Biology, Stanford University; Department of Cell Biology & Physiology, Washington University of St Louis School of Medicine

## Abstract

Detecting positive selection is essential to understanding evolution. Many methods to detect positive selection use simple classifications of genetic variants (e.g. synonymous/nonsynonymous). Here, we propose that these methods can be considered special cases of a more general framework to detect positive selection with any variant effect prediction method. Using evolutionary conservation and deep learning-based variant effect predictions, we apply this framework genome-wide in the human lineage and identify positive selection on protein sequences of genes involved in brain and heart development, chromatin accessibility in binding sites of dozens of transcription factors, and non-coding substitutions in compact genomic regions with reinforcing cell type-specific effects on *cis*-regulatory activity. Consistently, the dominant theme was positive selection on genes regulating neuronal connectivity, suggesting that fine-scale changes in brain wiring were essential to the evolution of human cognition. Overall, this framework represents a powerful, versatile tool to investigate adaptive evolution across the tree of life.

## Introduction

Over the course of the last few million years, humans have evolved a diverse array of unique traits not seen in other primates^1^. However, with these unique traits has come a large number of human-specific diseases. Common diseases including epithelial cancer, atherosclerosis, autism spectrum disorder (ASD), schizophrenia (SCZ), preeclampsia, Parkinson’s disease, and Alzheimer’s disease (AD) are far more prevalent in humans than in other apes, including our closest living relatives, chimpanzees (chimps)^1–7^. Despite intense interest in the genetic basis of these traits, we know very little about the genetic changes underlying uniquely human traits and susceptibility to these conditions. The core difficulty in this problem is three-fold. First, there are millions of genetic differences between humans and chimps, and it is likely that only a moderate fraction of these changes affect phenotype^8,9^. Second, although gene duplications and nonsynonymous substitutions (i.e., single nucleotide changes that change the amino acid sequence of a protein) likely played an important role in human evolution^10–15^, myriad data provide evidence for a more important role for non-coding mutations^16–20^, the effects of which are generally far more challenging to predict. Finally, it is likely that uniquely human traits are quite complex and highly polygenic, being shaped by the collective action (and interaction) of thousands of human-chimp differences affecting thousands of genes^16,21^.

The sheer number of genetic differences between humans and other apes necessitates the use of high-throughput methods to identify consequential genetic differences^8,20^. One approach to this problem is to predict the effects of human-chimp differences and focus on those with the strongest predicted effect. However, this is usually not sufficient to identify causal genetic differences. For example, there are over 25,000 human-derived substitutions that alter protein sequence, many of which likely have no detectable effect on phenotype^8^. As a result, variant effect predictions are often used to scan for signals of positive selection^20,22,23^ both because positively selected genetic variants must have had effects on organismal phenotypes (and so are prioritized for experimental follow-up) and because understanding the role of positive selection in shaping genetic variation is a key goal of evolutionary biology^22,24^.

Methods to detect positive selection generally fall into two categories. First, many methods, such as the McDonald-Kreitman (MK) test^22^, use a simple model to predict variant effect and binarily classify substitutions as either functional, meaning that the substitution is likely to affect biological function, or non-functional. For example, nonsynonymous substitutions in protein-coding regions could be considered functional and synonymous changes non-functional. One can then compare the number of fixed substitutions in each class to a null model of neutral evolution to detect positive selection^22,25–27^. This null model is often based on the number of polymorphic substitutions in each class: since polymorphic variants are unlikely to be selected for or against and therefore have nearly neutral effects^22^, an excess of fixed, functional substitutions provides evidence for positive selection^22,25^. Although this framework was initially primarily applied to protein-coding sequence^22,24^, it can also be applied to non-coding regions of the genome^25,28^. For example, genome-wide measurements of transcription factor binding enabled the development of INSIGHT, a method used to detect positive selection on transcription factor binding sites^25^. In addition, methods based on the MK test can be used to estimate the proportion of fixed substitutions that were fixed by positive selection, denoted α^24^, which has provided key insights into general principles governing protein evolution^10,24,29–31^.

The second class tests for differences in evolutionary rate in small genomic regions beyond what would be expected from neutral evolution^20,32^. The first of these methods has been fruitfully applied to identify Human Accelerated Regions (HARs), short primarily non-coding sequences that were highly conserved in mammals but have evolved rapidly in the human lineage^20,33,34^. In this case, sequence conservation is used to predict functionality. More recently, testing for rapid evolution in less conserved regions of the genome has enabled the identification of Human Ancestor Quickly Evolved Regions (HAQERs)^32^. Although variant effect predictions are not used to identify HAQERs, they were validated by contrasting the number of fixed and polymorphic sites in HAQERs relative to other regions of the genome^32^. Notably, the vast majority of HARs and HAQERs are non-coding and many have been shown to have divergent *cis*-regulatory properties from the ancestral sequences^32–36^.

Although most methods rely on the relatively simple variant classifications outlined above, recent advances in variant effect prediction could enable a much deeper understanding of positive selection on non-coding regions. For example, deep learning-based methods have made it possible to accurately predict the effects of substitutions on a variety of important molecular properties, including chromatin accessibility (CA)^37^, histone modifications^38^, 3D genome conformation^39,40^, transcription initiation^41^, transcription factor binding^42,43^, activity in reporter assays^44^, and RNA splicing^45^. As a result of this rapid progress, these new variant effect prediction methods should enable the development of powerful methods that leverage the diversity of variant effects to detect coding and non-coding polygenic positive selection and provide vital insight into evolution across the tree of life.

Here, we propose that methods which use binary variant effect predictions represent a small subset of a larger, much more general framework in which any variant effect prediction can be used to detect positive selection across the genome. By using this framework with conservation scores and predictions of variant effect on CA across 34 cell types, we discovered diverse signals of positive selection on the coding and non-coding portions of individual genes, sets of genes associated with human phenotypes, transcription factor binding sites, HARs, and HAQERs. Furthermore, this framework enables the comparison of the relative strength of positive selection across different genes, genomic regions, and variant effect predictions so can provide insight into the general principles governing evolution across the genome. In this vein, we use this framework to provide strong empirical support for the idea that Alu elements frequently experienced positive selection in the human lineage, the hypothesis that substitutions with cell type-specific effects were more likely to be fixed by positive selection, and that positive selection increased the specificity of cis-regulatory landscapes in developing neurons. Finally, we identify and validate an abundant class of small genomic elements with clusters of multiple reinforcing changes in cell type-specific CA that likely experienced positive selection in the human lineage.

### A framework to detect positive selection using variant effect predictions

A large body of work has shown that polymorphic sites with high minor allele frequency—known as common variants—are generally close to neutral (i.e., not positively or negatively selected)^22,24,29^. On the other hand, a subset of fixed substitutions generally have larger effects on fitness and were either positively selected or neutral since deleterious variants are unlikely to reach fixation (with some exceptions e.g., hitchhiking of slightly deleterious variants). Methods like the MK test combine this information with a binary variant effect prediction that classifies substitutions into neutral and non-neutral (or non-functional and functional), where the non-neutral sites are more likely to alter the function of some genomic element^22,25^. Therefore, if the fixed substitutions are enriched in the functional category, this rejects the null hypothesis of neutrality and instead implies that they are likely to have a larger impact on fitness than neutral variants. Since the variants were fixed, this impact on fitness must have generally been positive (since deleterious variants would typically not have fixed), implying the action of positive selection.

However, binary classifications fail to capture the full range of biological variation. For example, evolutionary conservation and effects on chromatin accessibility are continuous measures of predicted variant effect^20,37^. Using conservation as an example, the same logic outlined in the preceding paragraph holds true. If the distribution of conservation scores for fixed substitutions is significantly shifted toward higher scores than the polymorphic distribution, then we can reject the null hypothesis and conclude that there has been positive selection on that set of fixed substitutions (Fig. 1A). Alternatively, if the fixed and polymorphic distributions are not significantly different, then there is no evidence for positive selection (Fig. 1B).

**Figure 1:**
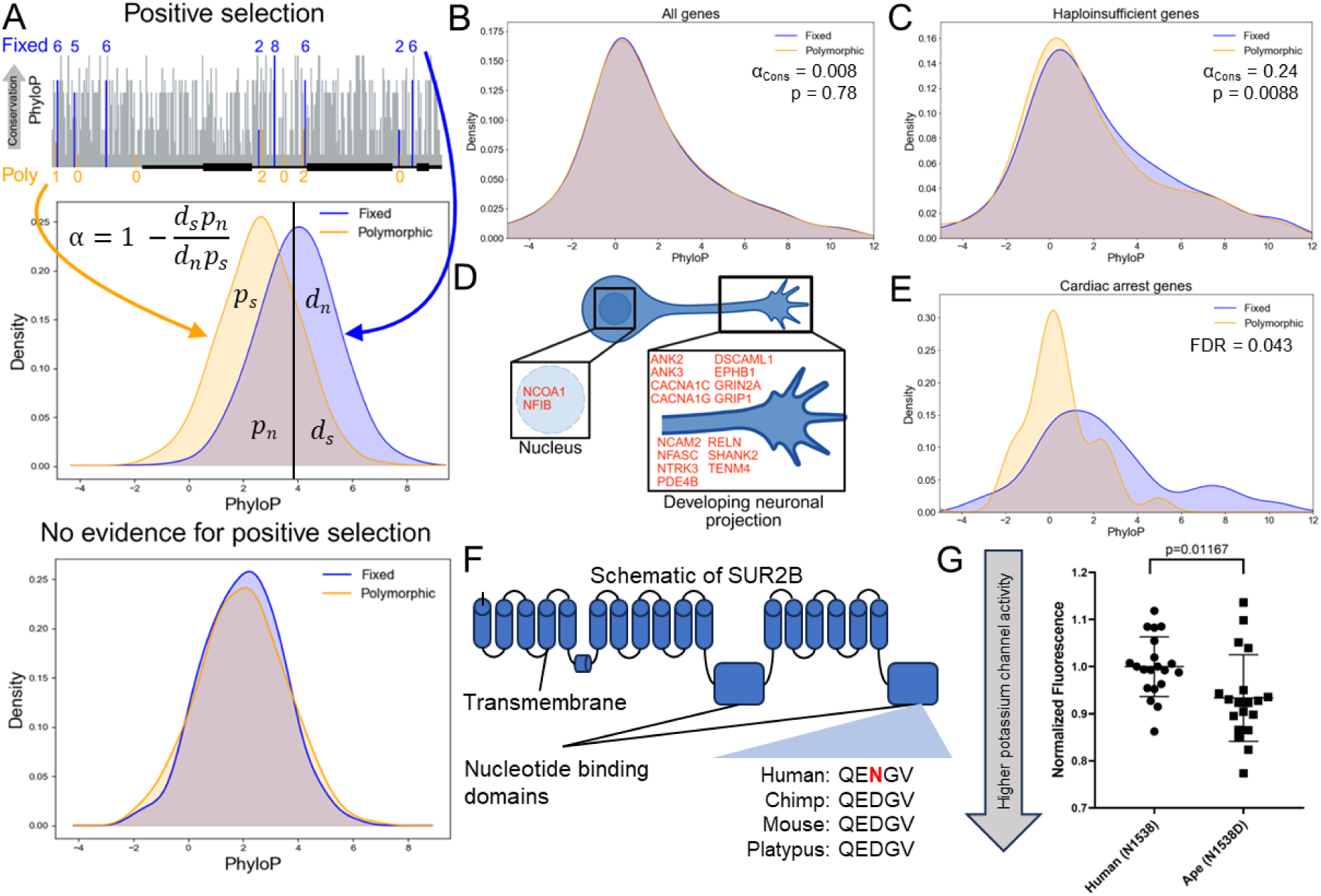
Positive selection on conserved non-synonymous sites. **A)** Outline of test for positive selection. **B)** The genome-wide distribution of conservation scores for fixed and polymorphic nonsynonymous substitutions are highly similar. In all cases, the cutoff for computing α_Cons_ is the 60^th^ percentile of the polymorphic conservation score distribution. **C)** Distribution of conservation scores for fixed and polymorphic substitutions in haploinsufficient genes. **D)** Schematic of developing neurons with proteins that contain a fixed human-derived nonsynonymous substitution in a site with PhyloP > 5 and have peak expression during mid-late fetal development in human cortical neurons. **E)** Distribution of conservation scores for fixed and polymorphic substitutions in genes linked to cardiac arrest. **F)** Schematic of SUR2B, which is encoded by *ABCC9*. The human-specific D1538N substitution is located in a nucleotide-binding domain at the C-terminal end of the SUR2B protein. **G)** Plot of normalized fluorescence, a readout of cell membrane potential (see Methods) affected by potassium channel conductance, comparing the human and chimpanzee SUR2B sequences. Data were analyzed with an unpaired t-test.

With the MK test, one can also estimate α, a lower bound on the proportion of nonsynonymous sites that were fixed by positive selection^24^. In direct analogy to this, we can generalize α to any variant effect prediction by specifying a cutoff to separate sites into putatively functional and non-functional^28^. For example, we can select a conservation score cutoff and classify sites as functional and non-functional to compute α_Cons_, a lower bound on the proportion of fixed substitutions meeting the conservation cutoff that were positively selected (Fig. 1A; throughout, we use the symbol α with a subscript that denotes what variant effect prediction was used to compute it. Although any variant of α is only well-defined between zero and one, negative α can result from polymorphic sites having greater functional effects than fixed sites^24,30^). We can then compare α_Cons_ across different sets of genomic regions and with different variant effect predictions to gain insight into the general principles governing which sites are positively selected, in a manner similar to how α derived from the MK test is used to quantify positive selection on proteins^31^. Moreover, this framework benefits from the deep understanding of the behavior of the MK test under various conditions^29,30,46,47^. For example, this framework can easily be adapted to an asymptotic version that corrects for mildly deleterious polymorphisms^30^.

### Positive selection on non-synonymous sites in genes regulating the development of neuronal connectivity and the heart

To apply this method to real data, we used gnomAD^48^, a repository of uniformly processed human whole genome sequencing data to identify polymorphisms and compute the derived allele frequency (DAF) using chimp and gorilla as outgroups (see Methods). To measure conservation, we computed PhyloP scores, a commonly used metric for conservation, using the Zoonomia 447-way placental mammal multi-way alignment^49,50^, masking the human genome. We then validated our method with simulations^51^, by recapitulating established results on positive selection in the human lineage^10,47^, and demonstrating weak effects of potential confounding factors such as background selection^21^ on our method (Supp. Text. 1-3).

With this dataset in hand, we first tested for positive selection on conserved nonsynonymous sites genome-wide using polymorphisms with high derived allele frequency (0.5 < DAF < 0.9) to remove most mildly deleterious polymorphisms. Consistent with previous work, we found no genome-wide evidence for positive selection on amino acid sequences in the human lineage using our test (Fig. 1B)^30^, with slightly (though not significantly) stronger evidence for positive selection using an asymptotic version of the test that corrects for segregating mildly deleterious sites (Supp. Fig. 1, α_Cons_ = 0.053, the confidence interval contains the previously estimated value of 0.13^30,52^).

We next tested whether haploinsufficient genes (defined as genes with probability of loss of function intolerance greater than 0.9^48^) collectively have stronger or weaker evidence for positive selection than genes that are not haploinsufficient, reasoning that conserved sites in these genes may evolve differently. Surprisingly, we found evidence for positive selection only in haploinsufficient genes using our framework (Fig. 1C, α_Cons_ = 0.24, p = 0.0088, Fisher’s exact test, with the “Cons” subscript denoting that α is computed from PhyloP scores). This effect was particularly pronounced among haploinsufficient genes that are depleted of rare nonsynonymous variants^48^ (Supp. Fig. 2A-C, α_Cons_ = 0.33, p = 0.0032 for nonsynonymous depleted; α_Cons_ = -0.2, p = 0.26 for all other haploinsufficient genes, Fisher’s exact test). In other words, one-third of changes in their most conserved amino acid sites were driven by positive selection, a rate 41-fold higher than the proteome as a whole (although this could be the result of increased constraint on these genes in the human lineage, see Supp. Text 3). This suggests that the most easily perturbed proteins in the genome—those sensitive to both a 50% reduction in dosage as well as a greater proportion of nonsynonymous changes—counterintuitively also show the strongest signals of adaptive evolution in the human lineage.

To better understand the biological consequences of this selective pressure, we tested whether nonsynonymous-depleted haploinsufficient genes were enriched for any Gene Ontology (GO) categories^53,54^. In addition to several broad categories, they were also enriched for more specific categories essential for neuronal signaling and development: ion channels and regulators of neuronal projections (axons, dendrites, and synapses) (Supp. Table 1). Positive selection on proteins that regulate the development of neuronal projections (e.g., EPHB1 and RELN, two proteins that regulate axonal development, Fig. 1D) is also supported by additional evidence (Supp. Text 4).

We next unbiasedly tested whether genes associated with any human phenotypes^55^ showed particularly strong evidence for positive selection relative to all genes, using a more relaxed DAF cutoff to enable more phenotypes to be tested. Genes associated with fetal cardiac arrest had considerably more highly conserved fixed sites than polymorphisms (Fig. 1E, FDR = 0.043 enrichment relative to genome-wide background, one-tailed Mann-Whitney U Test). In contrast, there was no evidence for positive selection using the MK test (odds ratio = 0.95, p = 0.89, Fisher’s exact test; see Supp. Text 5 for why this might be the case). Notably, human heart morphology is strikingly different from that of other great apes^56,57^ and human cardiac arrests are caused by very different underlying mechanisms^2^.

The most conserved fixed substitution in the cardiac arrest category lies in the ion channel subunit gene *ABCC9*, encoding the SUR2B isoform (Fig. 1F) which is primarily expressed in vascular smooth muscle and regulates heart, craniofacial, and brain development, hair growth, and vocal pitch^58–60^, all of which are strikingly diverged between humans and other great apes^1,61^. To test whether this fixed difference might have phenotypic effects, we tested the effects of reverting this amino acid to the ancestral residue on the activity of ATP-sensitive (K_ATP_) channels^62^. Human SUR2B-containing channels had decreased conductance relative to ancestral SUR2B-containing channels (Fig. 1G), confirming that this change led to functional divergence that could impact organismal phenotypes, especially considering the dosage sensitivity of *ABCC9*^58,59^. Collectively, these results suggest that positive selection on conserved nonsynonymous sites has shaped human nervous and cardiovascular development as well as demonstrate the utility of our approach when applied to the coding genome.

### Positive selection on conserved sites in the non-coding genome

Next, we focused on the non-coding genome and, first, further validating our method. Unlike nonsynonymous sites, many non-coding sites lie in evolutionarily newer regions of the genome. As PhyloP scores tend to be less accurate when the alignment only contains a small total branch length^49,50^ (i.e., the total divergence time across all species is relatively low), we removed non-coding sites with fewer than 250 species in the alignment (see Methods). Although much less is known about the evolution of non-coding sites, one consistent result is that there has been stronger positive selection on the X chromosome relative to autosomes in humans and other apes^63–65^. Consistent with this, the X chromosome had the strongest evidence for positive selection among all chromosomes (α_Cons_ = 0.038, p = 0.0008, Fisher’s exact test; Fig. 2A, Supp. Fig. 3A). To further validate our method, we tested whether previously identified hotspots of selection on the X^63,65^ had evidence for positive selection with our approach. Encouragingly, we found 6.2-fold more positive selection in these regions (Supp. Fig. 3B, α_Cons_ = 0.13, p = 0.0035) relative to the rest of the X.

**Figure 2:**
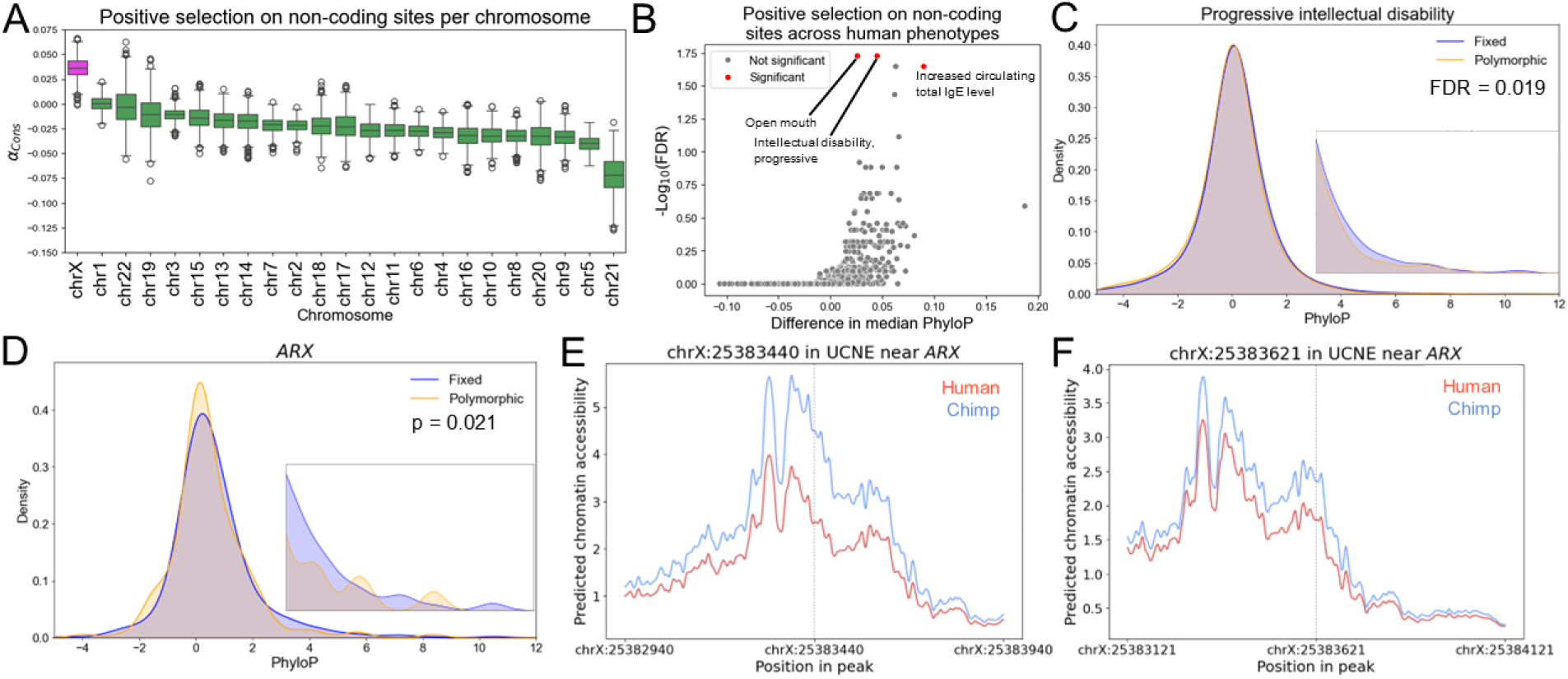
Positive selection on conserved non-coding sites in the human genome. **A)** 95% confidence interval for α_Cons_ on each chromosome based on 1,000 bootstraps. The cutoff for computing α_Cons_ is the 60^th^ percentile of the polymorphic conservation score distribution. Magenta color signifies that the 95% confidence interval does not contain 0. **B)** Positive selection on conserved non-coding sites across human phenotypes. Gene sets were considered significant if they had Mann-Whitney U test FDR < 0.05 and Fisher’s exact test nominal p-value < 0.05, with the cutoff being the 90^th^ percentile of the polymorphic conservation score distribution. **C)** Distribution of PhyloP scores for fixed and polymorphic sites near genes related to progressive intellectual disability. The inset shows the distribution magnified in the range 3-12. **D)** Same as in (C) but for fixed and polymorphic sites near *ARX*. **E)** Predicted effect of a human-specific substitution in an ultraconserved non-coding element (UCNE) on chromatin accessibility in fetal cortical progenitors. **F)** Same as in (E) but for a different substitution in the same UCNE.

However, we also found positive selection on the rest of the X chromosome suggesting positive selection outside of these known hotspots as well (Supp. Fig. 3C, α_Cons_ = 0.021, p = 0.041). To explore phenotypes that might be affected by positive selection on the X chromosome, we tested whether variants near genes associated with human phenotypes^55^ have evidence for positive selection on nearby conserved sites. The only significant phenotypes were related to toe development, e.g., genes whose loss of function leads to short toes (FDR = 0.00024, one-tailed Mann-Whitney U test, Supp. Fig. 3D-E). This was partially driven by sites near *GPC4* (Supp. Fig. 3F), a gene involved in brain and skeletal development. Overall, these findings support the idea that there has been stronger positive selection on the human X chromosome and that this may have shaped human skeletal development.

Encouraged by this, we tested for positive selection on non-coding sites genome-wide across all genes. However, we found no evidence for positive selection on conserved non-coding sites in the human genome with or without the asymptotic correction (Supp. Fig. 4A-B, α_Cons_ = -0.022, p = 9.8×10^-5^). Interestingly, we did find evidence for positive selection on conserved non-coding sites in wild house mice^66^ (*Mus musculus domesticus*, Supp. Fig. 5A-B, α_Cons_ = 0.040, p = 1.1×10^-19^, Fisher’s exact test) and the black flying fox^67^ (*Pteropus alecto*, Supp. Fig. 5C-D, α_Cons_ = 0.044, p = 2.7×10^-26^, Fisher’s exact test), though not for human using only three-way one-to-one orthologs (Supp. Fig. 5E-F, α_Cons_ = -0.014, p = 1.1×10^-19^, Fisher’s exact test) suggesting that our method has sufficient power to detect positive selection in non-coding sites of mammalian genomes. This is consistent with previous work that found stronger evidence for positive selection on nonsynonymous substitutions in house mice than humans^68,69^ and could be the result of a larger proportion of non-coding polymorphisms being mildly deleterious in humans relative to house mice and black flying foxes^69^.

We next tested for positive selection on conserved non-coding sites near genes associated with particular human phenotypes^55^. Three gene sets met our criteria for statistical significance (Fig. 2B, Methods) including Increased circulating total IgE level (Supp. Fig. 6A) and progressive intellectual disability (Fig. 2D). For example, one gene contributing to the total IgE level signal was *PLA2G7*, which is also thought to be an important modulator of lifespan^70^. Consistent with its signal of non-coding positive selection, *PLA2G7* also has lower *cis*-regulatory activity in humans compared to chimps across many different cell types^71^ (Supp. Fig. 6B).

One of the main drivers of the signal for intellectual disability was the transcription factor *ARX* (Fig. 2E). Interestingly, the three most conserved fixed substitutions (in sites chrX:24990968, chrX:25383440, and chrX:25001488) near *ARX* lie in experimentally validated enhancers^72,73^ active in the developing brain, with the first two listed located very near to ultraconserved elements^74,75^. One of these, at position chrX:25383440, lies in a CRE that primarily drives expression in forebrain neural progenitor cells^73^ and is predicted to decrease chromatin accessibility in forebrain neural progenitors (Fig. 2F, log_2_ fold-change = -0.5, see below for how predicted effects were generated). In addition, another nearby substitution in a conserved site is also predicted to decrease accessibility (Fig. 2G, log_2_ fold-change = -0.28). Given the extreme conservation of this validated CRE as well as the individual sites with substitutions, and large predicted effects on accessibility the divergence in the activity of this CRE, it may have had important phenotypic effects on human brain development.

### Testing for positive selection in the non-coding genome using machine learning-based variant effect predictions

Although conservation is a very useful predictor of variant effect, we reasoned that other variant effect predictions, such as those derived from machine learning models, would enable new insights into human evolution. As it is still difficult to predict the effect of variants on gene expression, we used ChromBPNet^37^ to predict the effects of variants on chromatin accessibility (CA), an important component of transcriptional *cis*-regulation that can also capture variant effects on transcription factor (TF) binding. The use of ChromBPNet for variant effect prediction has been extensively validated using a variety of experimental methods, including *in vitro* massively parallel reporter assays and *in vivo* reporter assays^37,76^. We trained ChromBPNet models on ATAC-seq data from 34 pure populations of human cell types from a variety of organs, body systems, and developmental stages^77–86^ (see Methods for a list of cell types and model validation) and predicted the effects of all fixed human-chimp differences as well as common human polymorphisms (DAF > 0.1) as input to our framework (Fig. 3A).

**Figure 3:**
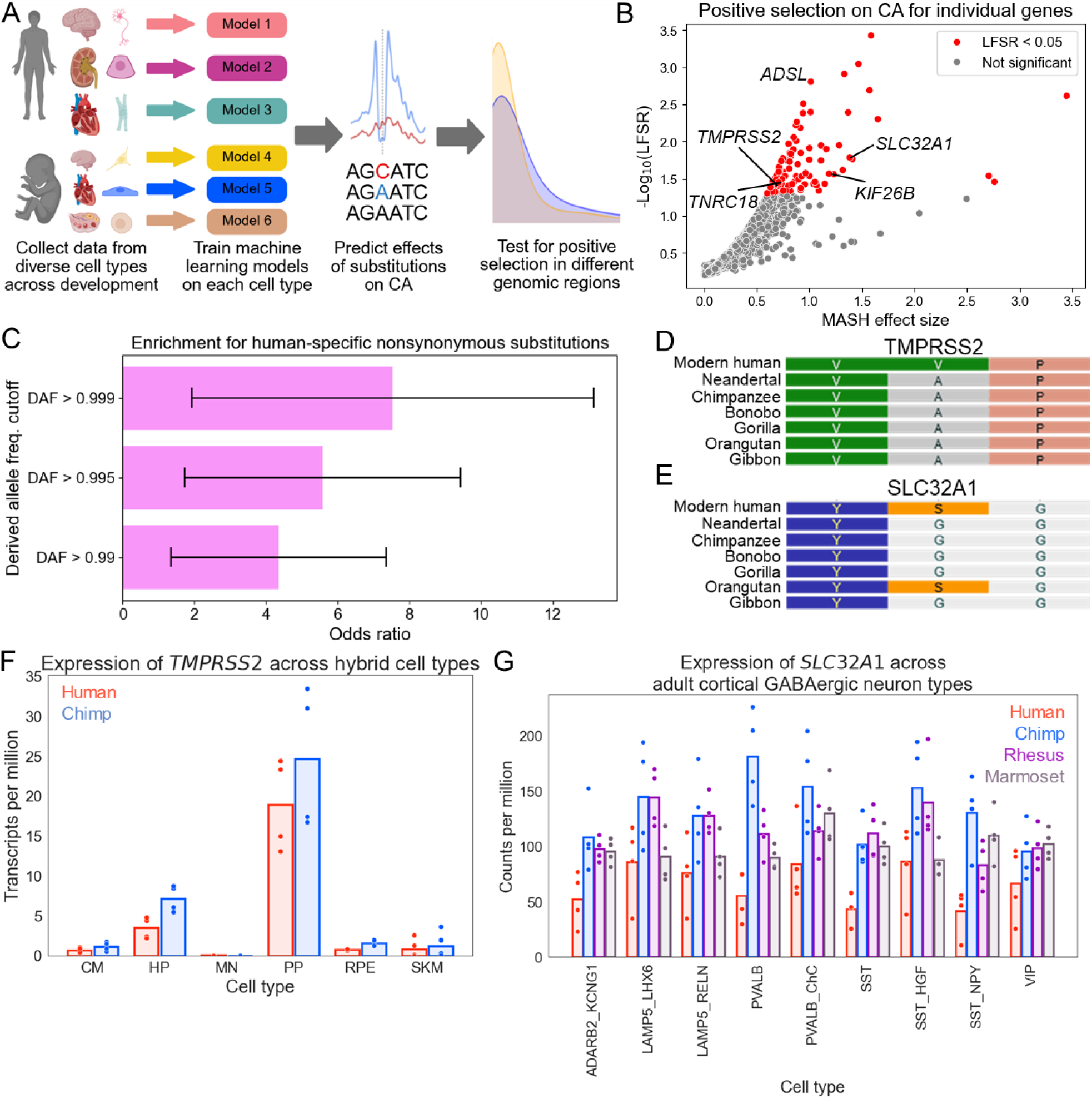
Detecting positive selection using machine learning-based predictions of variant effects. **A)** Outline of strategy to detect positive selection using predictions of variant effects on CA. **B)** Volcano plot of positive selection on CA per gene. Only genes with MASH effect size greater than zero are shown. The maximum effect size for each gene across cell types and corresponding local false sign rate are shown on the x- and y-axes respectively. **C)** Odds ratios and confidence interval for Fisher’s exact test for enrichment of genes with nearly fixed, human-derived, modern human/Neanderthal nonsynonymous substitutions. All three cutoffs yield p < 0.01 **D)** Sequence alignment of TMPRSS2 residues 32-34. **E)** Sequence alignment of SLC32A1 (also known as VGAT) residues 422-424. **F)** Allele-specific expression of *TMPRSS2* across six hybrid cell types (CM = cardiomyocytes, HP = hepatic progenitors, MN = motor neurons, PP = pancreatic progenitors, RPE = retinal pigmented epithelial cells, SKM = skeletal muscle). Chimp alleles are more highly expressed at FDR < 0.05 in CM, HP, PP, and RPE. **G)** Expression of *SLC32A1* across adult cortical GABAergic neuron types in the DLPFC. Cell types are denoted by their marker genes as in the original study. Chimpanzee expression is significantly greater than human (FDR < 0.05) in ADARB2_KCNG1, PVALB, SST, and ADARB2_KCNG1.

As an initial application of this approach, we tested for positive selection on changes in CA per gene, genome-wide in each cell type, using MASH^87^ to combine p-values across cell types. We identified 105 genes with evidence for selection in at least one cell type (Fig. 3B, MASH local false sign rate or LFSR < 0.05 in at least one cell type). We noticed that several of these genes are known to contain nearly fixed human-derived human-neanderthal nonsynonymous substitutions, including one in *ADSL*^88^ that has previously been shown to alter its activity (Fig. 3B). In line with this, genes with evidence for positive selection on CA since the human-chimp split are enriched for nearly fixed human-derived human-neanderthal nonsynonymous genetic differences and this enrichment gets stronger with more stringent DAF cutoffs (Fig. 3C, odds ratio = 5.57, p = 0.0029, Fisher’s exact test). This congruence was unexpected as it bridges not only time scales (human/chimp vs. human/neanderthal divergence), but also fundamentally different targets of selection (amino acid sequences vs. CA).

For example, TMPRSS2, which plays a key role in SARS-CoV-2 pathogenesis^89^, has an alanine at residue 33 in Neanderthals, Denisovans, as well as non-human great apes but a valine in the vast majority of humans^90^ (Fig. 3D). As another example, SLC32A1 (also known as VGAT), which is a marker for GABAergic neurons and mediates GABA and glycine uptake into synaptic vesicles^91^, has a serine as residue 423 but a glycine in Neanderthals^90^ as well as most other apes (Fig. 3E). Interestingly, *TMPRSS2* is differentially expressed between the human and chimp alleles in multiple different human-chimp hybrid iPSC-derived cell types^71^ (Fig. 3F) and *SLC32A1* has lower expression in humans relative to chimps, rhesus macaques, and marmosets in several adult cortical GABAergic neuron types^92^ (Fig. 3G). Overall, this suggests that genes whose protein sequence changes likely contributed to human-Neanderthal divergence have also been under selection for human-specific *cis*-regulation.

### Cell type-specific positive selection on CA in HARs and HAQERs

We next applied our method to human accelerated regions (HARs)^93^ and human ancestor quickly evolved regions (HAQERs)^32^, two sets of regions thought to have undergone positive selection in the human lineage. Notably and unlike most other methods to detect positive selection, comparing the distributions of predicted variant effects is orthogonal to methods based solely on evolutionary rate, enabling us to provide new insight into the selective forces shaping these well-studied regions. Furthermore, testing for positive selection on CA in different cell types can help pinpoint the cell types in which positive selection on these elements was most important. In general, we observe α_CA_ (i.e., α estimated using predicted effect on total chromatin accessibility) greater than 0 across most cell types in both regions (Fig. 4A-B), although this only reaches nominal significance in a few cell types for HARs (fetal cortical excitatory neurons and fetal chondrocytes) and HAQERs (adult ventricular cardiomyocytes, adult microglia, fetal cortical excitatory neurons, and fetal cortical neural progenitors), suggesting that HARs and HAQERs may play particularly important roles in these cell types (Fig. 4A-B). Consistent with this, previous work has suggested that both HARs and HAQERs are associated with brain development^20,32,34,36^ and that HARs (but not HAQERs) are associated with the skeletal system^94^. However, the association of HAQERs with the heart and immune system is previously unreported. For example, HAQERs near *NRROS* and *METTL4* contain multiple fixed substitutions predicted to increase accessibility in cardiomyocytes and both genes have higher expression from the human allele in human-chimp hybrid cardiomyocytes (Supp. Fig. 7-8).

**Figure 4:**
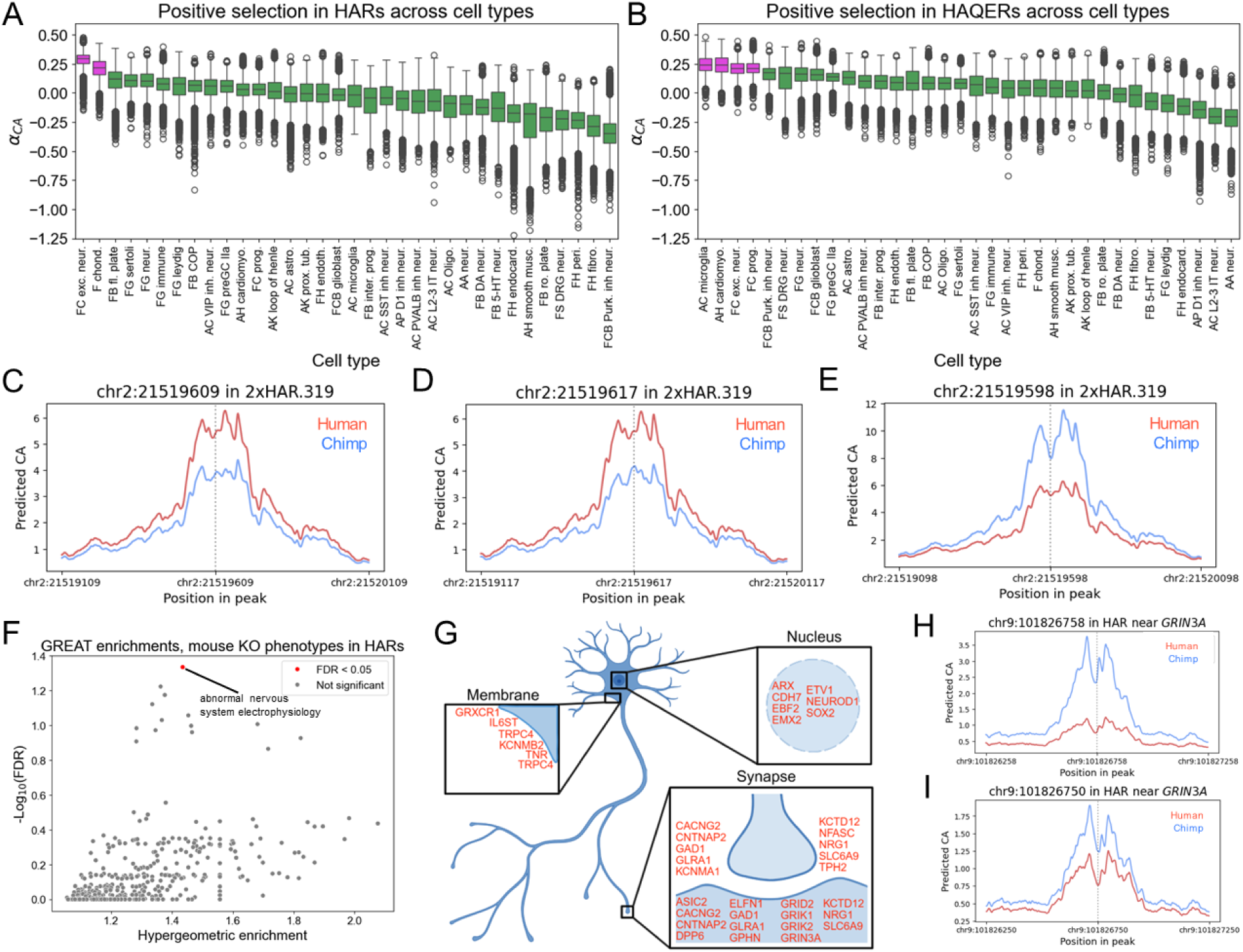
Positive selection on chromatin accessibility in HARs and HAQERs. **A)** Box plot for HARs in each cell type based on 1,000 bootstraps. The cutoff for computing α_CA_ is the 90^th^ percentile of the polymorphic conservation score distribution. Magenta color signifies that the 95% confidence interval does not contain 0. **B)** Same as in (A) but for HAQERs. **C-E)** Predicted effects of three fixed substitutions in 2xHAR.319 on chromatin accessibility. **F)** Volcano plot of GREAT enrichment for high predicted log_2_ fold-change in CA for mouse single gene KO phenotype categories. The fold-enrichment for each gene and corresponding FDR are shown on the x- and y-axes respectively. **G)** Schematic of a neuron with genes contributing to the enrichment in (F) labeled. Genes are placed in the cellular compartment (membrane, nucleus, presynapse at the top of the synapse inset, or postsynapse at the bottom of the synapse inset). **H-I)** Predicted effects of two fixed substitutions in a HAR near *GRIN3A* on chromatin accessibility in fetal cortical excitatory neurons. The smoothed predicted counts per base are shown.

Although the accuracy of machine learning predictions in predicting variant effects in HARs is well-established^35^, we sought to explore the effects of experimental repression (by CRISPR interference) of a HAR with substitutions having large predicted effects. In fetal cortical neurons, the HAR with the largest effect substitutions was 2xHAR.319. Recently published CRISPRi in this same cell type showed that the human allele increases the expression of the nearby gene *PUM2*, while decreasing the expression of another gene, *LAPTM4A*^36^. Four out of five substitutions in the HAR are predicted to increase CA in the human lineage (Fig. 4C-E, Supp. Fig. 9A-B, all log_2_ fold-changes > 0.24) and one substitution is predicted to decrease CA (Fig. 4E, log_2_ fold-change = -0.64). Overall, this suggests that some of the many HARs that appear to be undergoing compensatory evolution can still be highly divergent between humans and chimps and have phenotypic consequences.

As there are an insufficient number of polymorphic sites in HARs to further subdivide them, we adopted a different strategy to test whether particular sets of genes might have been affected by selection on CA in HARs. We tested whether fixed sites with large predicted effects on CA were enriched near gene sets associated with particular phenotypes or functions relative to fixed sites with very weak effects on CA using GREAT^95^, validating significant results with a permutation procedure. Focusing on HARs in fetal cortical excitatory neurons, we found that large effect sites were associated with several broad gene sets such as “Post-translational modification” (Supp. Fig. 10A-B, FDR = 7.2×10^-7^). However, they were also enriched near gene sets that imply more specific functions such as “Abnormal nervous system electrophysiology” (Supp. Fig. 10A-B, Fig. 4F FDR = 0.046), most of which either regulate transcription in the nucleus or localize to the synapse (Fig. 4G). For example, two fixed substitutions that are predicted to decrease accessibility in fetal excitatory neurons collectively repress a HAR near *GRIN3A* (Fig. 4H-I), an ion channel subunit that plays a key role in cortical computation^96^. Overall, the combination of machine learning with our new framework enables new insights into well-studied rapidly evolving regions of the human genome.

### Positive selection on chromatin accessibility of transcription factor binding sites

Much of *cis*-regulatory evolution is thought to occur through substitutions in TF binding sites (TFBS). Although several methods have been developed and fruitfully applied to detect selection on the binding sites of particular TFs^25^, these methods usually rely on comparisons to putatively neutral sites not in TFBS. However, sites outside of known TFBS frequently also have *cis*-regulatory effects^97–101^, introducing potential confounders similar to those present for the MK test due to selection on synonymous sites. Moreover, these methods do not nominate specific cell types in which positive selection has been focused. We reasoned that applying our framework when restricting to predicted TFBS would provide a powerful new way to detect selection on the binding sites of particular TFs.

Applying this idea to the predicted binding sites of 633 TFs^102^ across 34 cell types and combining information with MASH^87^, we detected positive selection on the binding sites of 37 TFs (MASH LFSR < 0.05, Supp. Fig. 11). Many TFs have evidence for positive selection on the CA of their binding sites in only one or a few cell types. For example, there is only evidence for positive selection on the CA of RXRA binding sites in adult kidney proximal tubule cells (Fig. 5A-B). One substitution in an RXRA binding site near the TSS of another TF that plays an important role in kidney function, *PAX2*^103^, is predicted to approximately double the accessibility of a candidate *cis*-regulatory element (Fig. 5C). As another example, we only find evidence for positive selection on CA of MEF2A binding sites in a few cell types with the strongest evidence coming from adult cortical excitatory neurons (Supp. Fig. 12A-B). Intriguingly, large-effect fixed sites in predicted MEF2A binding sites are enriched near regulators of synapse assembly in cortical excitatory neurons (Supp. Fig. 12C, FDR = 0.033) This is particularly interesting as MEF2A has been previously proposed to underlie the prolonged period of synaptogenesis specific to human cortical neuron development^104^.

**Figure 5:**
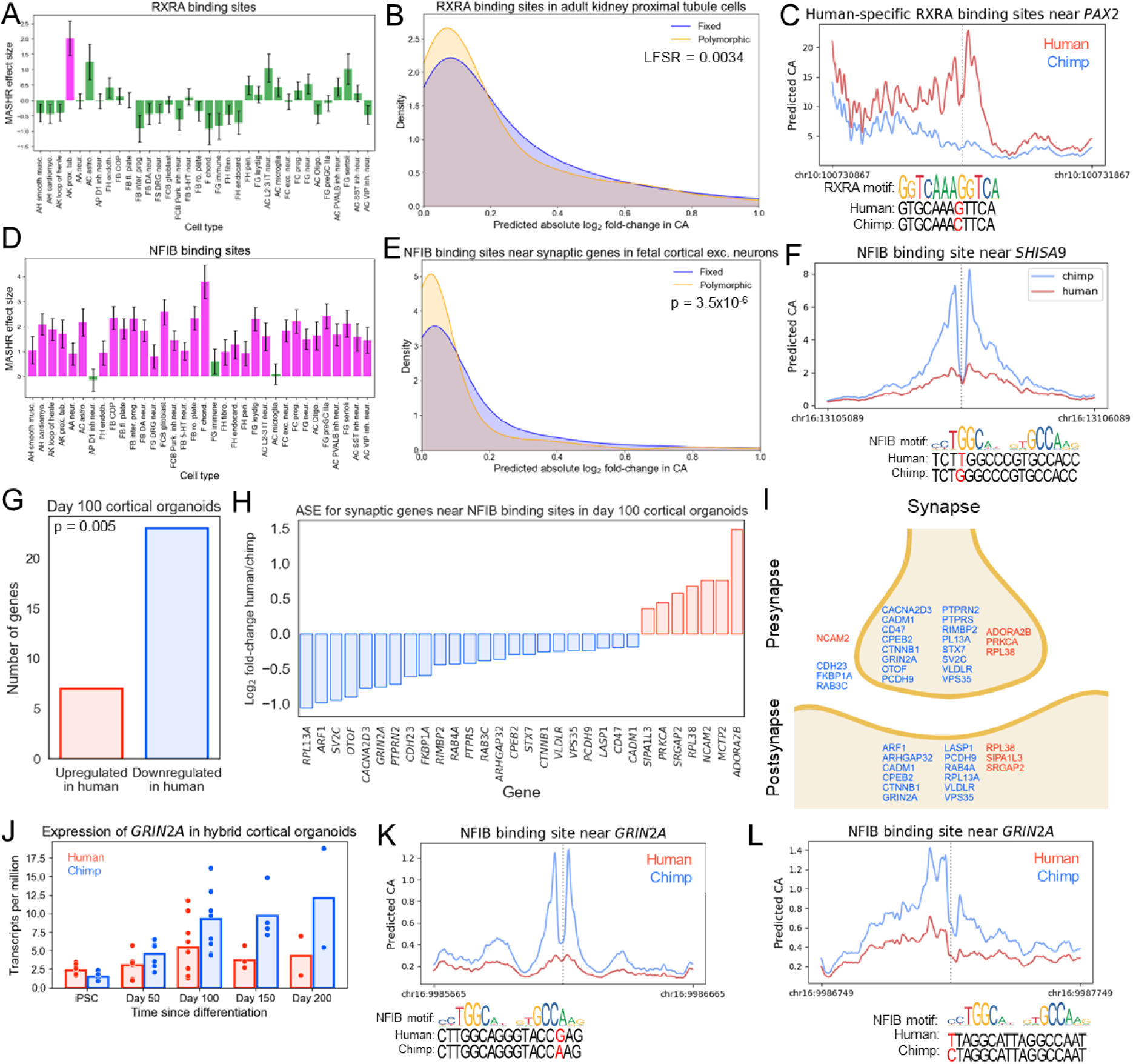
Positive selection on chromatin accessibility in transcription factor binding sites. **A)** Positive selection on RXRA binding sites across cell types. Bar height shows the estimated effect size from MASHR, the error bar is the posterior standard deviation from MASHR. Magenta indicates local false sign rate (LFSR) < 0.05. In all cases, the cutoff for Fisher’s exact test is the 60^th^ percentile of the polymorphic predicted log_2_ fold-change in CA distribution. **B)** Distribution of predicted absolute log_2_ fold-change in CA for fixed and polymorphic substitutions in RXRA binding sites in adult kidney proximal tubule cells. **C)** Predicted effect of fixed substitution at chr10:100731367 in a candidate CRE near *PAX2* on chromatin accessibility in adult kidney proximal tubule cells. The smoothed predicted chromatin accessibility is shown. The RXRA binding motif is shown below the plot along with the human and chimp sequence context for chr10:100731367. **D)** Same as in (A) but for NFIB binding sites. **E)** Distribution of predicted absolute log_2_ fold-change in CA for fixed and polymorphic substitutions in NFIB binding sites near synaptic genes in fetal cortical excitatory neurons. **F)** Predicted effect of fixed substitution at chr16:13105589 near *SHISA9* on chromatin accessibility in fetal cortical excitatory neurons. The smoothed predicted chromatin accessibility is shown. The NFIB binding motif is shown below the plot along with the human and chimp sequence context for chr16:13105589. **G)** Barplot showing number of putative NFIB target genes with increased (red) and decreased (blue) expression from the human allele in day 100 human-chimp hybrid cortical organoids. **H)** Differential expression of individual genes from (G). All genes have FDR < 0.05. **I)** Schematic of a synapse with genes from (H) labeled. Blue indicates lower expression from the human allele, red higher expression. Genes are placed in the synaptic compartment (presynapse at the top, postsynapse at the bottom, unknown localization outside the colored area) to which their encoded proteins localize. **J)** Expression of *GRIN2A* from the human and chimp alleles across a time course of human-chimp hybrid cortical organoid development. **K)** Predicted effect of fixed substitution at chr16:9986165 in a candidate CRE near *GRIN2A* on chromatin accessibility in fetal cortical excitatory neurons. The smoothed predicted counts per base are shown. The NFIB binding motif is shown below the plot along with the human and chimp sequence context for chr16:9986165. **L)** Same as in (K) but for chr16:9987249.

In contrast, several TFs have evidence for positive selection across many different cell types. For example, there is evidence for positive selection on the binding sites of ZNF460 and ZNF135 in most non-neuronal cell types and especially fetal chondrocytes (Supp. Fig. 13A-C). Interestingly, the predicted binding sites of both TFs overwhelmingly lie in Alu elements (e.g., 85% of fixed substitutions in ZNF460 binding sites are in Alu elements), consistent with previous work^105^ and suggesting that these TFs may play an important role in regulating Alu elements. Furthermore, this signal of positive selection is largely specific to ZNF460 binding sites in Alu elements (Supp. Fig. 13D-E, α_CA_ = 0.066 for sites in Alu elements, compared to α_CA_ = 0.019 for sites not in Alu elements, using fetal chondrocyte predictions). When expanding to all fixed and polymorphic sites within Alu elements, we find evidence for stronger positive selection than in non-Alu elements in 26 out of 34 cell types (Supp. Fig. 13F). Importantly, previous work has experimentally validated ZNF460 and ZNF135 binding to Alu elements and that Alu elements are enriched for differences in *cis*-regulatory activity between humans and macaques^105^. Collectively, this provides strong empirical support for the longstanding hypothesis that Alus provided an important source of adaptive *cis*-regulatory divergence in the human lineage^106,107^.

As a final example, we found evidence for positive selection on the binding sites of NFIB, a broadly expressed TF that plays a key role in the development of the brain and other organs^108–110^ across many cell types (Fig. 5D). Due to its known role in brain development^108^, we focus on NFIB binding sites in fetal cortical neurons (Supp. Fig. 14A). Interestingly, large effect fixed substitutions are enriched for a variety of synaptic terms in fetal cortical excitatory neurons (e.g., postsynaptic membrane and presynapse, FDR = 0.033 for both gene sets, Supp. Fig. 14B). Based on this, we restricted to known synaptic genes^111^ and compared the predicted effect size of fixed and polymorphic sites in fetal cortical excitatory neurons. Strikingly, we observed strong evidence for positive selection on synaptic genes (Fig. 5E) and little evidence for positive selection on non-synaptic genes (Supp. Fig. 14C), suggesting that the positive selection on NFIB binding sites primarily affects the expression of synaptic genes. For example, there is a human-specific substitution in a highly conserved site near *SHISA9*, an ASD-associated synaptic gene^112^, that disrupts an NFIB binding site leading to a decrease in CA (Fig. 5F, predicted log_2_ fold-change in CA = -0.95 To further explore this, we focused on synaptic genes near fixed substitutions with large predicted effect sizes with sign matching that of the predicted change in NFIB binding and differences in expression of the human and chimp alleles in human-chimp hybrid cortical organoids (which express NFIB, Supp. Fig 15A). Interestingly, there is a significant bias toward decreased expression of these genes from the human allele (Fig. 5G-I) suggesting that there has been positive selection for downregulation of these genes^113^ and supporting the idea that there has been positive selection on NFIB binding sites. For example, *GRIN2A*, an NMDA receptor subunit that is associated with a variety of neurodevelopmental outcomes including speech disruption^114,115^, has lower expression from the human allele in human-chimp hybrid cortical organoids^116^ (Fig. 5J) and two fixed substitutions near *GRIN2A* are predicted to reduce NFIB binding and CA (Fig. 5K-L). Similarly, *VLDLR*, a regulator of dendritic spine formation, has lower expression from the human allele^117^ (Supp. Fig. 15B) and a nearby fixed substitution is predicted to eliminate an NFIB binding site and decrease accessibility of the region (Supp. Fig 15C). Overall, this suggests that positive selection on NFIB binding sites has shaped the development and function of neuronal connectivity in the human lineage.

### Variants with cell type-specific effects on chromatin accessibility are more likely to be positively selected than those with broader effects

A long-standing theory is that variants with more context-specific effects are more likely to be fixed by positive selection^71,118,119^, primarily because they are less pleiotropic and so less likely to be subject to antagonistic pleiotropy^118,119^. This difference in pleiotropy is also often thought to be one of the primary reasons that positive selection more often acts on non-coding substitutions^118,119^. Despite the staying power of this theory, evidence for it is indirect^71,120^, (particularly for *cis*-regulatory evolution and in mammals), leaving it an open question whether this theory holds true in practice. We reasoned that we could use our framework to explicitly test whether variants with more context-specific effects on CA are more likely to be fixed by positive selection.

To do this, we focused on cell types with maximally diverse *trans*-regulatory environments by excluding similar cell types (e.g., we only included one type of fetal neuron), leaving us with two groups of eight cell types to ensure that any results replicate across different groups of cell types. We then computed dTau^71^, a measure of the cell type-specificity of the predicted effect of a variant on cell type-specificity, for each variant in each of the two groups of cell types. This metric ranges from 0 (if the predicted effect is identical in all cell types) to 1 (if the predicted effect is maximally different between one cell type and all others). For each cell type, we then restricted to variants with moderate predicted effect (the 10^6^ sites with the largest absolute log fold-changes, ∼8% of sites) and compared dTau for fixed and polymorphic sites.

Consistent with the idea that variants with more cell type-specific effects are more likely to be positively selected, the fixed substitutions generally had higher dTau than polymorphisms (e.g., for fetal cortical excitatory neurons in Fig. 6A, α_CTS_ = 0.069 where CTS stands for cell type-specific, p = 1.3×10^-42^, Fisher’s exact test). In principle, this could be driven by higher predicted log_2_ fold-change in CA in the top 8% of fixed sites relative to polymorphisms, independent of the cell type-specificity of those effects. However, we found α_CTS_ to be significantly higher than α_CA_ (Fig. 6B, α_CA_ = 0.014, p = 0.0032, Fisher’s exact test), suggesting that this does not explain the results for α_CTS_. Consistent with this, in both groups and across most cell types, we found high α_CTS_ absolutely and relative to α_CA_ (Fig. 6C, Supp. Fig. 16). To confirm this, we stratified variants by dTau and ran the test using predicted effect on CA in each bin. We consistently observed stronger evidence for positive selection in more cell type-specific bins (Supp. Fig. 17A) and this was robust to the removal of several cell types (Supp. Fig. 17B-F). Collectively, these results provide support for the longstanding theory that variants with more context-specific effects are more likely to be fixed by positive selection.

**Figure 6:**
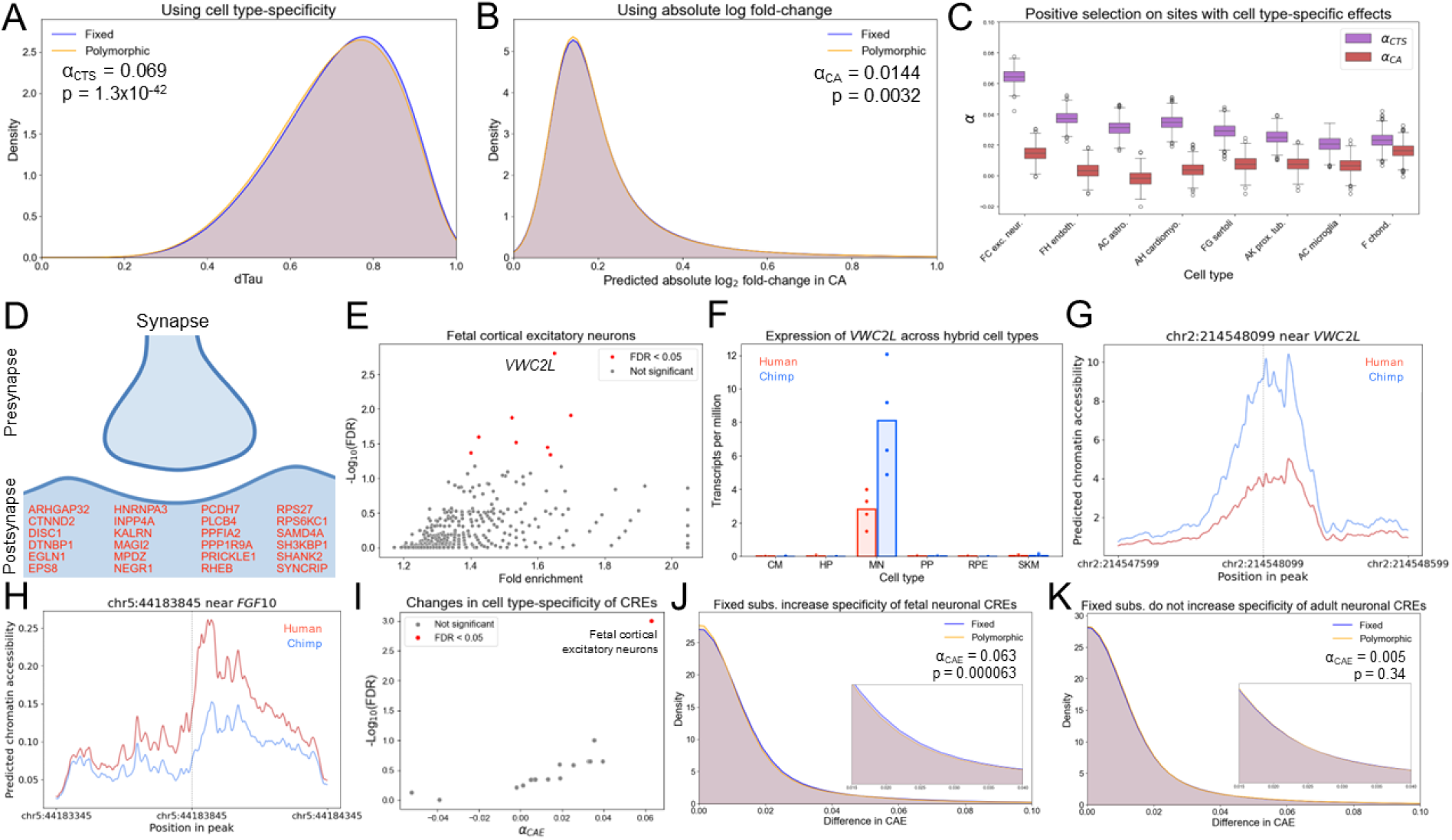
Positive selection on substitutions with cell type-specific effects. **A)** Distribution of cell type-specificity of predicted effects on CA for fixed and polymorphic substitutions with absolute log_2_ fold-change > 0.1 in fetal cortical excitatory neurons. The cutoff for computing α_CTS_ is the 30^th^ percentile of the polymorphic absolute log_2_ fold-change distribution. **B)** Distribution of predicted effects on CA for fixed and polymorphic substitutions with absolute log_2_ fold-change > 0.1 in fetal cortical excitatory neurons. The cutoff for computing α_CA_ is the 60^th^ percentile of the polymorphic absolute log_2_ fold-change distribution. **C)** Box plot comparing α_CTS_ and α_CA_ across cell types based on 1,000 bootstraps. The cutoff for computing α is the 60^th^ percentile of the polymorphic score distribution. **D)** List of genes that localize to the postsynaptic density and have at least one fixed site with dTau > 0.95 and predicted absolute log_2_ fold-change in CA > 0.5. **E)** Volcano plot of GREAT enrichment for high dTau per gene. The fold enrichment for each gene and corresponding local false sign rate are shown on the x- and y-axes respectively. **F)** Expression of *VWC2L* across six hybrid cell types (CM = cardiomyocytes, HP = hepatic progenitors, MN = motor neurons, PP = pancreatic progenitors, RPE = retinal pigmented epithelial cells, SKM = skeletal muscle). FDR < 0.05 in MN. **G)** Predicted effect of a fixed substitution at chr2:214548099 in a candidate CRE near *VWC2L* on chromatin accessibility in fetal cortical excitatory neurons. **H)** Same as in (G) but for chr5:44183845 near *FGF10* in pre-granulosa cells. **I)** Volcano plot showing evidence for positive selection on substitutions that increase cell type-specificity of candidate CREs. In all cases, the cutoff for computing α_CAE_ is the 90^th^ percentile of the polymorphic conservation score distribution. **J)** Distribution of differences in chromatin accessibility enrichment (CAE) for fixed and polymorphic substitutions in fetal cortical excitatory neurons. The inset shows the distribution in the range 0.014-0.04. **K)** Same as in (J) but for adult L2/3 intratelencephalic neurons.

Next, we explored which genes might be most affected by the stronger positive selection on variants with cell type-specific effects, focusing on fetal cortical excitatory neurons and fetal pre-granulosa cells (which play a key role in oocyte development) as these cell types have the strongest evidence for positive selection on variants with cell type-specific effects. To do this, we used GREAT to test whether fixed substitutions with large cell type-specific effects were enriched near genes or gene sets relative to all substitutions. Interestingly, we observed strong enrichment near genes related to the synapse (the top terms for Cellular Component are postsynapse and synaptic membrane, FDR < 0.0001, Supp. Fig. 18A-B, p = 7.7×10^-47^ for all SynGO^111^ genes, permutation test). For example, many genes that code for proteins that localize to the postsynaptic density have nearby substitutions with large, highly neuron-specific predicted effects on CA (Fig. 6D). This again points to widespread divergence in the regulation of genes involved in neuronal connectivity in the human lineage.

Several individual genes were also enriched (Fig. 6E, Supp. Fig. 19). For example, fixed substitutions were enriched for large cell type-specific effects near *VWC2L* in fetal cortical neurons (Fig. 6E, FDR = 0.0016). *VWC2L* is a poorly studied paralog of an important regulator of neuronal and glial fate (*VWC2*) and specifically differentially expressed between the human and chimp alleles in developing hybrid motor neurons (Fig. 6F, example substitution in Fig. 6G). As another example, *FGF10*, a key secreted regulator of oocyte development, is enriched for cell type-specific effects in preGC cells (example substitution in Fig. 6H, FDR = 6.9×10^-5^). Collectively, these results provide clues as to the phenotypic consequences of positive selection on variants with cell type-specific effects.

Based on these findings, we reasoned that stronger positive selection on substitutions with cell type-specific effects could result in an increase in the cell type-specificity of the overall *cis*-regulatory landscape of some cell types. For example, substitutions with neuron-specific effects on CA could make existing neuronal CREs, on average, more specific to neurons. To test this, we computed the difference in the chromatin accessibility enrichment (CAE) between the two alleles at each site and compared the distribution of this metric for fixed and polymorphic sites. As CAE is a measure of how specific the accessibility of a CRE is to a particular cell type, a positive difference in CAE indicates that the derived allele makes the region more specific to that cell type whereas a negative difference in CAE indicates that it makes it less specific to that cell type.

Out of 16 cell types we initially tested, which included one type of fetal neuron, the only cell type with evidence for positive selection increasing cell type-specificity was fetal cortical excitatory neurons (Fig. 6I-J, α_CAE_ = 0.063, FDR = 0.0010). In contrast, we observed no such trend for any adult or non-neuronal fetal cell types, including adult cortical excitatory neurons (Fig. 6K, α_CAE_ = 0.0050, FDR = 0.46). To explore this further, we expanded our analysis to include 2 additional types of fetal and adult neuron. Across various possibly parameter combinations, we consistently observed the strongest evidence for positive selection increase cell type-specificity in the three fetal neuronal types (Supp. Fig. 20). Overall, our analysis suggests that positive selection has resulted in increased specificity of *cis*-regulatory elements to fetal neurons, potentially reducing pleiotropy and increasing the evolvability of *cis*-regulation in fetal neurons in recent human evolutionary history.

### Identification and characterization of RAGs and RALs

Based on the strong theoretical backing for the idea that variants with cell type-specific effects are more likely to be positively selected^118,119^ as well as the newfound empirical evidence detailed above, we reasoned that we could use this information to prioritize *cis*-regulatory elements that have undergone positive selection in the human lineage. To do this, we combined this information with the well-established concept that substitutions with reinforcing effects on phenotype are a hallmark of positive selection^23,121,122^. We developed a pipeline to identify genomic regions in which multiple fixed substitutions have reinforcing effects on a *cis*-regulatory element (see Methods). We term these elements RAGs (**R**einforcing **A**ccessibility **G**ain) and RALs (**R**einforcing **A**ccessibility **L**oss). Here, we focus specifically on RAGs and RALs that have reinforcing effects on the cell-type specificity of CA, rather than CA across all cell types.

We define RAGs and RALs as genomic regions with three or more substitutions within 1 kb that each increase or decrease the cell type-specificity of CA beyond a set cutoff, with no substitutions acting in the opposite direction (see Methods). For RAGs, this can occur either because a substitution is predicted to increase CA in the cell type of interest but not in most other cell types, or decrease CA in most other cell types but not the cell type of interest (and vice versa for RALs, Methods). For example, four out of five substitutions in the RAG near *PKD2L1*, an ion channel subunit, increase accessibility in fetal cortical excitatory neurons, but have weak effects in all other cell types (Fig. 7A). In contrast, the remaining substitution decreases accessibility in most cell types but is predicted to slightly increase accessibility in fetal cortical excitatory neurons (Fig. 7A). Applying this pipeline, we identify 6,971 hRAGs and 6,091 hRALs in the human lineage as well as 7,972 cRAGs and 6,922 cRALs in the chimp lineage (with the larger number in chimp due to a larger number of input sites; see Methods). The distribution of the number of both types of element across cell types was highly similar for human and chimp (Fig. 7B) and differences between cell types largely reflect the overall cell type-specificity of predicted effects on CA to each cell type rather than a tendency for those effects to be particularly clustered in a cell type.

**Figure 7:**
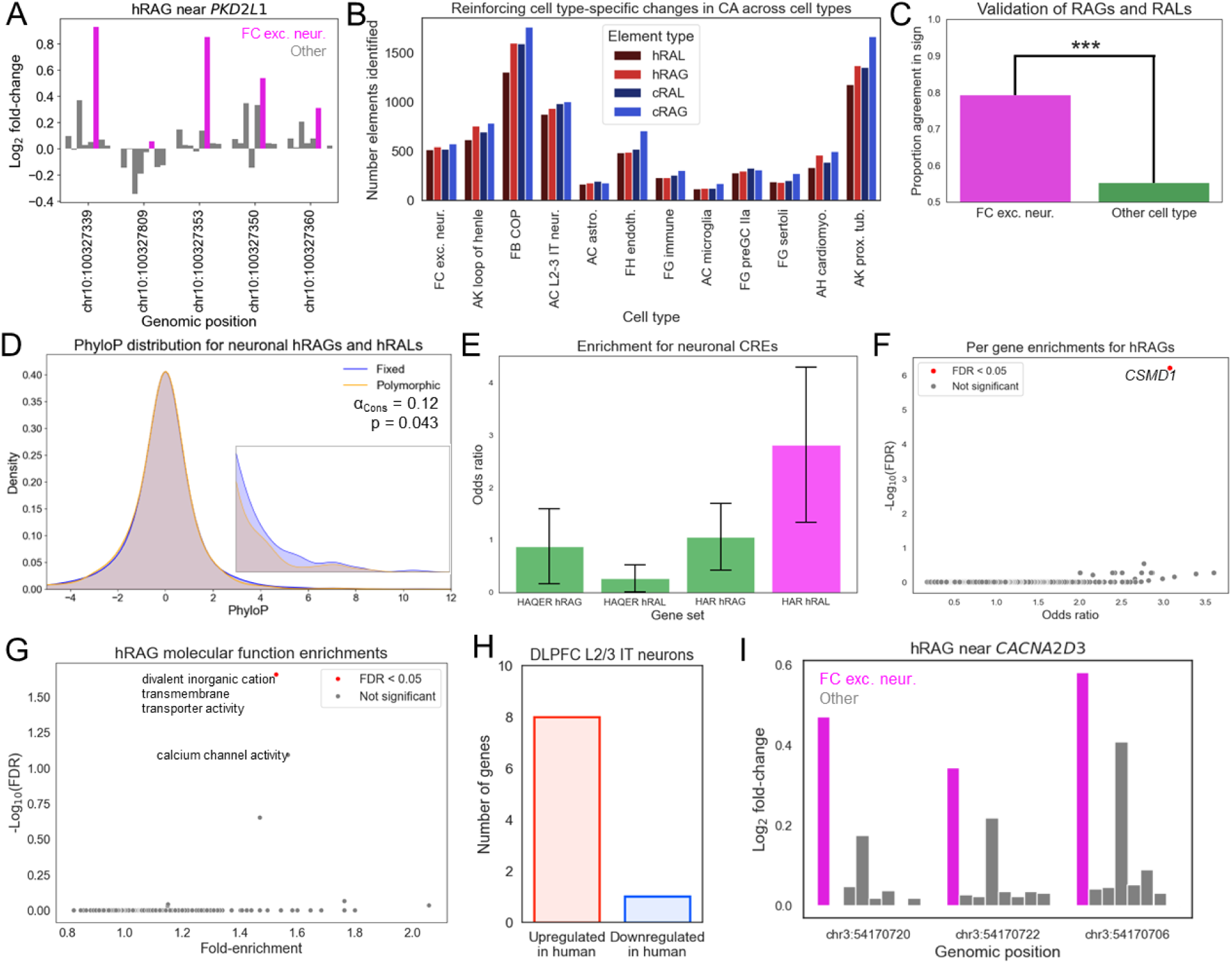
Mutually reinforcing cell type-specific changes in chromatin accessibility. **A)** Example of a **h**uman **R**einforcing **A**ccessibility **G**ain (hRAG) near *PKD2L1*. The magenta bars show the predicted log_2_ fold-change in CA in fetal excitatory neurons and the gray bars show the same thing for the other seven cell types used to identify the hRAG. **B)** Distribution of RAGs and RALs across cell types for human and chimpanzee. **C)** Validation of RAGs and RALs. The magenta bar shows the proportion agreement between the sign of CA change from the ChromBPNet predictions and experimentally measured differences in accessibility in iPSC-derived human-chimp hybrid excitatory neurons. The green bar shows the same thing for all other cell types. *** p < 0.001. **D)** Distribution of PhyloP scores for fixed and polymorphic sites in neuronal hRAGs and hRALs. The inset shows the distribution magnified in the range 3-12. The 90^th^ percentile of the polymorphic distribution was used to compute α_Cons_. **E)** Enrichment for neuronal hRAGs and hRALs in HARs and HAQERs relative to all other cell types. The error bars are the 95% confidence interval from a one-tailed Fisher’s exact test. Magenta indicates p < 0.05 and green indicates p > 0.05. **F)** Volcano plot showing enrichment of hRAGs near individual genes (dots). The Fisher’s exact odds ratio and -log_10_(FDR) are shown on the x- and y-axes respectively. **G)** Volcano plot showing enrichments for GO Molecular Function gene sets for hRAGs relative to cRAGs. **H)** Barplot showing number of upregulated and downregulated genes in L2/3 intratelencephalic (IT) neurons in the divalent inorganic cation transmembrane transporter activity category. **I)** Same as in (A) but an hRAG near *CACNA2D3* where the magenta bar represents the log_2_ fold-change in L2/3 IT neurons.

**Figure 8:**
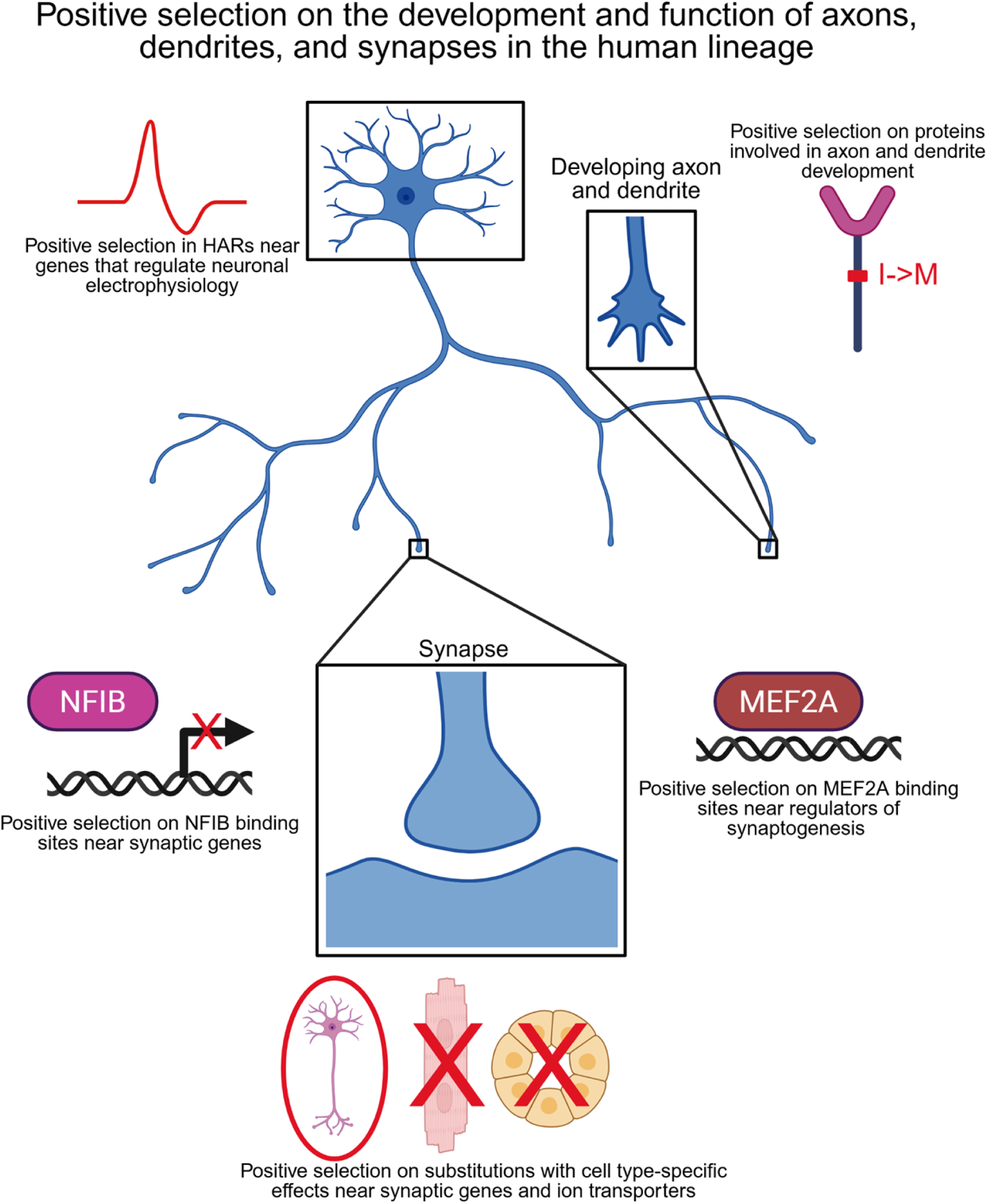
Schematic outlining positive selection on the development and function of neuronal projections discovered in this work.

To estimate the probability of observing each RAG/RAL by chance, we used common polymorphisms as a neutral null model, extending the logic in our selection test framework. Specifically, we asked whether the cell type-specific effect sizes in RAGs/RALs were greater than expected based on genome-wide polymorphisms using the binomial test (see Methods). We consider a RAG or RAL to be high-confidence if it has binomial p < 5×10^-7^ and medium-confidence otherwise. Across all cell types, we identified 434 high-confidence hRAGs, 341 hRALs, 486 cRAGs, and 399 cRALs. We used both medium- and high-confidence regions for the analyses below to increase power and avoid unduly penalizing cell types with a larger number of qualifying input sites (see Methods).

To experimentally validate RAGs and RALs, we focused on a recently published ATAC-seq dataset from human-chimp hybrid excitatory neurons^123^. Encouragingly, 79% of RAGs and RALs found in fetal cortical excitatory neurons that intersected differentially accessible peaks had the expected directionality for differential accessibility based on the predictions (Fig. 7D). In contrast, only 55% of RAGs and RALs found in other cell types had matching directionality (Fig. 7C, odds ratio = 3.1, p = 0.00013, Fisher’s exact test), confirming the accuracy of the predictions underlying RAGs and RALs. To provide additional evidence that hRAGs and hRALs were positively selected in neurons, we tested whether the fixed sites in hRAGs and hRALs generally had higher PhyloP scores. Consistent with positive selection, we observed larger PhyloP scores for fixed sites compared with polymorphic sites (Fig. 7D, α_Cons_ = 0.12, p = 0.043).

As hRAGs and hRALs were ascertained orthogonally to HARs and HAQERs, we tested if there was any overlap in these sets of regions. 18 and 19 RAGs and RALs intersect HAQERs and 46 and 50 RAGs and RALs intersect HARs (Supp. Fig. 21A), suggesting that HARs, HAQERs, RAGs, and RALs are largely in separate regions of the genome. Strikingly however, RALs that intersect HARs are enriched for excitatory neuron RALs (Fig. 7E), suggesting that one mechanism underlying positive selection in HARs may be selection for decreased CRE activity specifically in neurons. For example, four substitutions in a HAR near *MGAT5* are predicted to decrease accessibility in fetal cortical neurons but increase accessibility in several other cell types (Supp. Fig. 21B).

Finally, we explored how hRAGs and hRALs differ from their chimp counterparts as this could provide insight into which genomic changes underlie uniquely human phenotypes. As a first step in doing this, we tested whether hRAGs were enriched near any genes relative to the number of sites with cell type-specific effects that do not cluster in hRAGs (see Methods). Strikingly, the only gene that is enriched is *CSMD1*, which plays important roles in synapse development, is implicated in schizophrenia, and has increased expression from the human allele in human-chimp hybrid cortical neurons (Fig. 7F, FDR = 6.1×10^-7^, one-tailed Fisher’s exact test)^123,124^. *CSMD1* is also enriched for hRALs, but not for cRAGs or cRALs (Supp. Fig. 22A-C). Moreover, there are significantly more hRAGs and hRALs near *CSMD1* relative to their chimp counterparts (odds ratio = 1.72, p = 0.016 for gains, odds ratio = 1.65, p = 0.026 for losses), confirming that this signal is human-specific. Overall, this suggests that the *cis*-regulatory landscape of *CSMD1* has been considerably rewired in the human lineage.

We then used GREAT^95^ to test whether hRAGs called in neuronal cell types were enriched near any genes relative to cRAGs (as well as for RALs). The only significantly enriched category was “divalent inorganic cation transmembrane transporter activity” (Fig. 7F-G) which includes *PKD2L1* (Fig. 7A). In contrast, there are no significant enrichments for cRAGs, hRAGs, or cRALs in neurons (not shown). Interestingly, genes in this gene set are biased toward upregulation in L2/3 IT neurons with 8 out of the 9 significantly differentially expressed genes having higher expression from the human allele (Fig. 7H, p = 0.017, binomial test), which is the expected direction as hRAGs generally have increased accessibility from the human allele in the cell type they are called in. Moreover, this bias remains at a more relaxed FDR cutoff for differential expression (15 out of 20 genes upregulated, binomial p = 0.02, at FDR < 0.25 for differential expression). For example, an L2/3 IT neuron hRAG (Fig. 7I) is adjacent to *CACNA2D3*, a calcium channel subunit associated with ASD^125^ that has increased expression in human L2/3 IT neurons and deep layer excitatory neuron types relative to the expression in chimps, but only has higher expression relative to macaque and marmoset in deep layer neurons suggesting the change in *CACNA2D3* expression may be most important in those cell types (Supp. Fig. 23). Collectively, our results suggest that hRAGs and hRALs represent a class of candidate CREs that may have played an important role in human evolution.

## Discussion

Throughout the last few decades, our ability to detect positive selection has been intertwined with our ability to accurately predict variant effects. In the last few years, there has been a rapid expansion and improvement in our ability to predict the effects of genetic variation on an increasingly broad variety of molecular phenotypes, primarily driven by the development of new deep learning methods^37–44^. Here, we provide evidence that a well-established method to detect positive selection on the protein-coding regions of genes can be straightforwardly generalized to essentially any variant effect prediction. Although this concept has been applied to specific binary classifications of variant annotation before^25,28^, our results suggest that all these methods represent corner cases of a diverse range of possible tests, enabling new insights into how positive selection has shaped genetic and phenotypic variation across the tree of life

Applying this concept to human evolution, we demonstrate that this generalized framework is able to recapitulate and extend well-established results as well as provide strong support for longstanding evolutionary theories. For example, although previous studies have demonstrated that Alu elements are frequently co-opted as *cis*-regulatory elements^105^, we provide empirical evidence that Alu elements experienced stronger positive selection on chromatin accessibility relative to other repetitive regions in the human lineage. In addition, we provide strong support for the longstanding theoretical prediction that substitutions with more cell type-specific effects experience stronger positive selection^118,119^. As this is an important prediction of the idea that antagonistic pleiotropy is a major factor governing evolution^118,119^, this result provides support for the idea that a lack of antagonistic pleiotropy for non-coding sites is one of the primary reasons that positive selection tends to act on substitutions that affect *cis*-regulation (as opposed to substitutions that alter *trans*-regulation or protein sequences)^118,119^.

We anticipate that further application of this framework will enable researchers to tease apart precisely which aspects of *cis*-regulation (as well as other phenomena) are the main drivers of adaptive evolution. However, it is important to note that using non-binary variant effect predictions (e.g., conservation or log fold-change) is inherently more complex than using binary predictions. For example, it is important to investigate whether results are robust to the selection of different cutoffs when computing α (Supp. Text 6). In addition, it is difficult to compare two sets of sites if they have very different distributions of variant effect predictions. For example, the PhyloP distributions for nonsynonymous substitutions are more uniform (i.e., many sites are highly conserved) than for non-coding substitutions, making direct comparison between the two difficult. Overall, many evolutionary questions can be addressed with this framework in its current form and we anticipate that further development adapting it to more complex variant effect predictions will further expand the range of addressable questions.

In addition to using this framework to explore general principles governing human evolution, we also use it to provide new insight into what genetic differences underlie uniquely human traits. For example, we find evidence for positive selection on the protein-coding sequences of key regulators of heart development and experimentally validate the effects of a human-specific substitution in *ABCC9* on potassium channel activity. One striking finding is diverse signals of positive selection on the coding and non-coding regions of genes involved in the development and function of neuronal connections. For example, we find evidence for positive selection on nonsynonymous substitutions in proteins that regulate the development of neuronal connectivity, chromatin accessibility in NFIB binding sites, and the expression of synaptic genes, as well as on chromatin accessibility near genes that regulate the electrophysiological properties of neurons.

Although there has been much focus on the increased brain size and slower rate of neuronal maturation in humans, our results suggest that positive selection may also have resulted in important, fine-scale changes in neuronal connectivity and function specific to humans. The ∼86 billion neurons in the human brain make highly regulated, stereotyped connections between different neuronal types and brain regions that are shaped by genetic variation^126–128^. Combined with the highly complex gene regulatory landscapes of genes involved in neuronal wiring^129^ (56 out of 100 genes in the largest gene deserts are cell-surface proteins regulating neuronal wiring compared to 6 transcriptional regulators, Supp. Fig. 24), this represents a massive target space in which positive selection could act on genetic variation to fine-tune cognition and behavior. Although MRI-based methods have enabled exploration of gross connectomic differences between humans and other apes^130,131^, large-scale connectomics using comparative electron microscopy and postmortem tracing experiments will be vital in understanding fine-scale differences in neuronal connectivity that are unique to humans. Furthermore, humanized mouse models as well as more advanced *in vitro* and xenotransplantation methods that are able to model the development and function of fine-scale neuronal connectivity will be essential in understanding the genetic basis of uniquely human cognitive abilities.

Finally, building off our finding that substitutions with cell type-specific effects are more likely to be positively selected, we identify RAGs and RALs, hotspots of reinforcing, cell type-specific changes in chromatin accessibility, many of which were likely under positive selection in the human lineage. We validate many of these predicted elements in developing neurons and provide evidence that they may have affected human neuronal electrophysiology. Moreover, these efforts provide a list of *cis*-regulatory elements that likely had important phenotypic effects in the human lineage (by virtue of being positively selected) as well as strong predictions of the affected cell types. In the future, further characterization of the genes and genetic variants implicated by our framework could provide deeper insight into the genetic changes that make us human.

## Methods

### Human fixed and polymorphic substitution processing

Human genomic variation VCF files were downloaded from Gnomad version 3.1.2^48^. To subset to a set of high-confidence, common polymorphisms, we required that each polymorphism have a minor allele frequency (MAF) of at least 0.1 and at least 5 alleles measured in at least one of the seven major ancestry groupings in Gnomad using bcftools^132^. Although this likely makes our analysis somewhat conservative compared to analyses that only use African polymorphisms due to the bottlenecks that non-African populations underwent, it helps ensure that the fixed sites are truly fixed across many human populations and increases the sample size of polymorphic sites (as some sites only have moderately high derived allele frequency in non-African populations).

To identify fixed human-chimp differences, we used our previously published set of high-confidence human-chimp substitutions, further removing all sites with polymorphisms that had MAF > 0.001 in at least one ancestry group from Gnomad. We classified substitutions as human-derived if the gorilla GorGor6 reference allele matched the chimp PanTro6 reference allele, and chimp-derived if the gorilla allele matched the human hg38 allele. For human polymorphisms, we classified an allele as ancestral if it matched the chimp and gorilla reference alleles, derived if it didn’t match the chimp or gorilla alleles and the chimp and gorilla reference genomes had the same base, and ambiguous otherwise. We then filtered out any polymorphic or fixed sites that failed to uniquely, reciprocally liftover between hg38 and PanTro6 and hg38 and GorGor6.

To classify substitutions as nonsynonymous, synonymous, and non-coding, we used Ensembl VEP^133^ v109 through the web portal to predict the consequences of changing the human reference base to the alternate base. We classified substitutions as nonsynonymous if they were flagged as missense by VEP; synonymous if they were flagged as synonymous and were not flagged as missense, 5’ UTR, or 3’ UTR; and non-coding if they were not flagged as missense, 5’ UTR, or 3’ UTR. For example, if a substitution was classified as both intronic and 3’ UTR, that substitution would be filtered out of the set of non-coding substitutions. If instead that substitution were classified as both nonsynonymous and 3’ UTR, it would be counted as nonsynonymous. We assigned non-coding sites to the gene that had the nearest human-chimp orthologous promoter^71^.

For non-coding substitutions, we additionally removed sites in low complexity regions by filtering out any sites that intersected regions in the files “GRCh38_AllHomopolymers_ge7bp_imperfectge11bp_slop5.sort.bed” and “GRCh38_AllTandemRepeats.sort.bed” available from: https://github.com/genome-in-a-bottle/genome-stratifications/blob/master/GRCh38/LowComplexity/GRCh38-LowComplexity-README.md. We also assigned sites to repetitive elements using the file “hg38_2020_rmsk.bed” available from: https://repeatbrowser.ucsc.edu/data/. We noticed that several common repeats (e.g., L1PA2 and L1PA3) contained a disproportionate number of polymorphisms relative to fixed sites, potentially for technical reasons that could confound our results. To address this, we removed all sites intersecting a set of blacklist repetitive elements that had a Fisher’s exact odds ratio < 0.75 (i.e., there are at least 33% more polymorphisms in a type of repetitive element relative to fixed sites than expected by chance given the background ratio of fixed to polymorphic sites) indicating an excess of polymorphisms.

### Computing PhyloP scores

We used the 447-way alignment of placental mammals as input to compute PhyloP scores^49,50^. As estimates of the neutral substitution rate were not available at the time that PhyloP scores were computed, we derived this information following the same methodology used by the Zoonomia consortium, with the exception that we additionally checked for synteny against the primate last common ancestor (LCA) genome. Briefly, we first extracted a fasta file for fullTreeAnc238, corresponding to the ancestral genome of the Boreoeutherian LCA, from the hal file using hal2fasta from Cactus v2.6.13^134^ and then called repetitive regions using RepeatMasker v4.1.6^135^. We then removed any repeats that were RNA, microsatellites, low complexity, or unknown, as well as other miscellaneous categories recommended by the Zoonomia consortium^49^. We further removed any repeats that were not syntenic with any of fullTreeAnc5 (Xenarthan LCA), fullTreeAnc13 (Afrotherian LCA), fullTreeAnc115 (LCA of Euarchontoglires), fullTreeAnc237 (Laurasiatherian LCA), and Primates000 (Primate LCA), then lifted over to fullTreeAnc239 (the placental mammal LCA). We then selected 100,000 random sites in the remaining repeats and used PhyloFit from phast v1.5^136^ to generate the model file. We used this model file for all PhyloP score computation.

We will describe how PhyloP scores were computed referenced to the human genome as an identical procedure was used for the house mouse and the black flying fox with the only difference that it was referenced to the respective genomes of those species. We split the genome into 10 megabase segments (or smaller if a contig was less than 10 megabases) and converted to a MAF file using hal2maf, removed duplicate sites with mafTools^137^ mafDuplicateFilter v0.1, and then masked the reference species with phast^138^ maf_parse. We then computed PhyloP scores with phast phyloP^136^ using the following parameters: --msa- format MAF --method LRT --mode CONACC -d 6 --wig-scores. Finally, we merged the files for all the different genome chunks and intersected them with the files of fixed and polymorphic sites using bedtools^139^ v2.30.0 intersect.

In addition to PhyloP scores, we counted the number of species in each alignment block, removing any that had greater than 10% N or - characters in that block as an approximation of the statistical power available to complete PhyloP scores. For analyses of positive selection on conserved non-coding sites, we removed all sites for which this value was less than or equal to 250. This ensures that at least 8 non-primate species were present in the alignment as there are 242 primates. Similarly, for house mouse and black flying fox, we removed sites for which this value was less than or equal to 61 (the number of rodents and lagomorphs in the alignment plus 8) and 38 (the number of bats in the alignment plus 8) to ensure approximate equivalency across species. All this information was intersected with the list of fixed and polymorphic sites using bedtools^139^.

### House mouse fixed and polymorphic site processing

We used the polymorphism data available from Harr et al^66^. To process the data for use in our test, we first extracted the allele frequencies and major/minor alleles for *Mus musculus domesticus* and *Mus spretus*. We considered a site to be fixed if the major allele frequency for *Mus musculus domesticus* was 1 and polymorphic otherwise. We filtered to only sites that were fixed in *Mus spretus* to ensure a fair comparison between fixed and polymorphic sites and filtered out all polymorphic sites with minor allele frequency < 0.1. We then extracted the genotype information from *Mus musculus*, *Mus spretus*, and *Mus caroli* by first creating a maf file from the Zoonomia hal file using hal2maf with the --onlyOrthologs flag. We then removed duplicates with mafTools, used a custom python script to identify all sites that differ between *Mus musculus* and at least one of the other species, and intersected with the list of fixed and polymorphic sites. To determine the derived allele, we used the same strategy outlined for human, replacing human with *Mus musculus*, chimp with *Mus spretus*, and gorilla with *Mus caroli*. We then intersected with the PhyloP scores, the number of species in the alignment for *Mus musculus*, and found the nearest gene using a published set of transcription start sites^140^. We then used the ENSEMBL VEP^141^ web portal v112 with GRCm39 to classify sites as nonsynonymous, synonymous, UTR, and non-coding as described above.

### Black flying fox fixed and polymorphic site processing

The VCF file from He et al.^67^ kindly shared by the authors. We restricted to polymorphisms with minor allele frequency in the range 0.1-0.9. We then extracted and processed genotype information for *Pteropus alecto*, *Pteropus vampyrus*, and *Rousettus_aegyptiacus*, determined the derived allele, and intersected with PhyloP and the number of species in the alignment for *Pteropus alecto* as described for house mouse. As the genome annotation for *Pteropus alecto* is not as high quality as that of human and mouse, we instead intersected the sites with the TOGA^142^ genome annotation based on the Hg38 annotation available here: https://genome.senckenberg.de/download/TOGA/human_hg38_reference/Chiroptera/Pteropus_alecto__black_flying_fox__pteAle1/. We considered a site to be non-coding if it did not intersect any CDS in the annotation.

### General strategy for testing for positive selection

Computing α or testing for positive selection using Fisher’s exact test requires selecting a cutoff to split the variant effect prediction distributions (Fig. 1A). We reasoned that a consistent way to pick a cutoff across different comparisons would be to use the distribution of variant effects for the polymorphisms. For the same variant effect prediction metric, different cutoffs can be used to test for different things. For example, picking the 95th percentile of the non-coding PhyloP distribution would test for positive selection on highly conserved sites. On the other hand, picking the 60th percentile would test for positive selection on both moderately and highly conserved sites. However, picking the 60th percentile for the nonsynonymous PhyloP distribution would test for positive selection on highly conserved sites because a much larger proportion of coding sites are highly conserved. In general, we chose the cutoff based on what we were interested in testing. Furthermore, we always tested multiple different cutoffs to ensure that our results were qualitatively similar regardless of which cutoff was chosen (Supp. Text 6). In addition, because we are selecting the cutoff based on the polymorphic distribution, at least one site must be on the border between the two distributions and putting it in either the neutral or non-neutral bin is arbitrary. To deal with this, we compute α and the Fisher’s exact p-value both ways and report the mean of the two resulting values for α and the p-value respectively.

When testing for positive selection on genes or gene sets using PhyloP scores, we avoided picking a cutoff by using the one-sided Mann-Whitney U test to test for differences in the median of the fixed and polymorphic PhyloP distributions. We found that this method had more statistical power than picking a cutoff and using Fisher’s exact test. For non-coding sites, we only considered a gene set to be significant if it had Mann-Whitney U test FDR < 0.05 and one-sided Fisher’s exact p-value < 0.05 using the 90th percentile of the polymorphic PhyloP distribution as a cutoff to ensure that highly conserved sites were contributing to the observed signal. We used a nominal p-value cutoff for Fisher’s exact test because the requirement that the gene set had FDR < 0.05 for the Mann-Whitney U test ensures a low false discovery rate and additionally requiring Fisher’s exact FDR < 0.05 would be very strict given the lower power (especially at strict percentile cutoffs, see Supp. Text 6).

In general, if there were fewer than 1,000 total (fixed plus polymorphic) sites, we used Fisher’s exact test to test for positive selection and did not compute alpha as it is unstable when there are few sites. To combine information across cell types with MASH^87^, we used a two-sided test as this is necessary when using MASH. Otherwise, we used a one-sided test. Throughout, we use two different methods to test for positive selection and test if there is stronger evidence for positive selection on one set of sites compared to another. If we were focusing on a small number of categories (e.g., HARs, HAQERs, cell type-specificity bins), then we used a bootstrapping to compute confidence intervals for α. We randomly sampled (with replacement) sites from the polymorphic and fixed distributions and recomputed alpha 1,000 times. To compare the confidence intervals for different bins of sites (e.g., comparing the most cell type-specific bin with a other bins), we down-sampled the number of fixed sites in each bin to the minimum number of fixed sites across all bins and did the same for the polymorphic sites, following previously established methods^10^. On the other hand, if there were many categories (e.g., from Human Phenotype Ontology or TF binding sites), we instead used Fisher’s exact test to determine statistical significance and compared confidence intervals from Fisher’s exact test to determine if there was stronger evidence for positive selection on one set of sites or another. In general, these two methods give very similar results.

To perform the asymptotic version of the test, which corrects for segregating mildly deleterious polymorphism, we split polymorphisms into 8 bins such that each bin contained all the polymorphic sites with minor allele frequency in a specific range, as previously described^30^. The lowest minor allele frequency bin was 0.1-0.2 and the highest was 0.8-0.9. We then computed alpha in each bin and fit an equation of the form alpha = a + b*log(MAF + c) using the curve_fit function in scipy to each (minor allele frequency, alpha) pair. We then estimated alpha by evaluating the function at MAF = 1.

For the non-asymptotic version of the test, it is also necessary to pick a maximum and minimum DAF cutoff. In general, we used symmetric cutoffs so that potential inaccuracies in determining which allele was derived would not affect our results. The only exception to this was when testing for positive selection using large sets of nonsynonymous substitutions and the genome-wide test for non-coding substitutions, where we used sites with DAF in the range 0.5-0.9 because alpha was generally very negative genome-wide in lower DAF bins for nonsynonymous substitutions and we wanted to be able to fairly compare the nonsynonymous and non-coding results. For analysis of nonsynonymous gene sets, we used sites with DAF in the range 0.1-0.9 to maximize the number of gene sets that were testable. In all other cases, we only used sites with DAF in the range 0.25-0.75.

To generate plots of the fixed and polymorphic distributions, we smoothed the distributions using a Gaussian kernel-density estimate (KDE) using the the scipy gaussian_kde function with bw_method=0.3 and plotted the resulting KDE.

### Testing for enrichment of large-effect CA changes near genes and gene sets using GREAT

We split sites into 2 bins, those with predicted effect size greater than some cutoff and those with predicted effect size less than another cutoff. For predicted absolute log_2_ fold-change in CA, these cutoffs were either 0.25 and 0.1 or 0.1 and 0.025, respectively. We used the latter cutoffs when fewer than 10% of sites had log_2_ fold-change greater than 0.25 to ensure adequate power to detect enrichments. We then uploaded sets of sites with effect size greater than the upper cutoff as the foreground and effect size less than the lower cutoff, combined with the foreground, as the background and ran GREAT^95^ with default parameters, except that we required that there be at least 5 genes observed for each category and at least 20 genes annotated for each term. The only exception to this was when testing for enrichment of cell type-specific effects near specific genes. In this case, we used substitutions with dTau > 0.9 as the foreground and dTau < 0.5 to create the background. We also did not put any restriction on the number of genes observed when testing for enrichment near single genes.

To validate significant results and ensure that the choice of cutoff was not qualitatively impacting the results, we used a permutation procedure. We first computed the mean of the variant effect prediction on all sites for which the nearest gene was in a particular category. We then randomly sampled an equal number of sites from the set of all sites (regardless of which gene they were near) and computed the mean variant effect prediction for that sample of sites. We repeated this sampling procedure 10,000 times to build a null distribution and computed the Z-score and p-value from comparing the observed value to the null distribution.

### Testing for positive selection on conserved nonsynonymous and non-coding sites

To test for positive selection on nonsynonymous sites genome-wide, we restricted to polymorphisms with DAF in the range 0.5-0.9 and compared the fixed and polymorphic PhyloP distributions with the cutoff as the 60th percentile of the polymorphic PhyloP procedure. For haploinsufficient genes, we restricted to genes with Gnomad^48^ probability of loss of function intolerance (pLI) greater than 0.9 as has been done previously and repeated this. We considered a gene to be depleted of nonsynonymous variants in human populations if it had a Gnomad^48^ missense Z-score greater than 2, very depleted of nonsynonymous variants if it had a Z-score greater than 4, and not depleted of nonsynonymous variants if it had Z-score less than or equal to 2.

To test for positive selection on nonsynonymous substitutions in genes associated with human phenotypes, we restricted to polymorphic sites with DAF in the range 0.1-0.9 to enable more categories to be tested. As this includes many deleterious polymorphisms leading to a negative genome-wide α_Cons_, we quantile normalized fixed and polymorphic PhyloP scores for only this analysis so that they had approximately the same genome-wide distribution. We then tested all Human Phenotype Ontology (HPO)^55^ categories with greater than 15 genes, at least 50 fixed nonsynonymous sites, and at least 30 polymorphic nonsynonymous sites assigned to that category.

We tested for positive selection on non-coding sites genome-wide identically to nonsynonymous sites, except that we used the 95th percentile of the polymorphic distribution. To test for positive selection on non-coding sites near genes associated with particular phenotypes, we restricted to HPO categories with at least 15 genes, at least 100 fixed sites, and at least 50 polymorphic sites using the Mann-Whitney U test and Fisher’s exact test with cutoff equal to the 90th percentile of the polymorphic distribution. We considered a gene to have significant evidence for positive selection if it had Mann-Whitney FDR < 0.05 and Fisher’s exact test p-value < 0.05, with the latter condition ensuring that highly conserved sites are at least partially driving the signal for that gene set. To prioritize *ARX*, we tested all genes in the Progressive intellectual disability gene set with at least 100 fixed sites and 50 polymorphic sites and tested for positive selection with Fisher’s exact test with the cutoff being the 90th percentile of the polymorphic distribution. *ARX* had the lowest p-value.

To test for positive selection on each chromosome, we split sites by chromosome and used the above bootstrap procedure to estimate a confidence interval for α_Cons_. To further explore positive selection on the X chromosome, we used a one-sided Fisher’s exact test to test for positive selection after restricting to sites in previously published selective sweeps^65^ or near multi-copy testis expressed genes^63^, or removing those sites. To test for positive selection on human phenotypes, we used an identical procedure to the one outlined above for testing genome-wide, except that we restricted to sites on the X chromosome. We identified the two substitutions in ultraconserved non-coding elements by intersecting sites on the X chromosome with the list of ultraconserved non-coding elements^74^ available here: https://epd.expasy.org/ucnebase.

### Membrane potential assessments by voltage-sensitive fluorescent dye

Four batches of HEK293 cells stably expressing Kir6.1/wtSUR2B and four batches expressing Kir6.1/SUR2B[N1538D] K_ATP_ channels were generated independently, as described previously^143^. Cells were cultured in DMEM containing 10% FBS, 100 U/ml penicillin, 0.1 mg/ml streptomycin and 4 μg/ml doxycycline. Transparent 96-well cell-culture plates were coated with poly-L-lysine (0.1 mg/ml) one day before stable cells were plated at an appropriate density (20 - 200 cells per view in the following assay). After one day in culture, the medium was discarded and the cells were washed twice with low [K] buffer (NaCl 139 mM, KCl 1 mM, CaCl_2_ 2 mM, MgCl_2_ 1 mM, HEPES 10 mM, Glucose 10 mM, pH adjusted to 7.4 with NaOH). Cells were then incubated with 3 μM DiBAC4(3) in low K buffer for at least 20 min. DiBAC4(3) is a voltage-sensitive dye, that accumulates in cells at depolarized voltages. Increased K conductance leads to cell hyperpolarization and reduced DiBAC4(3) fluorescence. The stably transfected cells also express mCherry as a marker for cell detection. Separate images of green (DiBAC4(3)) and red (mCherry) fluorescence were taken with EVOS M5000 imaging system (10x lens). Exported tiff images were analyzed by CellProfiler with custom designed programs to assess green fluorescence in red detected objects (cells). Multiple measurements of each cell were exported as a csv format file and the fluorescence intensities of all cells, as well as the mean value in each image were extracted with a custom script written in Python^143^. The average intensity value of duplicate images was calculated as one data point for the statistical analysis and data presentation. Data points in Fig. 1G are from four batches of the two different cell lines measured on different days.

### Comparing positive selection in human, house mouse, and black flying fox

As we were unable to use ENSEMBL VEP due to differences in genome annotation quality, we considered a site to be non-coding if it did not lie in a CDS. We then restricted to 3-way one-to-one human, house mouse, black flying fox orthologs (downloaded from ENSEMBL Biomart) and computed alpha using the 95th percentile of the polymorphic PhyloP distribution as a cutoff to test for positive selection on highly conserved sites. We also computed alpha using the same parameters using the asymptotic method.

### Data processing for training of ChromBPNet models

To process the adult cortical microglia^77^ (Synapse accession syn26207321), fetal chondrocyte data^78,79^ (GEO accession GSE214394 and GSE122877), and fetal cortical excitatory neuron and neural progenitor data^80^ (dbGaP accession phs001958.v1.p1), we first trimmed reads with Trim Galore^144^ v0.6.7 using CutAdapt^145^ v1.18 in paired end mode using the following parameters: trim_galore --paired --stringency 5 --length 20 --fastqc --consider_already_trimmed 1000. We then aligned reads to the human Hg38 genome using bowtie2^146^ v2.3.5.1 with parameters: -X 2000 --very-sensitive-local -p 16. We then removed duplicate reads using picard^147^ MarkDuplicates with parameters REMOVE_DUPLICATES=true DUPLICATE_SCORING_STRATEGY=RANDOM and then used the command samtools^132^ v1.9 view -b -q 10 to remove low quality alignments. Finally, we sorted and merged all samples from control individuals with APOE3/APOE3 for the adult microglia study, and all samples for the other 3 cell types and used the resulting bam files as input to ChromBPNet. We called peaks by first converting to bed files with bedtools^139^ v2.30.0 bamtobed and then calling peaks with macs2^148^ v2.2.7.1 using the following parameters: -f BED -p 0.01 --nomodel --shift 75 --extsize 150 -B --SPMR --keep-dup all --call-summits.

For the fetal heart data^81^, we downloaded the data and metadata mapping sample and barcode to each cell type from the gene expression omnibus (GEO, accession GSE181346). We then used a custom python script to split the fragments files derived from standard single nucleus ATAC-seq data processing piplines by cell type, pooling across all samples.

For the adult kidney data^82^, we downloaded the fragments files from GEO (accession number GSE262931) and the metadata mapping sample and barcode to cell type was provided by the authors. We then used a custom python script to split the fragments files by cell type, pooling across all samples. We then called peaks on the fragments file for each cell type with macs2 with -f BEDPE -g hs --nomodel --shift 100 --ext 200 -p 0.01 --keep-dup all.

Adult heart data^83^, already split into cell type-specific files, was downloaded from the CARE data portal at http://ns104190.ip-147-135-44.us/data_CARE_portal/snATAC/bedfiles in bed format. We then called peaks using macs2 with parameters -f BED -p 0.01 --nomodel --shift -100 -- extsize 200 --call-summits --keep-dup all for each cell type.

The fetal brain data^84^ was downloaded from https://storage.googleapis.com/linnarsson-lab-human/ATAC_dev/10X and the sample and barcode to cell type mapping was downloaded from https://github.com/linnarsson-lab/fetal_brain_multiomics. We used a custom script python script to split files by cell type and pool across samples and then called peaks on each sample with macs2 using the following parameters: -f BEDPE -g hs --nomodel --shift 100 --ext 200 -p 0.01 -- keep-dup all.

The data for fetal gonads^86^ was downloaded from the BioStudies repository (accession E-MTAB-10570) and the metadata mapping sample and barcode to cell type was shared with us by the authors. A custom python script was used to split files by cell type and pool across samples, with female and male samples kept separate given the sex-specific aspects of gonad development. We called peaks on the fragments files using macs2 with the parameters: -f BEDPE -g hs --nomodel --shift 100 --ext 200 -p 0.01 --keep-dup all.

We downloaded the cell type-specific adult brain data^85^ from http://catlas.org/catlas_downloads/humanbrain/bedpe and the peaks from http://catlas.org/catlas_downloads/humanbrain/cCREs. We then used bedtools bedpetobam to convert to a bam file.

### Acronyms for cell types

We included 34 cell types in our analysis:

| Cell type full name | Cell type abbreviation | Dataset |
| --- | --- | --- |
| Adult neocortical microglia | AC microglia | Kosoy et al. 2022 |
| Fetal Chondrocytes | F. chond | Rice et al. 2023, Richard et al. 2020 |
| Fetal cortical excitatory progenitors | FC prog. | Liang et al. 2021 |
| Fetal cortical excitatory neurons | FC exc. neur. | Liang et al. 2021 |
| Fetal heart arterial endothelial cells | FH endoth. | Ameen et al. 2022 |
| Fetal heart cardiac fibroblasts | FH fibro. | Ameen et al. 2022 |
| Fetal heart pericytes | FH peri. | Ameen et al. 2022 |
| Fetal heart endocardium | FH endocard. | Ameen et al. 2022 |
| Adult kidney proximal tubule cells | AK prox. tub. | Loeb et al. 2024 |
| Adult kidney Loop of Henle cells | AK loop of henle | Loeb et al. 2024 |
| Adult heart smooth muscle cells | AH smooth musc. | Hocker et al. 2021 |
| Adult heart ventricular cardiomyocytes | AH cardiomyo. | Hocker et al. 2021 |
| Fetal cerebellum purkinje inhibitory neurons | FCB purk. inh. neur. | Mannens et al. 2024 |
| Fetal brain committed oligodendrocyte progenitors | FB COP | Mannens et al. 2024 |
| Fetal brain floor plate | FB fl. plate | Mannens et al. 2024 |
| Fetal brain intermediate progenitors | FB inter. prog. | Mannens et al. 2024 |
| Fetal cerebellum glioblasts | FCB glioblast | Mannens et al. 2024 |
| Fetal brain dopaminergic neurons | FB DA neur. | Mannens et al. 2024 |
| Fetal spine dorsal root ganglion neurons | FS DRG neur. | Mannens et al. 2024 |
| Fetal brain roof plate | FB ro. plate | Mannens et al. 2024 |
| Fetal brain serotonergic neurons | FB 5-HT neur. | Mannens et al. 2024 |
| Fetal gonads Sertoli cells | FG sertoli | Garcia-Alonso et al. 2024 |
| Fetal gonads pre-granulosa cells IIa | FG preGC IIa | Garcia-Alonso et al. 2024 |
| Fetal gonads neural cells | FG neur. | Garcia-Alonso et al. 2024 |
| Fetal gonads immune cells | FG immune | Garcia-Alonso et al. 2024 |
| Fetal gonads Leydig cells | FG leydig | Garcia-Alonso et al. 2024 |
| Adult neocortical layer 2/3<br>intratelencephalic neurons | AC L2-3 IT neur. | Li et al. 2023 |
| Adult neocortical VIP+ inhibitory neurons | AC VIP inh. neur. | Li et al. 2023 |
| Adult neocortical SST+ inhibitory neurons | AC SST inh. neur. | Li et al. 2023 |
| Adult neocortical PVALB+ inhibitory<br>neurons | AC PVALB inh.<br>neur. | Li et al. 2023 |
| Adult neocortical oligodendrocytes | AC Oligo. | Li et al. 2023 |
| Adult putamen D1 medium spiny inhibitory<br>neurons | AP D1 inh. neur. | Li et al. 2023 |
| Adult amygdala neurons | AA neur. | Li et al. 2023 |
| Adult neocortical astrocytes | AC astro. | Li et al. 2023 |

### Training and inference with ChromBPNet models

ChromBPNet^37^ is system to train convolutional neural network models that predict chromatin accessibility (CA) from DNA sequence. The models are trained on ATAC-seq or DNASE-seq chromatin accessibility data. For a DNA sequence of width 2114 bases, the models predict a chromatin accessibility profile at single-base resolution for a 1000-base central window.

The trained models can predict the CA profiles of sequence that the models have never seen before, for example, sequences derived by making one or more substitutions to natural DNA sequence. In addition, the predictions are cell-type specific, that is, the model implicitly learns the effects on CA of transcription factors (TFs) as they are present in a particular cell type. This allows precise comparisons of profile differences in different cell types. Moreover, since *trans* factors are conserved relative to *cis* sequence changes, the models can be used to compare different species. For example, previous work has shown that models trained on data from a single species^42^ can be used to accurately predict the properties of *cis*-regulatory elements in species hundreds of millions of years diverged. Given the much smaller divergence time (around seven million years) of humans and chimpanzees, this suggests that the *trans*-regulatory environments of human and chimp cells are likely highly similar.

Notably, the models are trained to “factor out” bias due to the Tn5 transposase used in the assays, permitting nearly-bias-free prediction of accessibility^37^. This is accomplished by training an auxiliary model on low-read data to capture the Tn5 bias alone, freezing its parameters, and using it during training of the main model to remove sequence-specific Tn5 bias.

One limitation of the models is that predictions on entirely non-natural sequence are likely to be less accurate than those on near-to-natural sequence (one, or a few, substitutions from natural sequence); however, for our purposes, we are only interested in near-to-natural sequence.

We trained ChromBPNet models on ATAC-seq data from 34 pure populations of human cell types from a variety of organs, body systems, and developmental stages and predicted the effects of all fixed human-chimp differences as well as common human polymorphisms (derived allele frequency (DAF) > 0.1) as input to our framework (Fig. 3A). We performed these predictions by substituting the alternative base for the base in the human reference genome in all cases, except when identifying RAGs and RALs for which we also performed predictions by substituting the alternative base for the base in the chimp reference genome.

The ChromBPNet models were trained with the following values of the “bias” parameter:

| Cell type | ChromBPNet bias parameter |
| --- | --- |
| Kosoy et al. 2022 | 0.5 |
| Hocker et al. 2021 | 0.3 |
| Loeb et al. 2024 | 0.3 |
| Rice et al. 2023, Richard et al. 2020 | 0.2 |
| Mannens et al. 2024 | 0.4 |
| Garcia-Alonso et al. 2024 | 0.4 |
| Ameen et al. 2022 | 0.4 |
| Liang et al. 2021 | 0.25 |
| Li et al. 2023 | 0.25 |

To plot the predicted effects on CA, we first multiplied the profile distribution by the total counts to obtain the predicted per-base chromatin accessibility for that region. We then smoothed the resulting trace using the scipy function gaussian_filter1d with the sigma parameter equal to 4. As the total counts depends partially on the read depth of the data used to train the model, one cannot directly compare total counts across models without normalization. However, one can compare log fold-changes or other variant effect predictions derived from ChromBPNet across models.

### Combining information across cell types with MASHR

To combine information for tests for selection on CA for genes and TFBS, we converted two-sided Fisher’s exact test p-values to Z-scores and input that to MASHR^87^ v0.2.79 using ASHR v2.2.63. We then computed canonical and data-driven covariance matrices using 5 principal components, estimated the null correlation using a z-score threshold of two, and ran then ran MASH. We considered genes or TFBS to be significant if they had local false sign rate (LFSR) less than 0.05 and a positive effect size (corresponding to a positive α_CA_) in at least one cell type in which that gene or TFBS had LFSR < 0.05.

### Testing for positive selection on HARs and HAQERs

To test for positive selection on HAQERs, we intersected the fixed and polymorphic sites with the regions defined as HAQERs from the supplemental material of Mangan et al^32^. We then used the predicted absolute log_2_ fold-change in CA as input to compute alpha, estimating the confidence interval using the bootstrap procedure outlined above. We performed an identical procedure for HARs using the list in the supplemental material of Girskis et al^93^.

To test for enrichment of sites in HARs with large predicted effect in fetal cortical neurons near particular gene sets, we used GREAT^95^, following the procedure outlined above using 0.1 as the upper cutoff and 0.025 as the lower cutoff. We also verified the enrichment for abnormal nervous system electrophysiology and post-translational modification using the permutation procedure outlined above.

### Identifying predicted transcription factor binding sites

We used the JASPAR 2022 core human and mouse transcription factor binding site^102^ (TFBS) files available here: https://jaspar.genereg.net/downloads/. We converted these from LMP to ILO format and then used the matrix_prob function in PWMScan^149^ to create matrix probability files, using the parameter --bg "0.29,0.21,0.21,0.29". We then used the matrix_scan function in PWMScan^149^ to identify TFBS genome-wide using the human, chimpanzee, and gorilla reference genomes and intersected each set of TFBS with the polymorphic and fixed sites.

### Testing for positive selection on transcription factor binding sites

As we identified TFBS in the human reference genome and the human reference genome contains all fixed human-derived substitutions, but only a fraction of the polymorphic sites have the derived allele in the human reference genome, we restricted only to those polymorphic sites where the derived allele was the reference allele when testing for positive selection on TFBS. We additionally restricted to sites in TFBS with p < 10^-5^ for either the human allele or both the chimpanzee and gorilla alleles, so for example if a site was in a TFBS with p < 10^-5^ in chimp but p > 10^-5^ in human and gorilla, that site would not be included. We then used Fisher’s exact test to test for differences between the fixed and polymorphic distributions of predicted effects on CA as described above, using the 60th percentile of the polymorphic distributions as a cutoff in each cell type separately. We then combined information across cell types using MASHR as described above.

To test for enrichment with GREAT^95^, we followed the procedure outlined above using an upper cutoff of 0.25 and a lower cutoff of 0.1 for NFIB and an upper cutoff of 0.1 and a lower cutoff of 0.025 for MEF2A. For permutation testing and testing for positive selection in NFIB binding sites on synaptic genes, we used the genes listed as coding for proteins that localize to the synapse by SYNGO. To determine the set of genes to test for biases in expression in day 100 human-chimp hybrid cortical organoids^116^, we selected all genes that met the following criteria:

1. Had an NFIB binding site containing a substitution assigned to that gene
2. That substitution had an absolute predicted log_2_ fold-change in CA greater than 0.1
3. The sign of the difference in predicted binding affinity matched the sign of the log_2_ fold-change in CA (consistent with the expectation that NFIB opens chromatin^150^.

We then further restricted to genes with FDR < 0.05 in the human-chimp hybrid cortical organoid data and used the binomial test with the background probability equal to the proportion of genes with FDR < 0.05 that have lower expression from the human allele to test for a bias toward higher or lower expression from the human allele in that gene set.

We used the RepeatMasker^135^ annotations for the human genome to assign substitutions to ZNF460 binding sites in Alu elements (i.e., any repeat starting with Alu) and then used Fisher’s exact test to compute the enrichment relative to the number of substitutions in Alus that were not in ZNF460 binding sites. We also used this assignment to Alu or non-Alu to contrast evidence for positive selection in ZNF460 binding sites in and not in Alu elements. To test for positive selection on Alu elements, we used Fisher’s exact test with the cutoff being the 60th percentile of the polymorphic distribution. We then compared this result to the result derived from either all sites not in Alus, or all sites with fewer than 250 species in the alignment (i.e., largely primate-specific) and in a non-Alu repeat.

### Exploring the relationship between positive selection and cell type-specificity

We split a set of sixteen cell types into two groups of eight cell types with diverse developmental origins. We created two groups to ensure that any results replicate when different groups of cell types are used. There were eight cell types in each group as this was the maximum number of cell types we could include in a group without using cell types with highly similar *trans*-regulatory environments. For example, twelve out of the thirty-four cell types we trained ChromBPNet^37^ models for are neuronal and, as they all have similar *trans*-regulatory environments, we could only use two of these (one for each group). The first group consisted of adult cortical microglia, fetal cortical excitatory neurons, adult heart cardiomyocytes, adult kidney proximal tubule cells, fetal heart arterial endothelial cells, fetal chondrocytes, fetal gonad Sertoli cells, and adult cortical astrocytes. The second group consisted of fetal gonad immune cells, adult cortical layer 2/3 intratelencephalic neurons, adult heart smooth muscle cells, adult kidney loop of henle cells, fetal heart pericytes, fetal heart cardiac fibroblasts, fetal gonad pre-granulosa IIa cells, and fetal brain committed oligodendrocyte precursor cells. We performed all analyses on both sets of cell types to ensure that our analyses were reproducible across different cell types. To estimate the cell type-specificity of candidate CRE accessibility, we estimated Tau^151^ and CAE using the predicted counts per million as input for the ancestral and derived alleles and took the average. CAE is based on the previously published metric expression enrichment^152^ (EE) and is defined identically, but uses CA instead of expression level as input. To estimate the cell type-specificity of differential accessibility, we first converted the predicted absolute log_2_ fold-change in CA and the predicted log_2_ fold-change in CA to Z-scores. We then estimated dTau and dCAE^71^ for the absolute log_2_ fold-change as previously described within each group of cell types.

We then tested for stronger positive selection on substitutions with cell type-specific effects in two ways in each cell type. First, we restricted only to the fixed and polymorphic sites with the top 10^6^ largest predicted effects (approximately 8% of the total number of fixed and polymorphic sites). We did this because sites with very low predicted effects tend to have very noisy estimates of cell type-specificity. We then computed confidence intervals for α_CA_ and α_CTS_ using bootstrapping as described above and compared the confidence interval for α_CTS_ to the actual estimate of α_CA_ and vice versa. If the lower bound on the α_CTS_ 95% confidence interval was greater than the actual estimate for α_CA_, we interpreted that as evidence for stronger positive selection on substitutions with cell type-specific effects.

Second, we binned all fixed and polymorphic sites into 8 different bins by dTau with the lowest and highest dTau bin containing 200,000 sites, the next lowest and highest bins containing 200,000 sites, the next bins containing 200,000 sites, and the last (middle) two bins containing the remaining sites. We then estimated a confidence interval for α_CA_ in each bin down-sampling to the minimum number of fixed and polymorphic sites across all bins before doing so. To further ensure that our results were robust to differences in the cell types used, we iteratively removed the cell type with the largest α_CA_ in the most cell type-specific bin for two iterations and recomputed dTau and confidence intervals.

To test for enrichment of cell type-specific effects near genes for fetal cortical excitatory neurons and fetal gonad pre-granulosa cells, we first filtered to sites with predicted absolute log fold-change in CA greater than 0.1. We then split sites into those with dTau > 0.9 (highly cell type-specific effects, foreground) and dTau < 0.5 (broadly shared effects, background). We then used GREAT and the permutation-based method to test for enrichment as outlined above.

To test for differences in how fixed and polymorphic substitutions affect the cell type-specificity of *cis*-regulatory elements, we computed the difference in CAE between the derived and ancestral allele. Notably, this does not relate directly to the predicted log_2_ fold-change in CA between the two alleles. For example, a substitution with a predicted log_2_ fold-change in CA of 0 in one cell type and -1 in all others would have a very large difference in CAE because the substitution would make that CRE more specific to that one cell type even if it did not affect CA in that cell type. We then restricted to only sites with CAE > 0.3 to enrich for sites that are in candidate CREs in the cell type of interest, and then used Fisher’s exact test with the cutoff being the 90th percentile of the polymorphic CAE distribution. We repeated this analysis after swapping the adult neurons with the fetal neurons and reran the test to ensure that the observed difference wasn’t due to which cell types were included in groups 1 and 2. We also swapped fetal cortical neurons for fetal dopaminergic neurons or fetal purkinje neurons and adult cortical neurons for adult cortical VIP inhibitory neurons or adult D1 medium spiny neurons and reran the test. Finally, we tested all combinations of the following parameters: CAE cutoff of 0.3, 0.4, or 0.5, using the 60th, 70th, 80th, and 95th percentile of the polymorphic distribution as the cutoff, removing repeats and pseudogenes, and restricting to sites with at least 250 species in the multiway alignment. We then took the average rank of each cell type (1 being the most evidence for fixed sites having higher difference in CAE, 20 being the least evidence) across all parameter combinations to ensure that our results were consistent across various analysis decisions.

### Identifying and analyzing RAGs and RALs

For the chimpanzee input, we filtered sites in the same way as for human-derived sites, with the exception that we did not remove sites that are polymorphic in chimpanzees beyond those that were filtered out as described in Starr et al.^153^ (which is why there are more chimpanzee input sites). We did not remove sites that were polymorphic in chimpanzee because publicly available datasets of chimpanzee polymorphism were generated by aligning reads to the human genome and we reasoned that this could lead to disproportionate false negative in highly diverged regions of the genome potentially leading to considerable biases^154^. We still removed polymorphisms from the human data because hRAGs/hRALs that are not driven by polymorphic substitutions are the most likely to contribute to human-chimp divergence and be promising for future follow-up experiments. Moreover, the strongly similar properties of the hRAGs/hRALs compared to the cRAGs/cRALs suggests that this decision does not bias our results to a detectable extent.

To identify RAGs and RALs, we used all the cell types in the two groups outlined above first restricted only to sites with an absolute difference in CAE greater than 0.025 (between 1,000 and 400,000 sites depending on the cell type). We then iterated through the list of sites and took all sites in a 1 kilobase window around each central site. If there were at least 3 sites with absolute difference in CAE greater than 0.025 and the same sign and 0 sites with absolute difference in CAE greater than 0.025 and the opposite sign, we called that as a RAG or RAL depending on the sign. To ensure a fair comparison between human and chimpanzee, we identified hRAGs, hRALs, cRAGs, and cRALs using human-referenced predictions in which the human reference allele was changed to either the ancestral allele (for human-derived sites) or the derived allele (for chimp-derived sites). We then also identified hRAGs, hRALs, cRAGs, and cRALs using chimp-referenced predictions in which the chimp reference allele was changed to either the ancestral allele (for chimp-derived sites) or the derived allele (for human-derived sites). For hRAGs, hRALs, cRAGs, and cRALs independently, we then merged any overlapping elements (which occur somewhat frequently due to initially calling elements using human-referenced and chimp-referenced predictions and rarely due to being called in multiple cell types. The latter occurs primarily because elements were called separately in cell type groups one and two. For example, several elements were called in both fetal cortical excitatory neurons and adult neocortical L2/3 intratelencephalic neurons because these cell types are somewhat similar and in different groups). Elements are then identified by the list of sites contributing to them being called as well as the cell type(s) they were called in.

To assign p-values to each RAG, we used the genome-wide probability that a substitution that is polymorphic (0.25 < DAF < 0.75) in humans increases or decreases the cell type-specificity of CA beyond the cutoff used (difference in CA greater than or less than 0.025 for RAGs and RALs respectively) as a reasonable neutral null model. We used the binomial test comparing the number of qualifying sites in each RAG/RAL to the total number of fixed sites with the background polymorphic probability as the probability for the binomial test to compute p-values for each RAG and RAL in each cell type. All else equal, RAGs/RALs in cell types with more qualifying input sites have less significant p-values. Due to the discrete nature of the test and that most RAGs and RALs have exactly 3 qualifying substitutions, any p-value cutoff used to filter RAGs or RALs for enrichment analyses would lead to an artificial depletion of RAGs and RALs in those cell types with more qualifying input sites. For example, using a p-value cutoff of 10^-4^ reduces the number of hRAGs called in adult cortical ITL23 neurons from 915 to 569, but only reduces the number in fetal cortical excitatory neurons from 517 to 504. Although ultimately the cutoff for “high-confidence” is somewhat arbitrary, we selected 5×10^-7^ as the cutoff for high-confidence RAGs and RALs as there are approximately 10^7^ human-derived fixed sites in our dataset after filtering (to be exact) with an average of five sites per one kilobase window. The latter suggests that not all tests are independent and given this non-independence we reasoned that 1/(10^7^/5) = 5×10^-7^ is a reasonable cutoff.

To validate RAGs and RALs, we focused on cell type group one which contains fetal cortical excitatory neurons. The human-chimp hybrid glutamatergic neuron ATAC-seq data were shared with us by the authors^123^. We restricted only to ATAC peaks with FDR < 0.05 for allele-specific chromatin accessibility. We then intersected that set of peaks with the list of RAGs and RALs and computed the number of peaks that agree in sign with the RAGs and the number that disagree. For hRAGs, the vast majority of sites increase accessibility specifically in fetal neurons and the remaining few decrease accessibility in other cell types while having no effect in fetal neurons. However, it is worth noting that these hRAGs do not take into account chimpanzee-derived substitutions which could lead to higher accessibility of the chimp allele or human-derived substitutions that decrease accessibility across all cell types. Overall, if hRAGs are accurately called, we would expect most of them to have higher accessibility from the human allele and so have a positive log_2_ fold-change in the human-chimp hybrid neuron ATAC data and count those as agreeing in sign. We repeated this logic for hRALs, cRALs, and cRAGs and summed across element types. We note also that this is likely an underestimate as to the true agreement as, for example, a cRAG could also contain a human-derived substitution that strongly increases accessibility in fetal neurons leading to disagreement in sign with experimental data. We would not expect consistent agreement in sign for elements called in any of the other eleven cell types. Therefore, for each of those cell types the number of agreeing and disagreeing ATAC peaks as described above. We then compared the number of agree/disagree peaks for fetal neuron elements to that for other cell types using Fisher’s exact test.

To test for positive selection on conserved sites in neuronal hRAGs and hRALs, we restricted to hRAGs and hRALs called in fetal cortical excitatory neurons, adult neocortical L2/3 intratelencephalic neurons, or both (the only types of neuron included) and expanded to a one kilobase window around the site with the median position. We then restricted to fixed and polymorphic sites in this window and tested for positive selection using the 90th percentile of the polymorphic PhyloP distribution as the cutoff. We next intersected the list of hRAGs and hRALs with the lists of HARs and HAQERs and counted the number of unique intersecting elements. We tested for enrichment of neuronal (or non-neuronal) hRAGs and hRALs in (or not in) HARs and HAQERs with Fisher’s exact test.

To test for enrichment of each type of element near specific genes using hRAGs as an example, we compared the number of hRAGs assigned (or not assigned) to a gene with the number of sites with difference in CAE > 0.025 assigned (or not assigned) to a gene using a one-sided Fisher’s exact test, restricting only to genes with at least 250 sites with difference in CAE > 0.025 and then FDR-corrected the p-values. To test for enrichments of neuronal hRAGs/hRALs relative to cRAGs/cRALs and vice versa using hRAGs as an example, we input the positions of hRAGs to GREAT as the foreground and the positions of hRAGs and cRAGs as the background. We considered any category with FDR < 0.05 to be significant. We then restricted to genes with FDR < 0.05 when comparing human expression to chimp expression in DLPFC L2/3 IT neurons and used the binomial test with background probability equal to the proportion of genes with FDR < 0.05 that have lower expression in human to test for a bias toward higher or lower expression in human for the genes in the divalent inorganic cation transmembrane transporter activity that have at least one hRAG assigned to them.

### Analysis of public gene expression datasets

The DLPFC snRNA-seq data^92^ was processed as previously described^113^. The human-chimp hybrid cortical organoid data was processed as previously described^153^. The human-chimp hybrid data from six iPSC-derived cell types was processed as previously described^71^.

### Quantification of gene desert size and neuronal wiring

To quantify gene desert size, we computed the distance between the transcription start sites^140^ of the two genes flanking each gene and assigned that distance to the central gene. We removed all genes directly adjacent to centromeres as this leads to inaccurate estimation of gene desert size. We then restricted to the 100 genes with the largest gene desert size. We considered a gene to be involved in neuronal wiring if it’s protein product localized to the cell surface^155^ and it was part of a protein family known to be involved in neuronal wiring (e.g., protocadherins, cadherins, latrophilins, teneurins, ROBO genes). We considered a gene to be a transcriptional regulator if it was a transcription factor or co-factor.

### Statistical analysis

All statistical tests were performed using scipy^156^ v1.14.1 except that MASHR was used as described above and confidence intervals for Fisher’s exact test were computed with the fisher.test function in R. The Benjamini-Hochberg^157^ method was always used to compute the false discovery rate.

## Acknowledgements

We thank Liqun Luo and other Luo lab members for helpful discussion. We also thank Leslie Magtanong and other members of the Fraser Lab for helpful discussions and feedback on the manuscript. Some subfigures were made with biorender. Model training and predictions were primarily run on the Stanford Marlowe cluster and we thank Stanford Research Computing for the compute time and their assistance. We thank Janet Song for sharing the human-chimp hybrid excitatory neuron ATAC data ahead of publication.

## Authors contributions

MEP trained ChromBPNet models and performed inference of substitution effects on chromatin accessibility. JG performed the electrophysiological characterization of human and chimpanzee SUR2B and analyzed the electrophysiology data. ALS performed all other bioinformatic analysis, visualization, validation, and writing of software with guidance from HBF. ALS and HBF wrote the main text and ALS created the figures with input from HBF. All authors edited and approved the manuscript. HBF and CGN provided funding for the study.

## Data availability

The DLPFC snRNA-seq data is available from https://data.nemoarchive.org/biccn/grant/u01_sestan/sestan/transcriptome/sncell/10x_v3/. The constraint metrics pLI and missense depletion as well as the polymorphism data was downloaded from https://gnomad.broadinstitute.org. The SFARI ASD-linked genes were downloaded from https://gene.sfari.org/. The data from the study of human-chimpanzee hybrid cortical organoids is available from the gene expression omnibus (GEO) accession GSE144825. The hybrid expression data from six cell types is available from GEO with accession GSE232949. Transcription factor position-weight matrices are available from https://jaspar.genereg.net/downloads/. The repeat masker data is available at https://hgdownload.soe.ucsc.edu/goldenPath/hg38/database/. The low complexity region annotations are available at: https://github.com/genome-in-a-bottle/genome-stratifications/blob/master/GRCh38/LowComplexity. The repeat annotations are available at https://repeatbrowser.ucsc.edu/data/.

The *Mus musculus* polymorphism data is available at http://wwwuser.gwdg.de/~evolbio/evolgen/wildmouse/.

The VCF file from He et al.^67^ kindly shared by the authors. The *Pteropus Alecto* genome annotation is available at: https://genome.senckenberg.de/download/TOGA/human_hg38_reference/Chiroptera/Pteropus_alecto__black_flying_fox__pteAle1/. The list and locations of UCNEs are available at: https://epd.expasy.org/ucnebase.

The adult cortical microglia ATAC data are available in the Synapse repository with accession syn26207321. The fetal chondrocyte ATAC data are available from GEO accession GSE214394 and GSE122877. The fetal cortical excitatory neuron and neural progenitor ATAC data are available from dbGaP accession phs001958.v1.p1. The fetal heart ATAC data are available from GEO accession GSE181346. The adult kidney ATAC data are available from GEO accession GSE262931. The adult heart ATAC data are available at http://ns104190.ip-147-135-44.us/data_CARE_portal/snATAC/bedfiles. The fetal brain ATAC data Is available at https://storage.googleapis.com/linnarsson-lab-human/ATAC_dev/10X and the sample and barcode to cell type mapping is available at https://github.com/linnarsson-lab/fetal_brain_multiomics. The adult brain ATAC data is available from http://catlas.org/catlas_downloads/humanbrain/bedpe and the peaks from http://catlas.org/catlas_downloads/humanbrain/cCREs. The fetal gonad ATAC data is available downloaded from the BioStudies repository accession E-MTAB-10570 and the metadata mapping sample and barcode to cell type was shared with us by the authors.

ChromBPNet model weights have been deposited at: http://mepalmer.net/34_models.tgz.

## Code availability

All code needed to reproduce the analyses described in this study is available at https://github.com/astarr97/PosSelect. ChromBPNet variant effect prediction code is available here: https://github.com/mepster/variant-scorer/tree/mepalmer_multigpu.

## Materials and correspondence

Correspondence should be addressed to HBF at and ALS at. Requests for materials will be fulfilled by the corresponding authors.

## List of supplementary materials

Supplemental Figures 1-24

Supplemental Tables 1-6

Supplemental Table 1 contains the results of testing for positive selection on nonsynonymous sites in human phenotype ontology gene sets using PhyloP.

Supplemental Table 2 contains the results of testing for positive selection on non-coding sites in human phenotype ontology gene sets using PhyloP.

Supplemental Table 3 contains the results of testing for positive selection on non-coding sites for individual genes using ChromBPNet predictions.

Supplemental Table 4 contains the results of testing for positive selection on non-coding sites for the binding sites of transcription factors using ChromBPNet predictions.

Supplemental Table 5 contains the results for hRAGs and hRALs. Supplemental Table 6 contains the results for cRAGs and cRALs.

Supplemental Texts 1-6

Supplemental Text 1 is on validating our framework to detect positive selection using simulations

Supplemental Text 2 is on the agreement of our results with previously published results.

Supplemental Text 3 discusses various sources of false positives and false negatives for our framework as well as the MK test.

Supplemental Text 4 discusses positive selection on nonsynonymous substitutions in genes that regulate neuron projection (i.e., axon, dendrite, and synapse) development

Supplemental Text 5 discusses potential sources of disagreement between testing for positive selection on nonsynonymous sites using our framework with PhyloP scores and the MK test.

Supplemental Text 6 discusses the sensitivity of our results to different parameter choices.

