## Supplemental Figures for "A framework to detect positive selection using variant effect predictions reveals widespread adaptive evolution of human neurons"

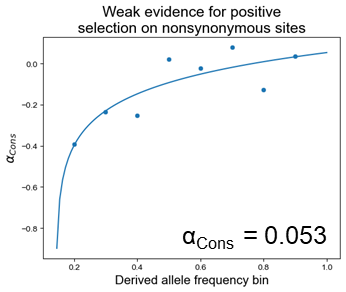

**Supplemental figure 1, related to figure 1:** Result from asymptotic comparison of conservation score distribution for fixed and polymorphic substitutions. The points are positioned at the upper limit of each derived allele frequency bin: 0.1-0.2, 0.2-0.3, …, 0.8-0.9.

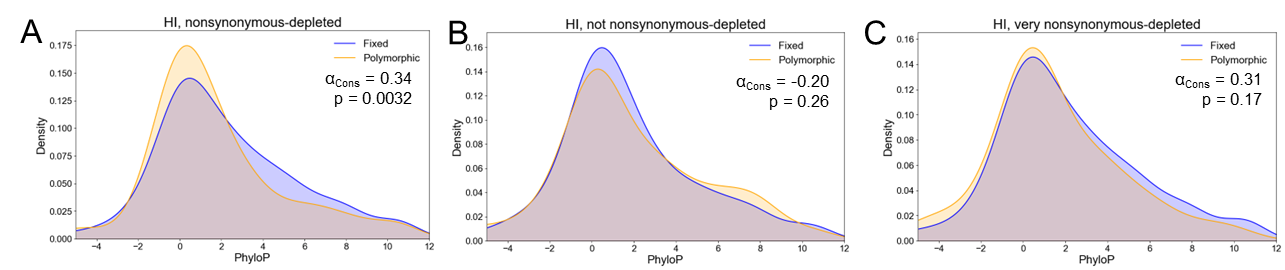

**Supplemental figure 2, related to figure 1: A)** Distribution of conservation scores for fixed and polymorphic substitutions in haploinsufficient genes that are depleted of nonsynonymous variants (i.e. Z-score > 2). **B)** Distribution of conservation scores for fixed and polymorphic substitutions in haploinsufficient genes that are not depleted of nonsynonymous variants. **C)** Distribution of conservation scores for fixed and polymorphic substitutions in haploinsufficient genes that are highly depleted (i.e. Z-score > 4) of nonsynonymous variants.

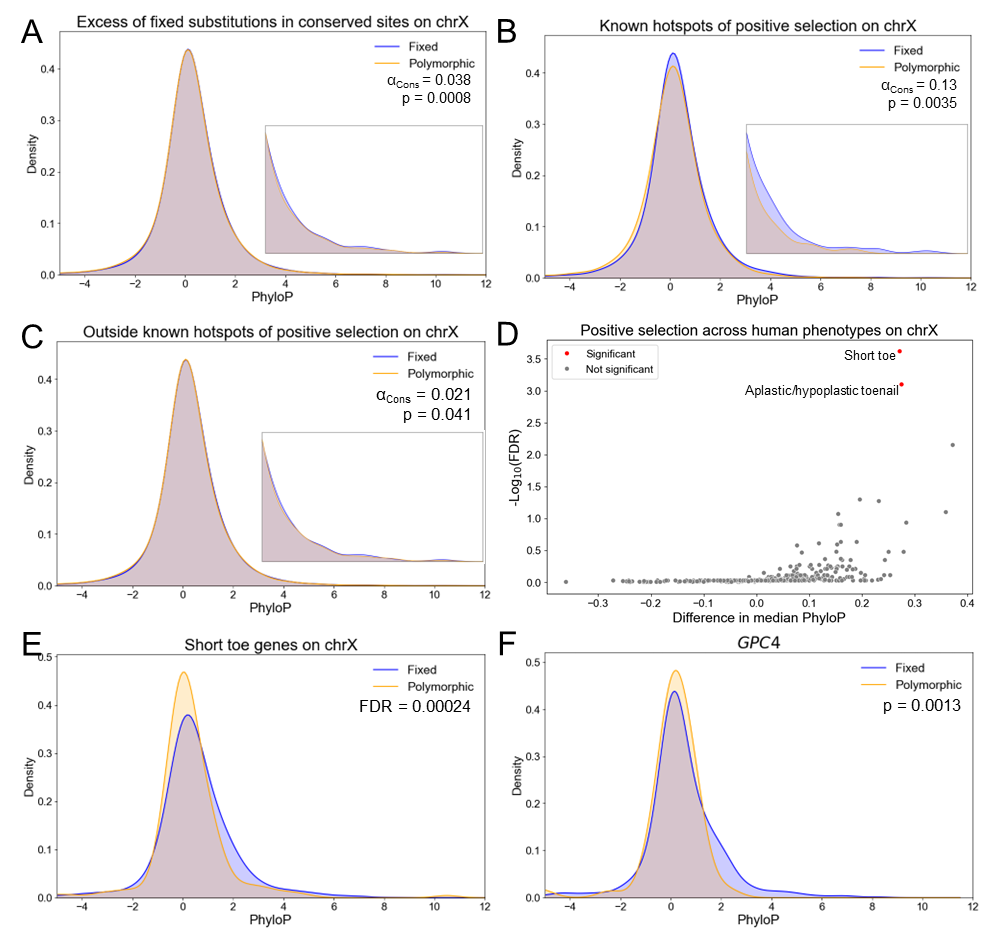

**Supplemental figure 3, related to figure 2**: **A)** Distribution of PhyloP scores for non-coding fixed and polymorphic sites on the X chromosome. The inset shows the distribution magnified in the range 3-12. For A-E, the cutoff for computing α_Cons_ is the 60^th^ percentile of the polymorphic conservation score distribution. **B)** Same as in (A) but for sites on the X chromosome near multi-copy testes-specific genes or in regions of known selective sweeps. **C)** Same as in (A) but for sites on the X chromosome not near multi-copy testes-specific genes or in regions of known selective sweeps. **D)** Volcano plot showing positive selection on conserved non-coding sites on the X chromosome across human phenotypes. Gene sets were considered significant if they had Mann-Whitney U test FDR < 0.05 and Fisher’s exact test nominal p < 0.05, with the cutoff being the 90^th^ percentile of the polymorphic conservation score distribution. **E)** Same as in (A) but only for sites near genes in the “Short toe” gene set. **F)** Same as in (A) but only for sites near *GPC4*. P-value is from one-sided Fisher’s exact test.

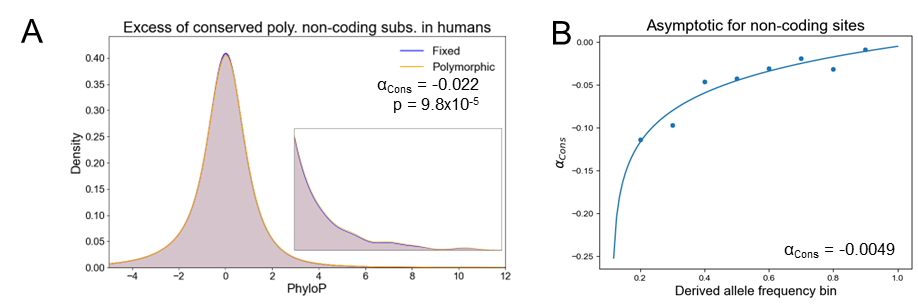

**Supplemental figure 4, related to figure 2: A)** Distribution of PhyloP scores for non-coding fixed and polymorphic sites genome-wide. The inset shows the distribution magnified in the range 3-12. The cutoff for computing α_Cons_ is the 95^th^ percentile of the polymorphic conservation score distribution. **B)** Result from asymptotic comparison of conservation score distribution for non-coding fixed and polymorphic sites. The point are positioned at the upper limit of each derived allele frequency bin: 0.1-0.2, 0.2-0.3, …, 0.8-0.9.

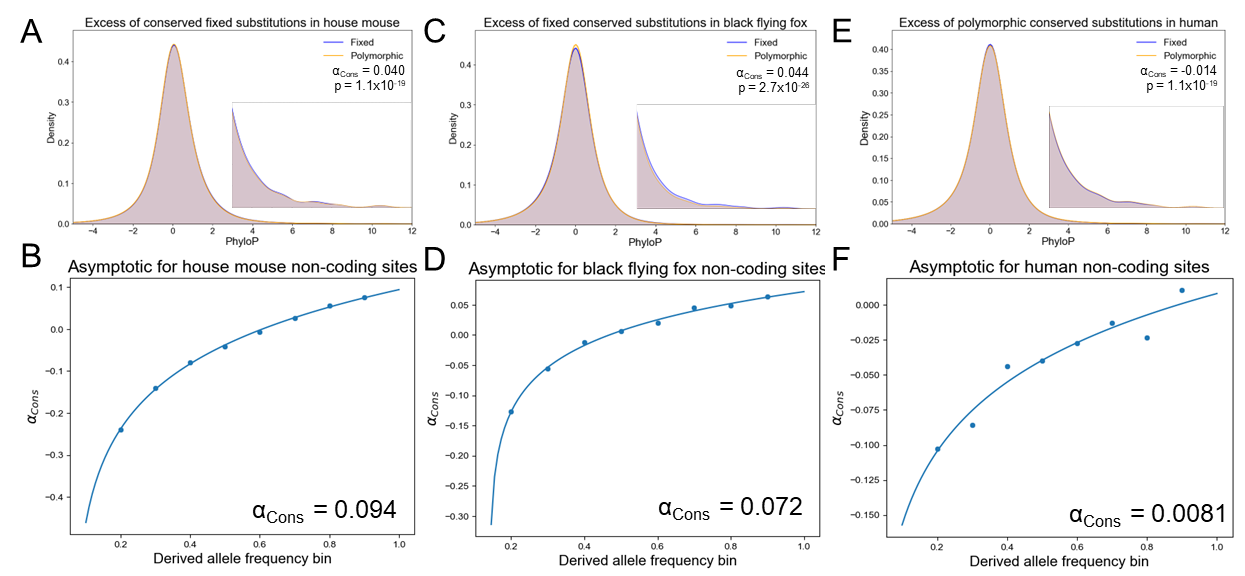

**Supplemental Figure 5, related to figure 2: A)** Distribution of PhyloP scores for non-coding fixed and polymorphic sites genome-wide for house mouse (*Mus musculus domesticus*)*.* The inset shows the distribution magnified in the range 3-12. The cutoff for computing α_Cons_ is the 95^th^ percentile of the polymorphic conservation score distribution. **B)** Result from asymptotic comparison of conservation score distribution for non-coding fixed and polymorphic sites genome-wide for house mouse. The point are positioned at the upper limit of each derived allele frequency bin: 0.1-0.2, 0.2-0.3, …, 0.8-0.9. **C)** Same as in (A) but for black flying fox (*Pteropus alecto*). **D)** Same as in (B) but for black flying fox. **E)** Same as in (A) but for human (*Homo sapiens*) using equivalent parameters to A-D. **F)** Same as in (B) but for human.

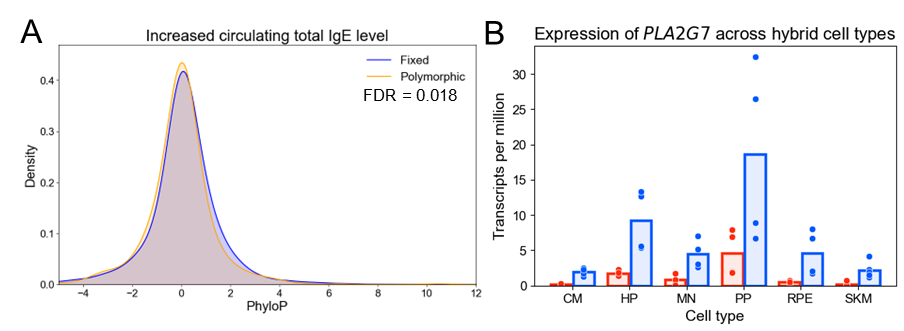

**Supplemental Figure 6, related to figure 2: A)** Distribution of PhyloP scores for non-coding fixed and polymorphic sites near genes in the “Increased circulating total IgE level” gene set*.* Mann-Whitney U test FDR is shown. **B)** Allele-specific expression of *TMPRSS2* across six human/chimpanzee hybrid cell types (CM = cardiomyocytes, HP = hepatic progenitors, MN = motor neurons, PP = pancreatic progenitors, RPE = retinal pigmented epithelial cells, SKM = skeletal muscle). FDR < 0.05 in all cell types. Red = human allele, blue = chimpanzee allele.

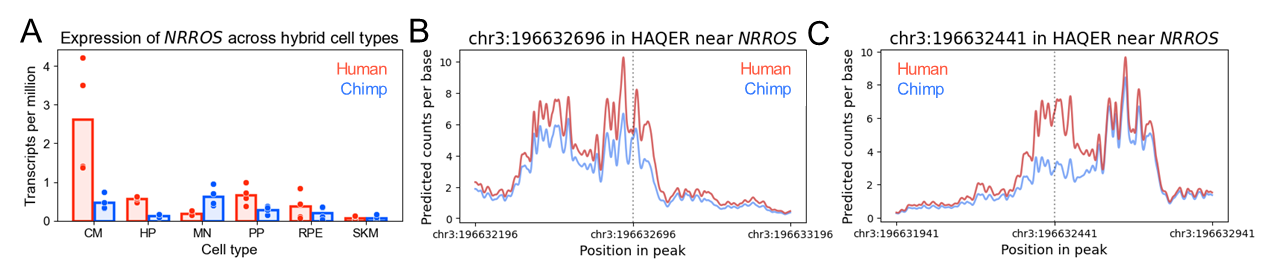

**Supplemental Figure 7, related to figure 4: A)** Allele-specific expression of *NRROS* across six human/chimpanzee hybrid cell types (CM = cardiomyocytes, HP = hepatic progenitors, MN = motor neurons, PP = pancreatic progenitors, RPE = retinal pigmented epithelial cells, SKM = skeletal muscle). FDR < 0.05 in CM, MN, HP, and PP. **B)** Predicted effect of a human-specific substitution at chr3:196632696 in a HAQER near *NRROS* on chromatin accessibility in adult ventricular cardiomyocytes. **C)** Same as in (B) but for chr3:196632441.

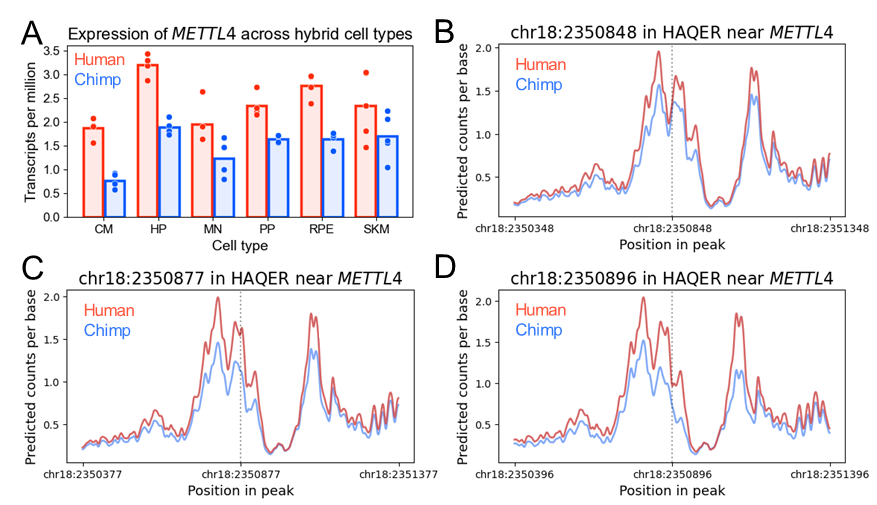

**Supplemental Figure 8, related to figure 4: A)** Allele-specific expression of *METTL4* across six human/chimpanzee hybrid cell types (CM = cardiomyocytes, HP = hepatic progenitors, MN = motor neurons, PP = pancreatic progenitors, RPE = retinal pigmented epithelial cells, SKM = skeletal muscle). FDR < 0.05 in all cell types. **B)** Predicted effect of a human-specific substitution at chr18:2350848 in a HAQER near *METTL4* on chromatin accessibility in adult ventricular cardiomyocytes. **C)** Same as in (B) but for chr18:2350877. **D)** Same as in (B) but for chr18:2350896.

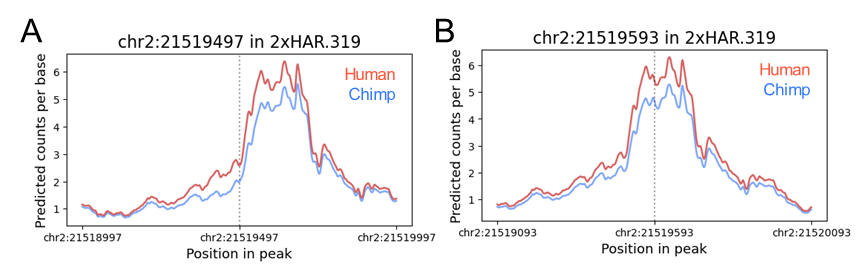

**Supplemental Figure 9, related to figure 4: A)** Predicted effect of a human-specific substitution at chr2:21519497 on chromatin accessibility in 2xHAR.319 in fetal cortical excitatory neurons. **B)** Same as in (A) but for chr2:21519593.

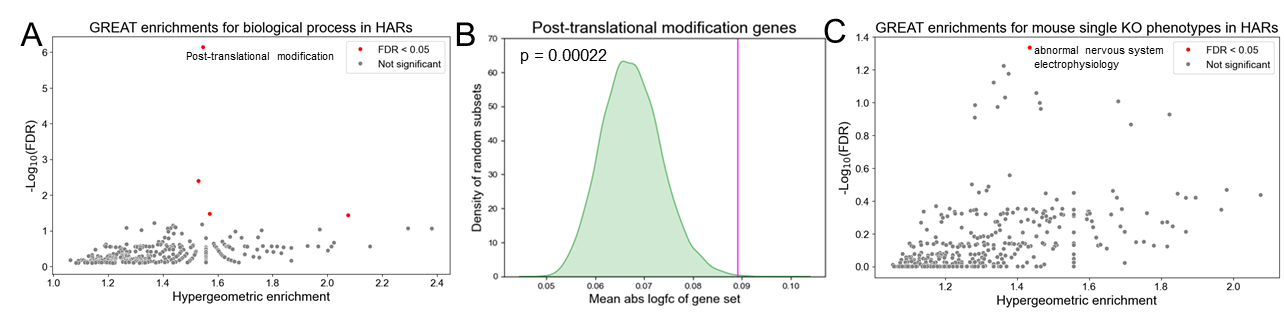

**Supplemental figure 10, related to figure 4: A)** Volcano plot of GREAT enrichment for high predicted absolute log_2_ fold-change in CA for GO biological process categories. The fold enrichment for each gene and corresponding local false sign rate are shown on the x- and y-axes respectively. **B)** Green distribution shows the expected mean predicted absolute log_2_ fold-change based on 10,000 bootstraps for predicted log_2_ fold-change in CA for substitutions in HARs near genes in the GO category post-translational. Magenta line shows the observed mean for substitutions in HARs near genes in the GO category “post-translational”. **C)** Same as in (A) but for mouse single gene KO phenotype.

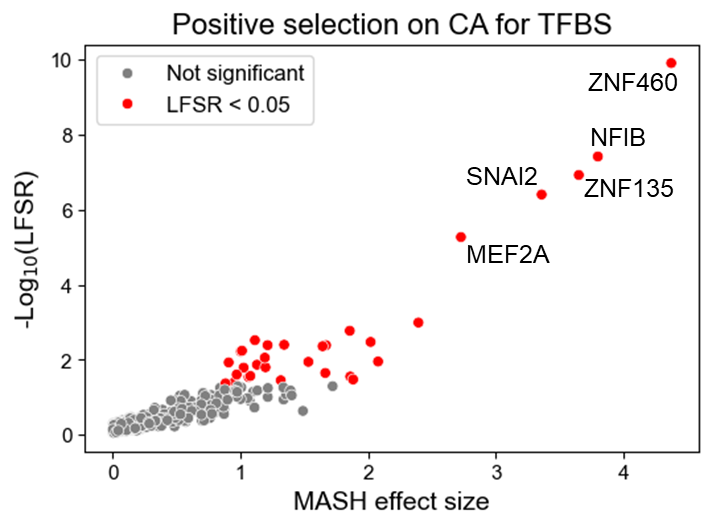

**Supplemental figure 11, related to figure 5:** Volcano plot of positive selection on CA for binding sites of individual transcription factors. Only TFBS with positive MASH effect size are shown. The maximum effect size for each gene across cell types and corresponding local false sign rate are shown on the x- and y-axes respectively.

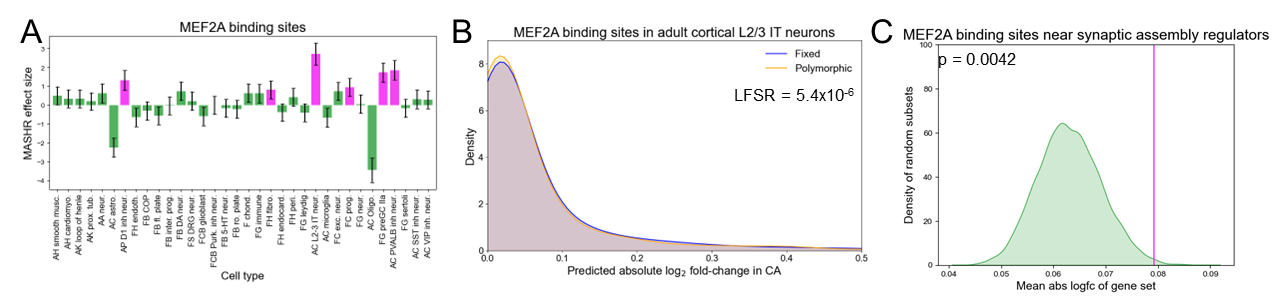

**Supplemental figure 12, related to figure 5: A)** Positive selection on MEF2A binding sites across cell types. Bar height shows the estimated effect size from MASHR; the error bar is the posterior standard deviation from MASHR. Magenta indicates local false sign rate (LFSR) < 0.05. In all cases, the cutoff for computing Fisher’s exact test is the 60^th^ percentile of the polymorphic absolute predicted log_2_ fold-change in CA distribution. **B)** Distribution of predicted effects on CA for fixed and polymorphic substitutions in MEF2A binding sites in layer 2/3 intratelencephalic neurons. **C)** Green distribution shows the expected mean absolute log fold-change based on 10,000 bootstraps for predicted log_2_ fold-change in CA for substitutions in MEF2A binding sites near synaptic assembly regulators. Magenta line shows the observed mean for synaptic assembly regulators.

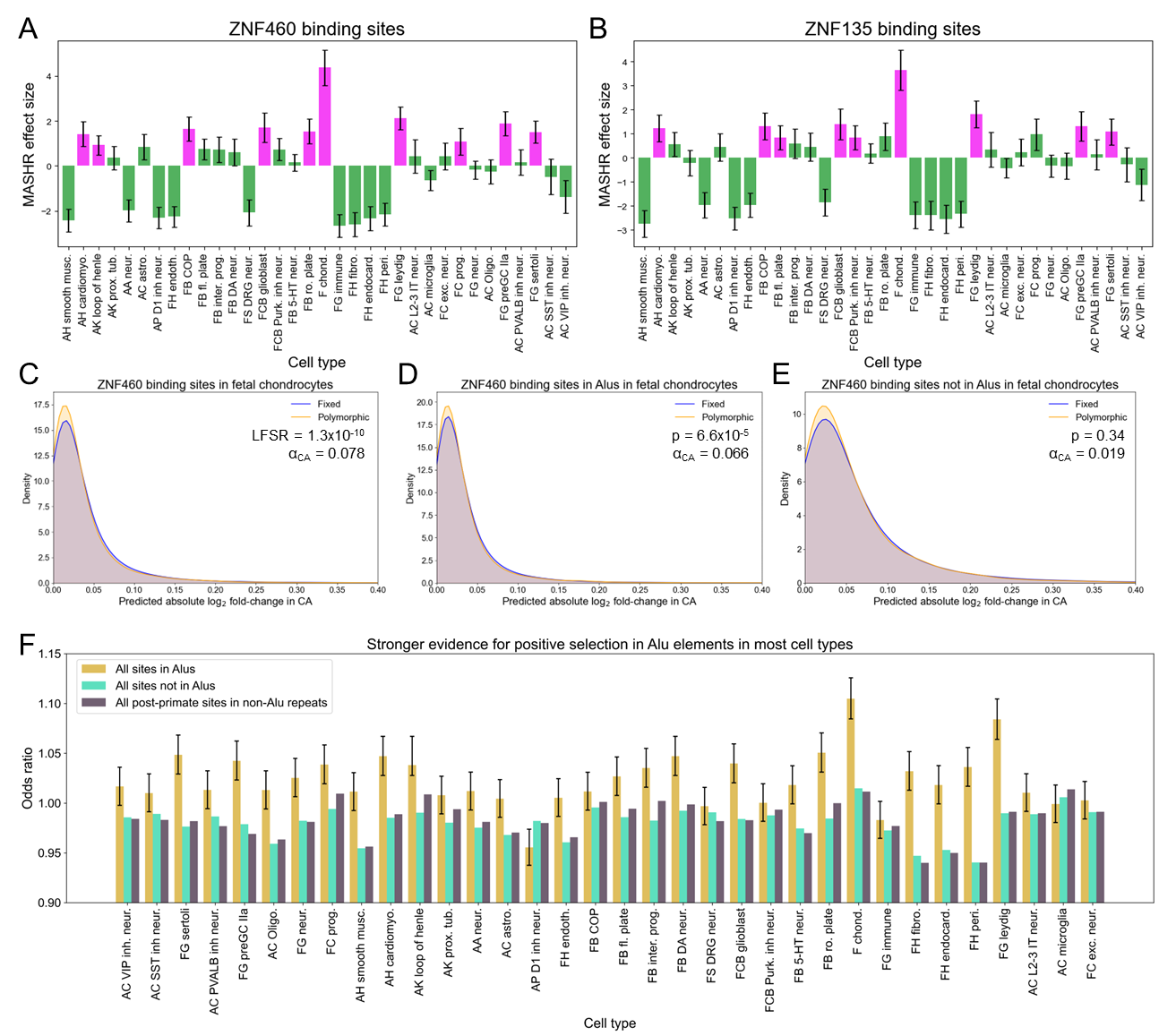

**Supplemental figure 13, related to figure 5: A)** Positive selection on ZNF460 binding sites across cell types. Bar height shows the estimated effect size from MASHR, the error bar is the posterior standard deviation from MASHR. Magenta indicates local false sign rate (LFSR) < 0.05. In all cases, the cutoff for computing Fisher’s exact test is the 60^th^ percentile of the polymorphic absolute predicted log_2_ fold-change in CA distribution. **B)** Same as in (A) but for ZNF135. **C)** Distribution of predicted absolute log_2_ fold-change in CA for fixed and polymorphic substitutions in ZNF460 binding sites in fetal chondrocytes. **D)** Same as in (C) but for ZNF460 binding sites in Alu elements. **E)** Same as in (C) but for ZNF460 binding sites not in Alu elements. **F)** Bar plot of Fisher’s exact odds ratio for sites in Alus, all sites genome-wide, and sites in repetitive elements with fewer than 250 species in the alignment (largely primate-specific). The error bar shows the 95% confidence interval for the odds ratio.

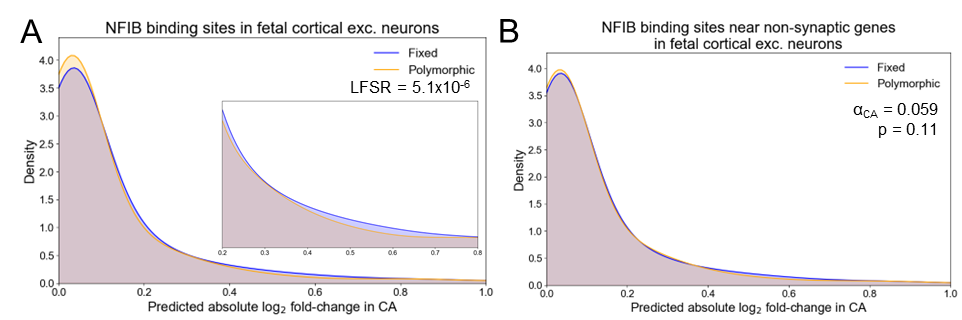

**Supplemental figure 14, related to figure 5: A)**. Distribution of predicted absolute log_2_ fold-change in CA for fixed and polymorphic substitutions in NFIB binding sites in fetal cortical excitatory neurons. The inset shows the distribution magnified in the range 0.2-0.8. **B)** Same as in (A) but for NFIB binding sites not near synaptic genes.

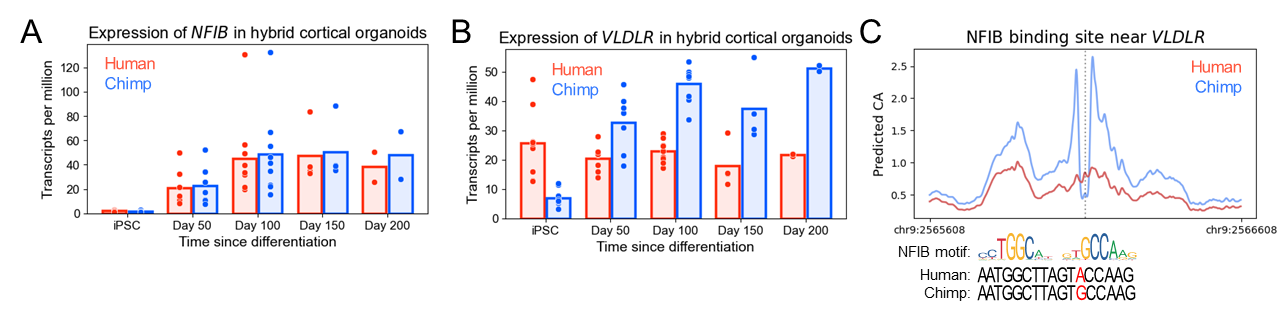

**Supplemental figure 15, related to figure 5: A)** Allele-specific expression of *NFIB* across a time course of human/chimp hybrid cortical organoid development. **B)** Same as in (A) but for *VLDLR*. **C)** Predicted effect of fixed substitution at chr9:2566108 in a candidate CRE near *VLDLR* on chromatin accessibility in fetal cortical excitatory neurons. The smoothed predicted counts per base are shown. The NFIB binding motif is shown below the plot along with the human and chimp sequence context for chr9:2566108.

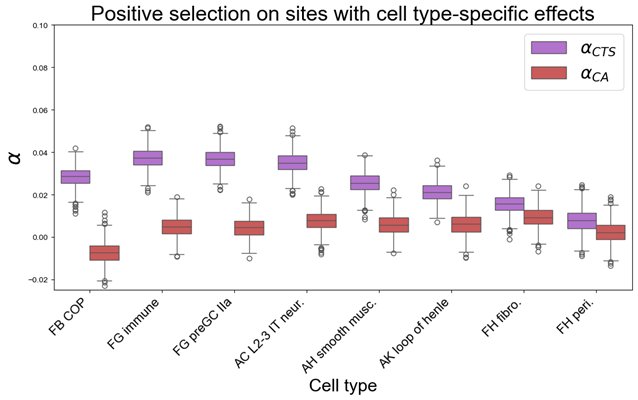

**Supplemental figure 16, related to figure 6:** Box plot comparing α_CTS_ and α_CA_ across cell types based on 1,000 bootstraps. The cutoff for computing α is the 60^th^ percentile of the polymorphic score distribution.

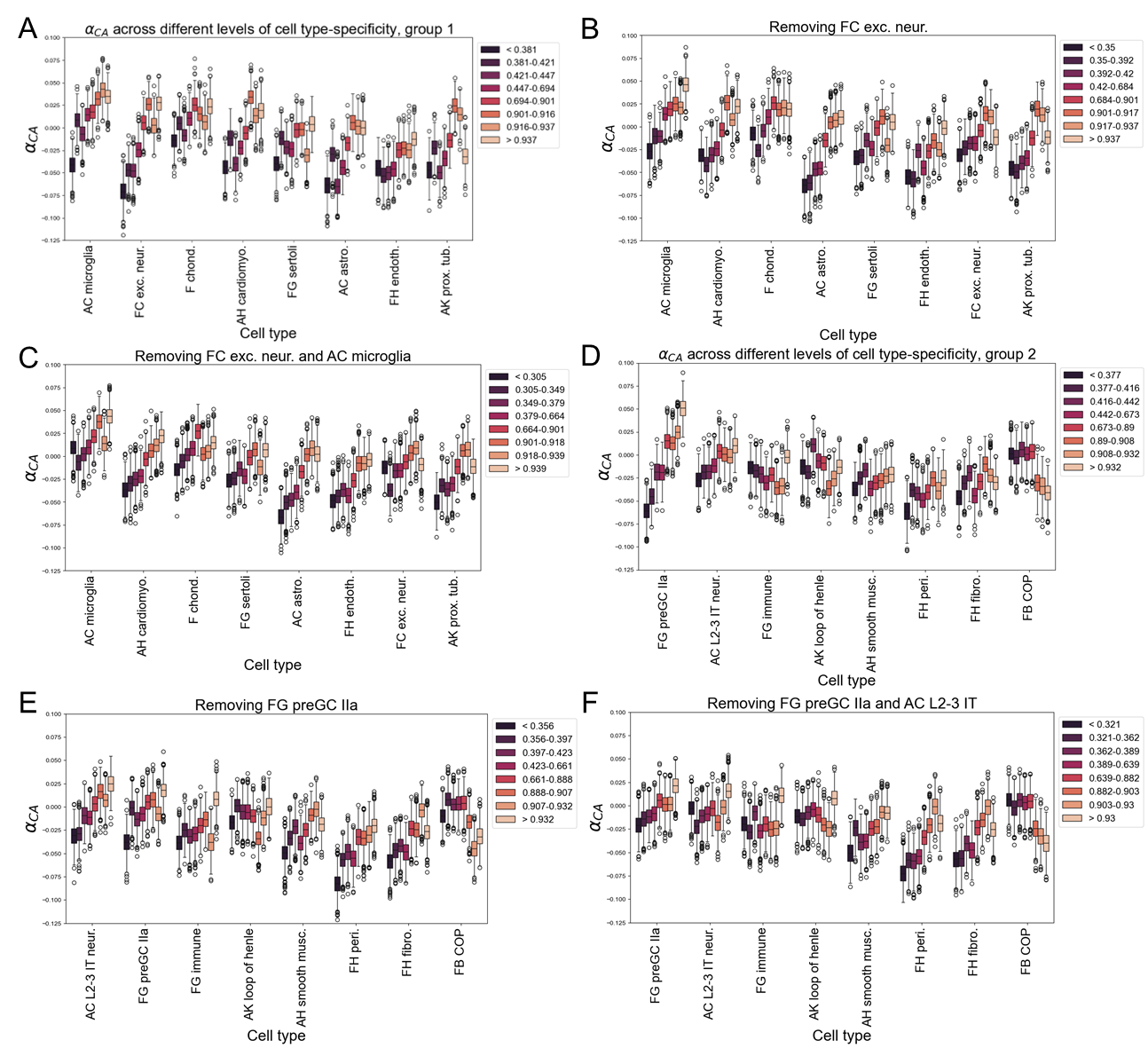

**Supplemental figure 17, related to figure 6: A)** Box plot comparing α_CA_ across different dTau bins based on 1,000 bootstraps for 8 different cell types. The colors indicate the dTau bins, with the lightest color being the bin with the most cell type-specific substitutions. **B)** Same as in (A) but after removing fetal cortical excitatory neurons from the computation of dTau. **C)** Same as in (B) but with the additional removal of microglia. **D)** Same as in (A) but for the second group of eight cell types. **E)** Same as in (D) but after removing preGC IIa cells. **F)** Same as in (E) but after the additional removal of L2-3 IT neurons.

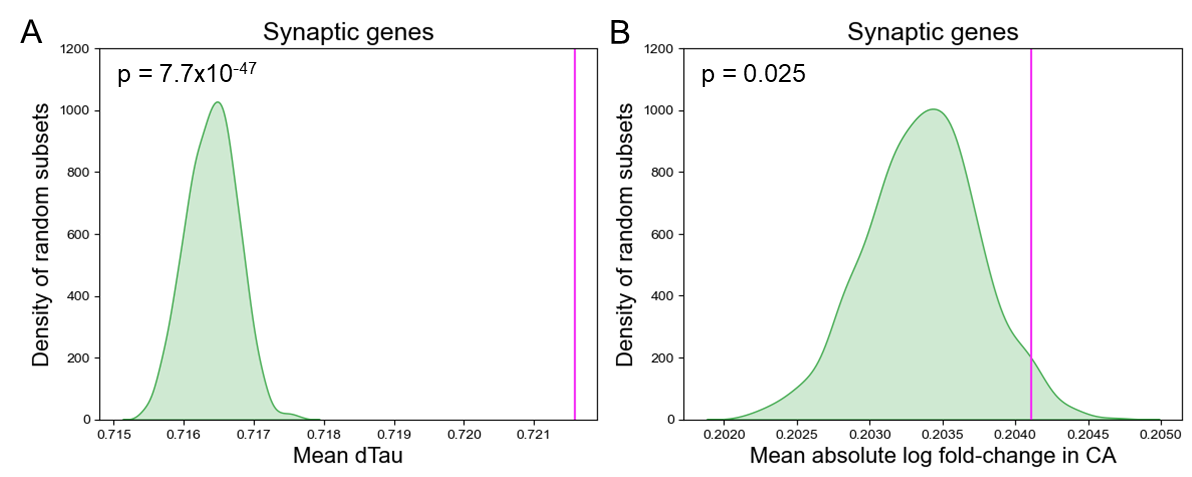

**Supplemental figure 18, related to figure 6: A)** Green distribution shows the expected mean dTau based on 1,000 bootstraps for substitutions with predicted absolute log_2_ fold-change greater than 0.1 near synaptic genes. Magenta line shows the observed mean for synaptic genes. **B)** Same as in (A) but for mean predicted absolute log_2_ fold-change in CA.

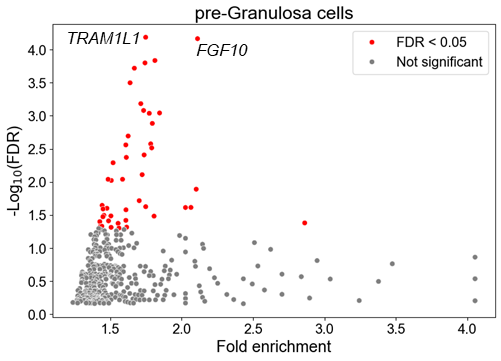

**Supplemental figure 19, related to figure 6:** Volcano plot of GREAT enrichment for high dTau per gene in pre-Granulosa IIa cells. The fold enrichment for each gene and corresponding local false sign rate are shown on the x- and y-axes respectively.

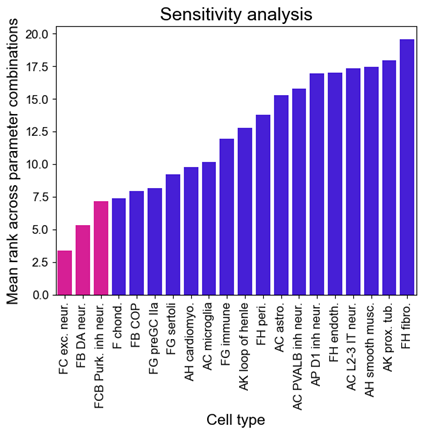

**Supplemental figure 20, related to figure 6:** Mean rank of each cell type across all combinations of different parameter choices (Methods). The height of the bars represents the average rank of each cell type (rank = 1 being the most evidence for fixed sites having higher difference in CAE, rank = 20 being the least evidence) across all parameter combinations to ensure that our results were consistent across various analysis decisions. The three fetal neuronal cell types are shown in magenta.

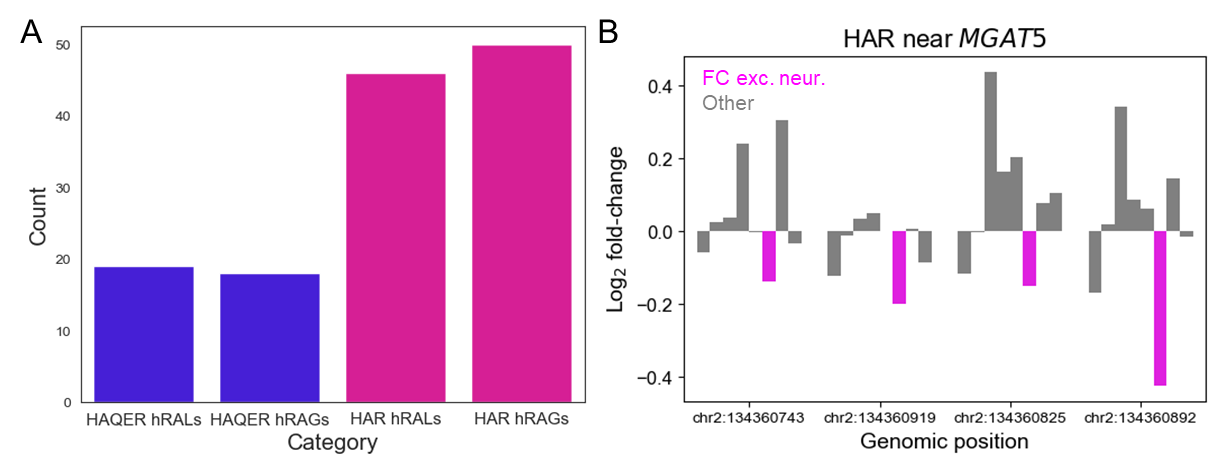

**Supplemental figure 21, related to figure 7: A)** Number of hRAGs and hRALs intersecting HARs and HAQERs. **B)** Example of an hRAL intersecting a HAR near *MGAT5*. The magenta bars show the predicted log_2_ fold-change in CA in fetal excitatory neurons and the gray bars show the same thing for the other seven cell types used to identify the hRAL.

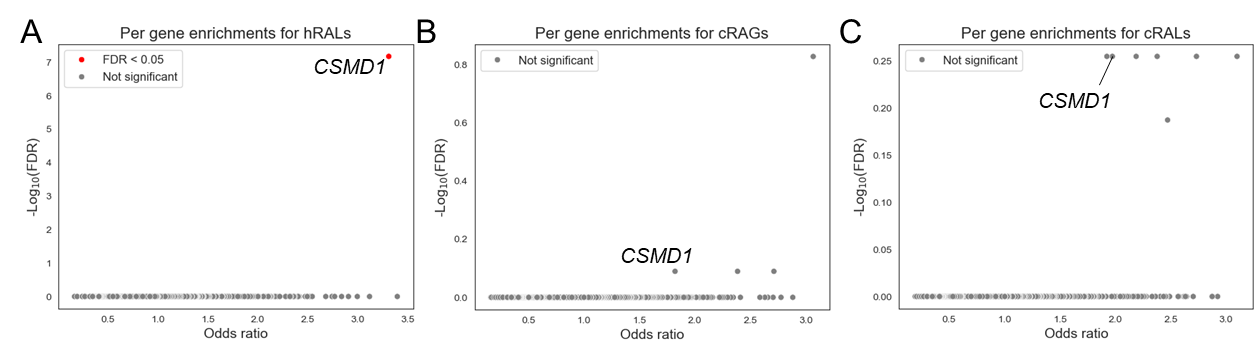

**Supplemental figure 22, related to figure 7: A)** Volcano plot showing enrichment of hRALs near individual genes (dots). The Fisher’s exact odds ratio and -log_10_(FDR) are shown on the x- and y-axes respectively. **B)** Same as in (A) but for cRAGs. **C)** Same as in (A) but for cRALs.

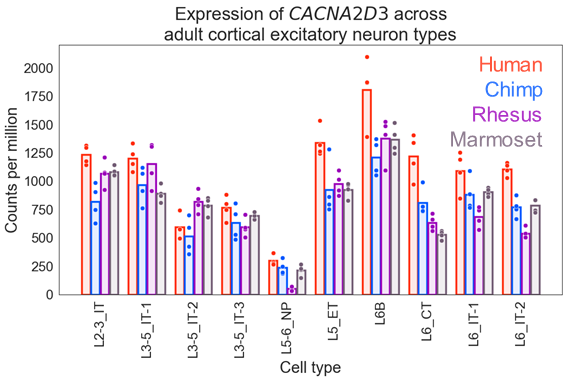

**Supplemental figure 23, related to figure 7:** Expression of *CACNA2D3* across cortical excitatory neuron types. Expression is significantly higher in human relative to chimp at an FDR < 0.05 in L2-3_IT, L5_ET, L6B, L6_CT, and L6_IT-2.

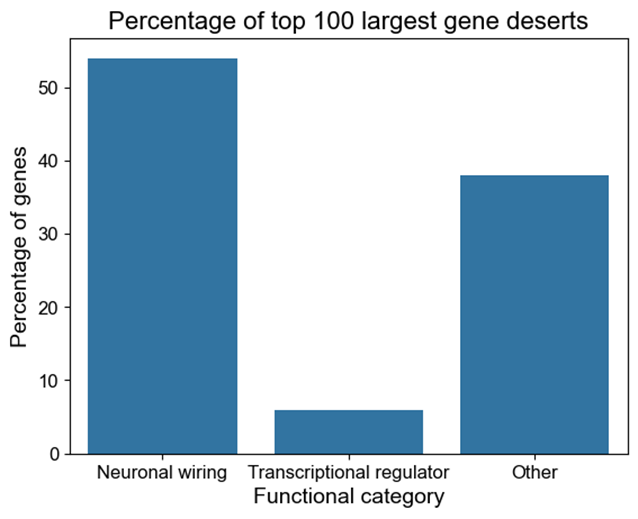

**Supplemental figure 24, related to figure 8:** Percentage of top 100 largest gene deserts assigned to genes that code for cell surface proteins involved in neuronal wiring, transcriptional regulators, or other functions. This likely severely underestimates the proportion of neuronal wiring genes as the “other” category includes genes that regulate neuronal wiring but do not code for proteins that localize to the cell surface as well as many genes that are far from other genes because they are adjacent to neuronal wiring genes in large gene deserts.
