## Supplemental Texts for "A framework to detect positive selection using variant effect predictions reveals widespread adaptive evolution of human neurons"

**Supplemental text 1: Validating the framework to detect positive selection with simulations**

Simulations have been extensively used to evaluate the performance of the MK test^1–4^ across a wide variety of different demographic scenarios. As our framework in some sense represents a generalization of the MK test, these simulations also support the utility of our method. To further test the validity of our method, we performed simulations using a well-established model of human demographic history^2,5^ implemented in SLiM^6^ v4.3 that has been used to evaluate modifications of the MK test. The core difficulty in implementing simulations for this framework is that, unlike the MK test, there is a continuous variant effect prediction rather than a binary one and that continuous variant effect prediction must be converted to selection coefficients and then back to variant effect predictions after the simulations has concluded. We therefore developed a framework to perform this mapping (Supplemental Methods).

We performed 1,000 simulations of individual 200 kb elements and then combined information across all simulations, recording the selection coefficients for each polymorphism and fixed substitution that resulted from the simulation. The results of the simulations are shown below. It is important to note that the parameter controlling the proportion of positively selected sites reflects the proportion of all sites that are positively selected and that α_Cons_ is a lower bound on the proportion of fixed substitutions with PhyloP score greater than the 95th percentile of the polymorphic PhyloP distribution that were fixed by positive selection. Therefore, although α_Cons_ should scale with the proportion of all sites that are positively selected, the two values should not be equal. Consistent with simulations for the MK test, α_Cons_ is negative (Supp. Text 1, Fig. 1A, α_Cons_ = -0.032) when there is no positive selection due to segregating mildly deleterious polymorphisms. It is near zero when 0.1% of sites are positively selected (Supp. Text 1, Fig. 1B, α_Cons_ = 0.01), and very positive when 1% of all non-neutral sites are positively selected (Supp. Text 1, Fig. 1C, α_Cons_ = 0.27). This is consistent with the idea that our method can detect positive selection and that α reflects the proportion of sites above the chosen cutoff that were fixed by positive selection.

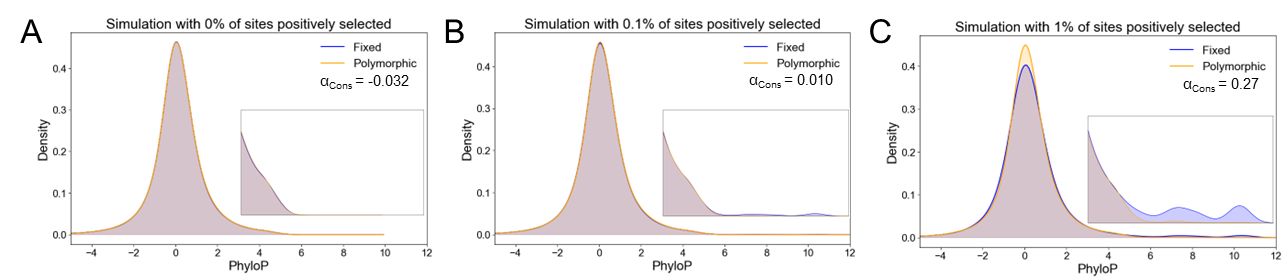

**Supplemental Text 1, Figure 1: Results of simulations. A)** The distribution of fixed and polymorphic PhyloP scores for simulations assuming 0% of sites positively selected. In all cases, the cutoff for computing α_Cons_ is the 95^th^ percentile of the polymorphic conservation score distribution. It is important to note that α_Cons_ is a lower bound on the proportion of fixed substitutions with PhyloP score greater than the 95^th^ percentile of the polymorphic PhyloP distribution that were fixed by positive selection. Therefore, although α_Cons_ should scale with the proportion of sites that were positively selected, the two values should not be equal. **B)** Same as in (A) but assuming 0.1% of all sites are positively selected. **C)** Same as in (A) but assuming 1% of all sites are positively selected.

**Supplemental text 2: Agreement with previous results**

Although simulations are important validation, another key validation is agreement with previously published results. Here, we will discuss validation that did not appear in the main text in detail and briefly mention validation that was discussed in the main text. One well-established result is that sites in viral interacting proteins (VIPs) generally experience stronger positive selection than other proteins^7^. Consistent with this, we estimate α_Cons_ = 0.29 for VIPs (Supp. Text 2, Fig. 1, very similar to the value of 0.27 using the asymptotic MK test^7^) compared to α_Cons_ = 0.053 for all proteins (Supp. Fig. 1), although this is not a significant difference, partially due to the very low number of polymorphic nonsynonymous substitutions in VIPs. Another well-established result is that there has been stronger positive selection on amino acid sites on the surface of proteins (more solvent-exposed)^4^. Consistent with this, we compute α_Cons_ = 0.060 for sites in residues exposed to solvent and α_Cons_ = -0.12 for sites in buried residues (Supp. Text 2, Fig. 2A-B), although again the lack of common polymorphic nonsynonymous sites in buried residues restricts statistical power. However, relaxing the lower DAF bound to 0.1, there is sufficient power to conclude that there has been stronger positive selection on more exposed residues (α_Cons_ 95% confidence interval for buried residues of -0.52 to -0.19 compared to α_Cons_ = -0.10 for exposed residues).

In addition to this, several results throughout the main text agree with previous empirical or theoretical results. We observe stronger evidence of positive selection on the X chromosome than the autosomes and particularly strong evidence in regions of the X chromosome previously thought to have undergone positive selection in the human lineage. We also observe stronger evidence for positive selection in house mice than in human, consistent with previous results based on the MK test.

When using ChromBPNet predictions, we find evidence for positive selection in HAQERs and HARs (both of which are thought to have undergone positive selection in the human lineage) in cell types that have previously been suggested to be particularly relevant to HAQERs and HARs. We also find evidence for stronger positive selection on substitutions with more cell type-specific effects (consistent with a large body of theoretical work) and in Alu elements relative to other parts of the genome (consistent with previous theories and findings that Alu elements are enriched for cis-regulatory divergence). Collectively, our results consistently agree with and extend previous research, providing support for the efficacy of our framework in detecting positive selection.

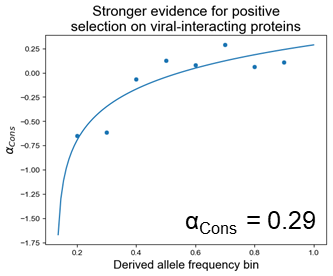

**Supplemental Text 2, Figure 1: Evidence for positive selection on viral interacting proteins (VIPs).** Result from asymptotic comparison of conservation score distribution for fixed and polymorphic nonsynonymous substitutions in VIPs. The point are at the upper limit of each derived allele frequency bin and are 0.1-0.2, 0.2-0.3, …, 0.8-0.9.

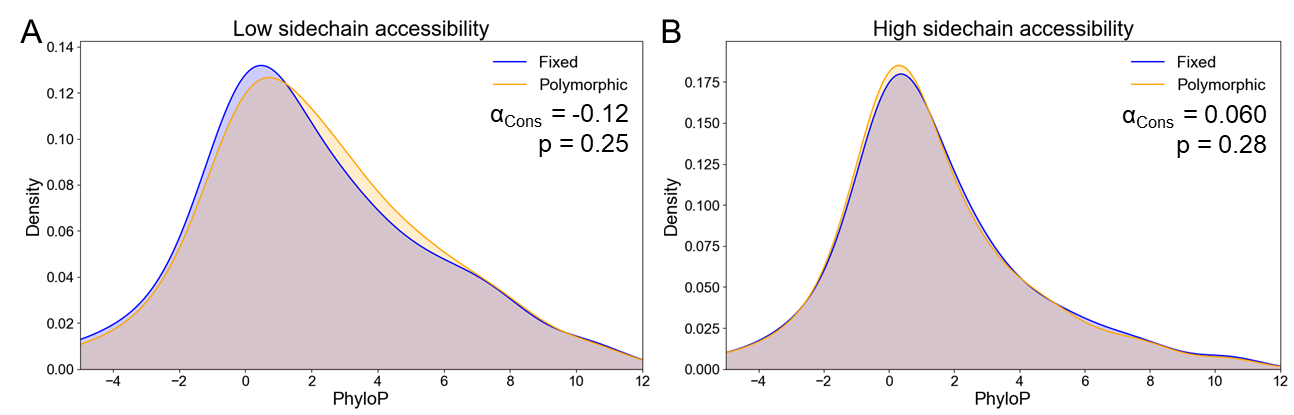

**Supplemental Text 2, Figure 2: Relationship between positive selection and solvent accessibility. A)** The genome-wide distribution of conservation scores for fixed and polymorphic nonsynonymous substitutions in residues with low sidechain accessibility. In all cases, the cutoff for computing α_Cons_ is the 60^th^ percentile of the polymorphic conservation score distribution. **B)** The same as in (A) but for high sidechain accessibility.

**Supplemental text 3: Potential causes of false positives and false negatives**

As discussed above, the similarity of our framework to the MK test enables us to borrow knowledge gained from evaluation of the MK test across different demographic scenarios. For example, our method is likely conservative when effective population size is lower in the present than in the past (leading to weak loss of constraint across the genome) as segregating polymorphisms will be more deleterious^4^. As another example, epistasis can lead to false negatives and false positives for both tests^8^.

However, there are some important distinctions between the two that are somewhat dependent on what metric is used to test for positive selection. One simple example is that when using nonsynonymous sites, our framework does not take into account synonymous sites so cannot be confounded by selection on synonymous sites in the way the MK test can. There are also more complex possibilities (see below).

**Method to test for differences between fixed and polymorphic distributions does not capture observed differences**

Second, regardless of what metric is used, it may require altering how the differences between the fixed and polymorphic distributions are evaluated. For example, in a scenario in which positive selection is primarily occurring on sites that have been recurrently positively selected throughout placental mammal evolution, we would expect an excess of sites with large negative PhyloP scores rather than large positive PhyloP scores. This is because sites with recurrent positive selection in mammals are expected to evolve faster than neutral sites, leading to negative PhyloP scores^9^. If the positive selection on these sites were to continue to occur in humans, then nonsynonymous substitutions in these sites would be more likely to fix than nonsynonymous substitutions in other sites, ultimately leading to an excess of fixed sites with negative PhyloP scores.

Alternatively, if both recurrently positively selected and highly conserved sites are under stronger positive selection than putatively neutrally evolving sites, then there may be more fixed sites at either tail of the PhyloP score distribution. An example of this can be seen looking at the PhyloP distribution for sites in VIPs (Supp. Text 3 Fig. 1A). There is a fatter tail of fixed sites with negative PhyloP scores relative to polymorphic sites, but also a somewhat fatter tail of fixed sites with large positive PhyloP scores (Supp. Text 3 Fig. 1A). Consistent with this, we observe α_Cons_ = 0.2 (p = 0.11, one-sided Fisher’s exact test) when removing sites with PhyloP less than -2 (Supp. Text 3 Fig. 1B). Furthermore, when removing sites with PhyloP greater than 2 and testing for an excess of fixed sites with large negative PhyloP, we find strong evidence for positive selection (Supp. Text 3 Fig. 1C, α_Cons_ = 0.51, p = 0.014, one-sided Fisher’s exact test). These results, in conjunction with the asymptotic results for VIPs (Supp. Text 2, Fig. 1) suggest that there has been positive selection on recurrently selected sites in VIPs as well as in highly conserved sites in the human lineage.

Interestingly, sites in VIPs with large negative PhyloP (less than -3) were enriched in regulators of phagocytosis relative to sites with positive PhyloP (FDR = 0.03, odds ratio = 11.2, Fisher’s exact test). There is also evidence for positive selection on these phagocytosis genes with the MK test as well (p = 0.045, odds ratio = 7.27). Many of the genes contributing to the enrichment (*CD36*^10,11^, *MARCO*^12^, *LDLR*^13^, *MSR1*^14^) are associated with Alzheimer’s disease and/or atherosclerosis, raising the possibility that recurrent positive selection on these genes due to viral pressure may have increased risk for these diseases in the human lineage. Furthermore, this demonstrates that careful choice of methods used to compare fixed and polymorphic distributions of predicted variant effects can provide important insight into positive selection.

**Loss or gain of constraint**

In addition to the possibility of genome-wide loss of constraint discussed above, it is also possible that more localized loss or gain of constraint can respectively lead to false negatives or false positives for both our framework and the MK test. As a specific example, if a *cis*-regulatory element were to be highly conserved throughout mammalian evolution but lose function (possibly as the result of positive selection) in the human lineage 2 million years ago, then it would have been evolving under constraint for much of human evolution but be under no constraint when polymorphisms arose, leading to an excess of polymorphisms at highly conserved sites in that element. We anticipate that this may have a larger effect in *cis*-regulatory elements relative to protein coding genes as it is more difficult to identify *cis*-regulatory elements that have lost function than it is for protein coding genes (which usually have an early stop codon if they have entirely lost function and so can be identified as pseudogenes and excluded from further analysis).

Alternatively, if a previously neutrally evolving repetitive element were to gain *cis*-regulatory activity (possibly as the result of positive selection) that provided a fitness advantage 2 million years ago, then it would be evolving under constraint when polymorphisms arose but not when many variants became fixed. The latter scenario need not be the result of positive selection, since the element may have gained activity through neutral evolution and then become constrained due to loss of function in another *cis*-regulatory element that the newly active element was now compensating for. Overall, although it is difficult to know the extent to which local loss or gain of constraint confounds the results of the MK test or our framework, it is worth keeping these in mind as potential alternative explanations for either the presence or absence of evidence for positive selection.

**Background selection**

Another potential cause of false positives or false negatives is differences in the strength of background selection across the genome^15^. Background selection (BGS) is usually defined as the loss of neutral diversity as a result of negative selection on linked sites. As a result of this, there may be fewer polymorphisms relative to fixed substitutions in regions of the genome with stronger background selection (although it is very difficult to test this as measurements of the strength of background selection are based on human polymorphism data)^15^. If polymorphisms in regions with stronger background selection tend to be in neutral sites more often than in regions with weaker background selection, that could lead to false negatives. On the other hand, if those polymorphisms tend to be in non-neutral sites more often than in regions with weaker background selection, that could lead to false positives.

To test the potential role of background selection in driving our results, we reasoned that if BGS were playing a role in driving our results, then it would weaken the correlation between the estimated strength of positive selection (in this case the -log_10_(Fisher’s exact p-value) multiplied by the sign of α) estimated for the *cis*-regulatory neighborhood of a gene and the average estimated strength of positive selection across 25 kb windows tiling the *cis*-regulatory neighborhood of that gene. To see why, we can consider an extreme example in which a gene has two 25 kilobase regions. One region has no functional importance and so has no background selection and contains a mix of fixed and polymorphic sites that are all neutral. The other region is extremely conserved, has extremely strong background selection, and many fixed substitutions and a few polymorphic substitutions in conserved sites as a result. If we were to test for positive selection in each region individually, we would expect α to be approximately 0 in both regions with p-values close to 1. On the other hand, if we were to aggregate sites from both regions together, we would see an excess of fixed substitutions in conserved sites, α greater than 0, and a low p-value (this is an example of Simpson’s paradox, though it is actually more of an unintuitive statistical effect than a true paradox, and also can occur when using the MK test^16^).

However, random noise will also decrease the correlation between the per-window average estimate and aggregated estimate. Therefore, we compare the correlation between the per-window average estimate and the aggregated estimate with the correlation after shuffling the positions of the sites, as this will eliminate any effects of BGS on the difference between the per-window average estimate and the aggregated estimate. Consistent with the idea that BGS is not playing a major role in causing false negatives or false positives, we find Pearson’s rho = 0.587 using the actual data and Pearson’s rho = 0.594 using the shuffled data when using PhyloP scores as input (Supp. Text 3 Fig. 2A-B). In addition, using predicted log_2_ fold-change in CA in fetal cortical neurons, we observe Pearson’s rho = 0.67 using the actual data and Pearson’s rho = 0.71 using the shuffled data (Supp. Text 3 Fig. 2C-D) and this is generally true across all different cell types (Supp. Text 3 Fig. 2E). Overall, this analysis suggests that BGS is unlikely to play a major role in generating either false negatives or false positives, although we cannot fully rule out that BGS could affect our results.

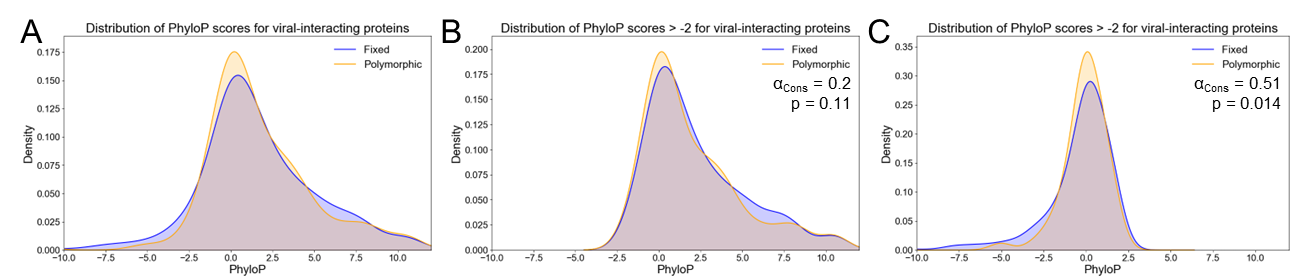

**Supplemental Text 3, Figure 1: Viral interacting proteins (VIPs) as an example of positive selection on conserved and rapidly evolving sites. A)** The genome-wide distribution of conservation scores for fixed and polymorphic nonsynonymous substitutions VIPs. **B)** The same as in (A) but removing sites with PhyloP less than -2. In all cases, the cutoff for computing α_Cons_ is the 60^th^ percentile of the polymorphic conservation score distribution. **C)** The same as in (A) but removing sites with PhyloP greater than 2 and testing for enrichment for negative PhyloP scores in the fixed distribution relative to the polymorphic distribution.

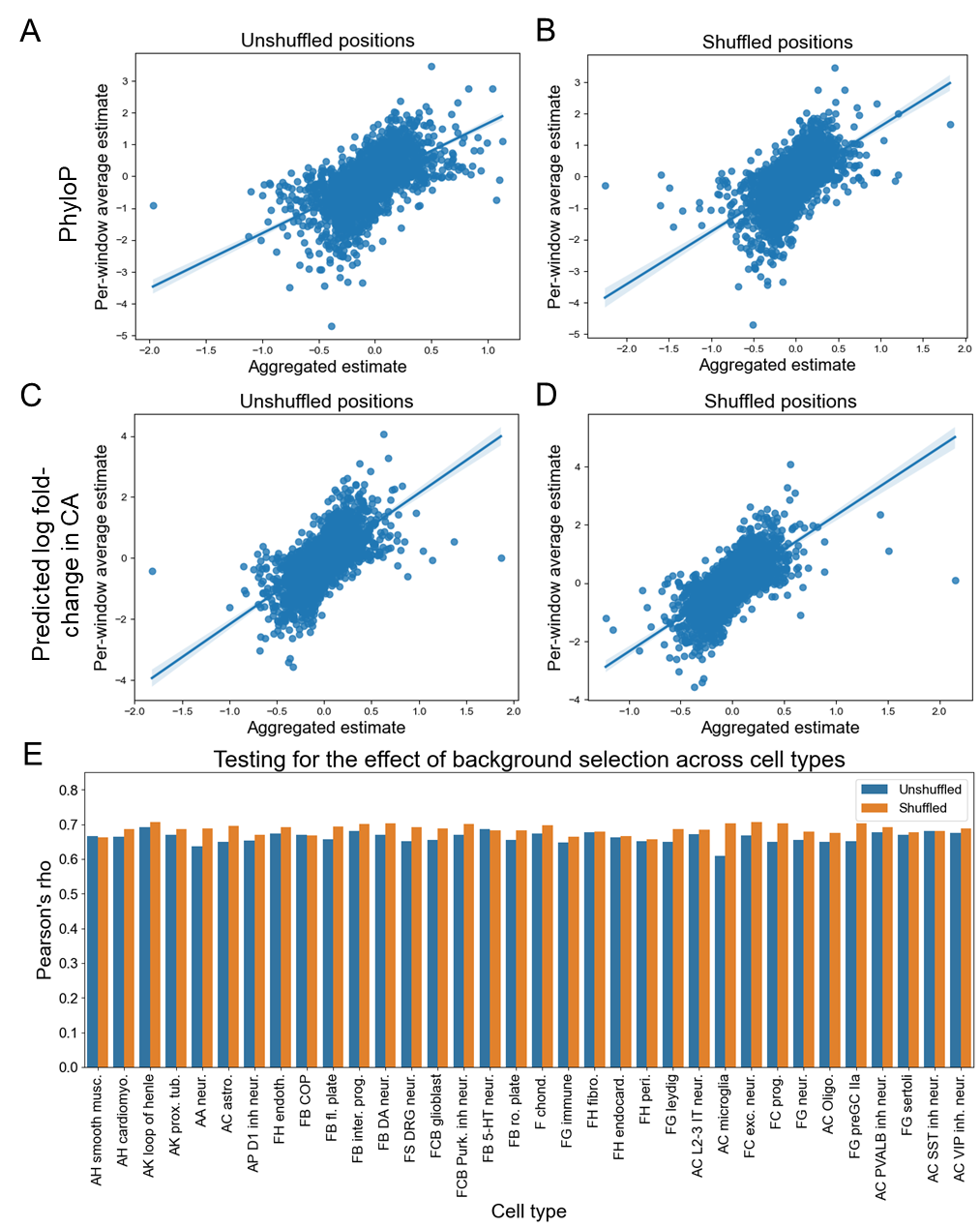

**Supplemental Text 3, Figure 2: Evaluating potential effects of background selection. A)** Scatterplot with the –log_10_(Fisher exact p-value) multiplied by the sign of α_Cons_ for each gene, termed the aggregated estimate, on the x-axis and the average –log_10_(Fisher exact p-value) multiplied by the sign of α_Cons_ across 25 kilobase windows tiling the region assigned to each gene, per gene. Line represents the line of best fit and 95% confidence interval. **B)** The same as in (A) but computed after shuffling which window each site was assigned to. **C)** The same as in (A) but for predicted absolute log fold-change in CA in fetal cortical excitatory neurons. **D)** The same as in (B) but for predicted absolute log fold-change in CA in fetal cortical excitatory neurons. **E)** Comparison of correlation between aggregated estimate and per-window average estimate with shuffled and unshuffled positions across cell types.

**Supplemental text 4: Positive selection on nonsynonymous substitutions in neuron projection genes.**

As there are an insufficient number of nonsynonymous polymorphisms in this set of neuron projection genes (there are 19) to apply our framework, we used the MK test^1^ (see Supp. Text 5 for a comparison of the MK test and our framework when applied to nonsynonymous sites). First, and consistent with the results in the text, we find evidence for positive selection on haploinsufficient genes relative to genes that are not haploinsufficient (odds ratio = 1.01, 95% confidence interval of 0.89-1.14, one-tailed Fisher’s exact test for haploinsufficient genes compared to odds ratio = 0.86 genes that are not haploinsufficient). However, relative to the evidence for positive selection for haploinsufficient nonsynonymous-depleted genes (Fig. 1E, Supp. Fig. 2C, left, odds ratio = 1.04, p = 0.33, one-tailed Fisher’s exact test), we found a significant excess of nonsynonymous changes in the neuron projection subset (Fig. 1E, Supp. Fig. 2C, right, odds ratio = 1.65, p = 0.038, one-tailed Fisher’s exact test) suggesting that there has been particularly strong positive selection on the neuron projection gene subset. Because many of these genes are associated with autism spectrum disorder (ASD, 54 out of 117 of the neuron projection genes have been linked to ASD^17^) and we have previously shown that there was polygenic positive selection for downregulation of ASD-associated genes in the human lineage^18^, we tested whether the neuron projection subset was also disproportionately downregulated. Indeed, we observe a bias toward downregulation in both adult human layer 2/3 intratelencephalic neurons^19^ (Supp. Text 4, Fig. 1C, p = 0.027, binomial test) and human-chimp hybrid cortical organoids^20^ (Supp. Text 4, Fig. 1D, p = 0.13 at an FDR cutoff for differential expression of 0.05, Supp. Text 4, Fig. 1E-F, p = 0.03, binomial test at an FDR of 0.1). The bias in the hybrid cortical organoids in particular suggests that there may have been positive selection on the expression of this set of neuron projection genes as well^21^.

Finally, we wanted to determine whether this positive selection was more likely to affect neuronal projection development (e.g. by altering the wiring of the brain) or adult neuronal function. To do this, we evaluated whether the expression of these neuron projection genes was enriched for genes that peak in expression during late human fetal cortical neuronal development or in adult neurons^22^. Strikingly, there was a strong enrichment for genes that peak during late fetal development relative to adult neurons (24 out of 117 genes peak in trimester 2 or trimester 3 in at least one neuronal cell type, odds ratio = 7.5, p = 0.023, one-tailed Fisher’s exact test), suggesting that the positive selection we have detected is most likely to affect neuronal projection development. To further test this, we used the MK framework to test for positive selection on only the subset of neuron projection genes with peak expression during late fetal development compared to the rest of the neuron projection genes. While there was evidence for positive selection on the genes that peak in expression during late fetal development (Supp. Text 4, Fig. 1G, odds ratio = 7.3, p = 0.0191, one-tailed Fisher’s exact test), there was no such evidence for the other neuron projection genes (Supp. Text 4, Fig. 1G, odds ratio = 1.33, p = 0.34, one-tailed Fisher’s exact test). As a specific example, EPHB1, which plays a key role in axon guidance in and peaks in expression during mid to late embryonic development of deep layer cortical neurons in both mice and humans^23^, contains a fixed human-chimp nonsynonmous substitution (I976M) in a base that is conserved across all other placental mammals in the alignment. Collectively, this points to positive selection on protein-coding sites in genes that regulate the development of neuronal connectivity and provides a rare example of positive selection acting on both the coding and non-coding genome of the same set of genes.

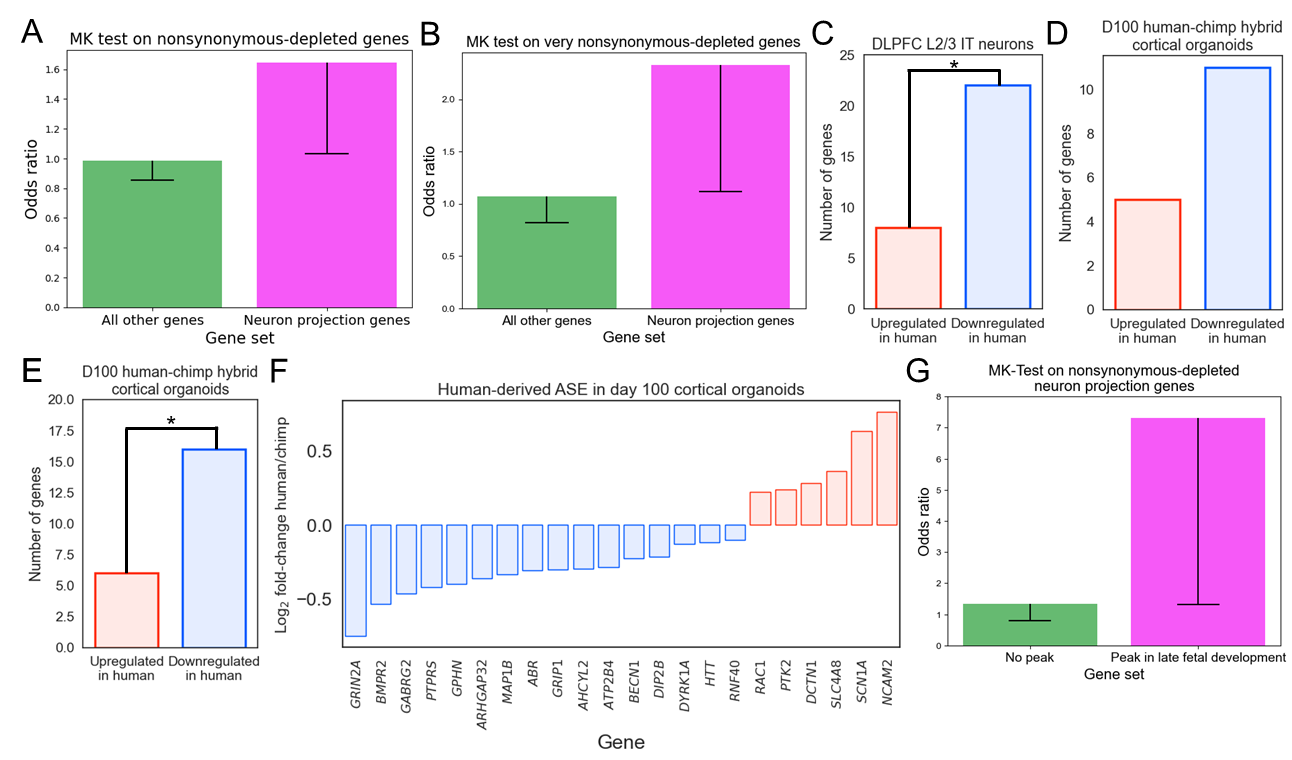

**Supplemental Text 4, Figure 1: A)** Odds ratios and one-sided confidence interval for the MK-test run on haploinsufficient, nonsynonymous depleted genes involved in neuronal projections compared to all haploinsufficient, nonsynonymous depleted genes. Magenta color indicates p < 0.05, green indicates p > 0.05. **B)** Same as in (A) but for very nonsynonymous-depleted genes. **C)** Number of downregulated and upregulated genes in dorsolateral prefrontal cortex (DLPFC) layer 2/3 intratelencephalic neurons. **D)** Same as in (C) but comparing human and chimp alleles in hybrid cortical organoids. **E)** Same as in (D) but using genes with FDR < 0.1 comparing expression from the human and chimp allele to increase power. **F)** Differential expression of genes from (E). All genes have FDR < 0.1. **G)** Odds ratios and one-sided confidence interval for the MK-test run on haploinsufficient, nonsynonymous depleted genes involved in neuronal projections with peak expression in late fetal development compared to those that do not peak in late fetal development.

**Supplemental text 5: Disagreement between the MK test and testing for positive selection with PhyloP scores**

In some cases, testing for positive selection using our framework in conjunction with PhyloP scores can detect positive selection when the MK test might not, particularly in cases where the effective population size has decreased. It has previously been shown that a decrease in effective population size results in an increase in the number of deleterious polymorphisms^4,24^. Importantly, it is expected that there will be an increase in the number of polymorphic mildly deleterious substitutions, but a disproportionately much smaller increase for strongly deleterious substitutions^24^. As an extreme example, substitutions that are embryonic lethal will remain at an allele frequency of 0 even with a massive decrease in effective population size. This increase in the number of mildly deleterious polymorphisms is expected to increase the number of polymorphic nonsynonymous substitutions relative to the number of polymorphic synonymous substitutions, potentially leading to false negatives when using the MK test to detect positive selection. However, when comparing the distributions of PhyloP scores, we might expect that while there could be an increase in the number of polymorphisms in sites with moderately high PhyloP scores, there would be very little change in the number of polymorphisms in sites with very high PhyloP scores as these likely have large, mostly negative effects on fitness. This is consistent with what we observe in simulations (Supp. Text 1 Fig. 1A-C) in which the decreasing effective population size leads to an excess of polymorphisms in sites with moderately high PhyloP scores, without affecting the number of polymorphisms in sites with very high PhyloP scores. We think this is the most likely explanation for the difference in results between the MK test and our framework for genes associated with cardiac arrest (Fig. 1E) and suggests that this framework may be particularly useful in species which had recent decreases in effective population size.

There are also situations in which the MK test will be able to detect positive selection but our framework will not. For example, if there are many fixed nonsynonymous substitutions and very few polymorphic nonsynonymous substitutions then our framework would not be applicable but there could be very strong evidence for positive selection using the MK test (e.g. neuron projection genes in Supp. Text 4). In addition, if the variant effect prediction being used is a poor predictor of the effects of nonsynonymous substitutions, then our framework would likely produce a false negative result. For example, many testis-specific genes are evolutionarily new or part of large families of highly similar proteins, potentially making conservation a poor predictor of variant effect.

**Supplemental text 6, sensitivity analyses:**

In the majority of our analyses, the primary parameter to be chosen is the percentile cutoff of the polymorphic distribution of variant effect predictions (e.g. PhyloP scores) to use to divide the sites into putatively neutral and non-neutral. In some sense, this parameter should be chosen based on a combination of what the user thinks the cutoff should be based on the underlying biology of the statistic being compared for fixed and polymorphic sites, what the user is interested in testing, and what is feasible given the number of sites available. In general, a larger number of sites enables the use of a stricter cutoff since, all else being equal, power is maximized for Fisher’s exact test with an equal number of sites in the two rows of the 2x2 table (Supp. Text 6 Fig. 1). To illustrate this in practice, we are able to choose a stringent cutoff (95th percentile) when investigating all non-coding sites genome-wide as there are millions of sites. However, we pick a less stringent cutoff when testing individual gene sets, which generally only have a few thousand sites.

However, it is important to determine whether our results, such as the evidence for positive selection across different sets of regions or cell types that we would like to compare, are not overly sensitive to this cutoff. Although there are inevitably quantitative differences that result based on what cutoffs are chosen, the results are generally qualitatively insensitive to this cutoff. For example, we observe strong positive correlations between the effect size estimate from MASHR using either the 60th percentile or the 80th percentile of the polymorphic distribution as a cutoff for both genes and gene sets (Supp. Text 6 Fig. 2A-D, all p < 10^-71^). Similarly, our results for HARs and HAQERs are consistent at different cutoffs (Supp. Text 6 Fig. 3A-D, all p < 0.001). Finally, we observe similar results for positive selection on cell type-specific substitutions whether using the 60^th^ or the 80^th^ percentile of the polymorphic distribution as a cutoff (Supp. Text 6, Fig. 4A-B). Collectively, these results show that although it is important to select a cutoff to compute α with some care, our results are generally robust to changing this cutoff.

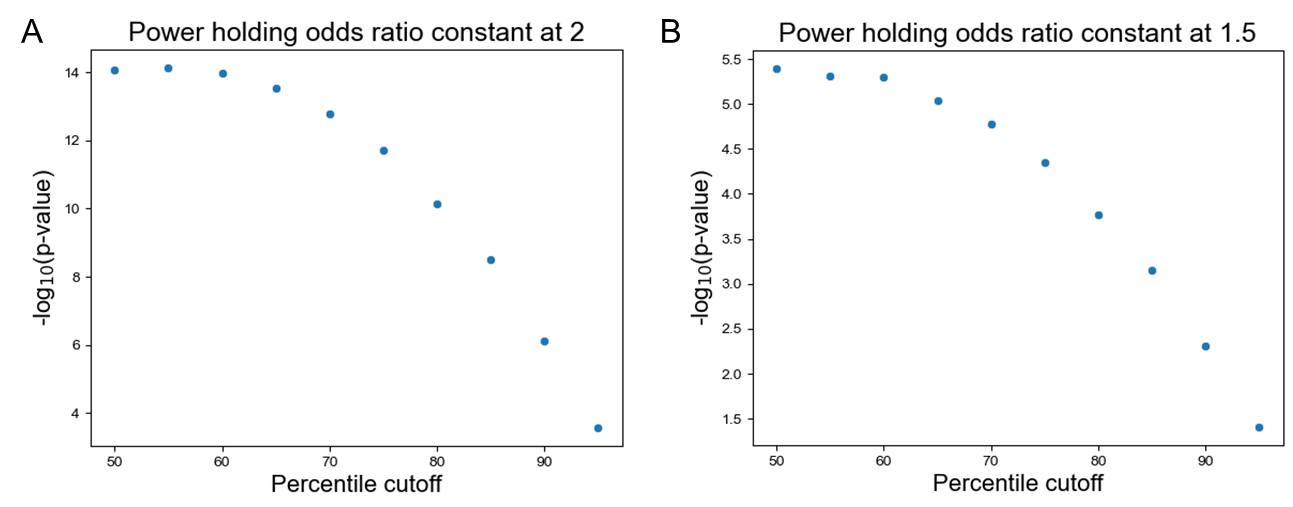

**Supplemental Text 6, Figure 1: Exploration of power of test across percentile cutoffs. A)** Scatterplot of the percentile cutoff of the polymorphic distribution used to bin sites compared to the –log_10_(p-value) for Fisher’s exact test holding the odds ratio constant at 2. Power decreases as the cutoff increases. **B)** Same as in (A) but for odds ratio of 1.5.

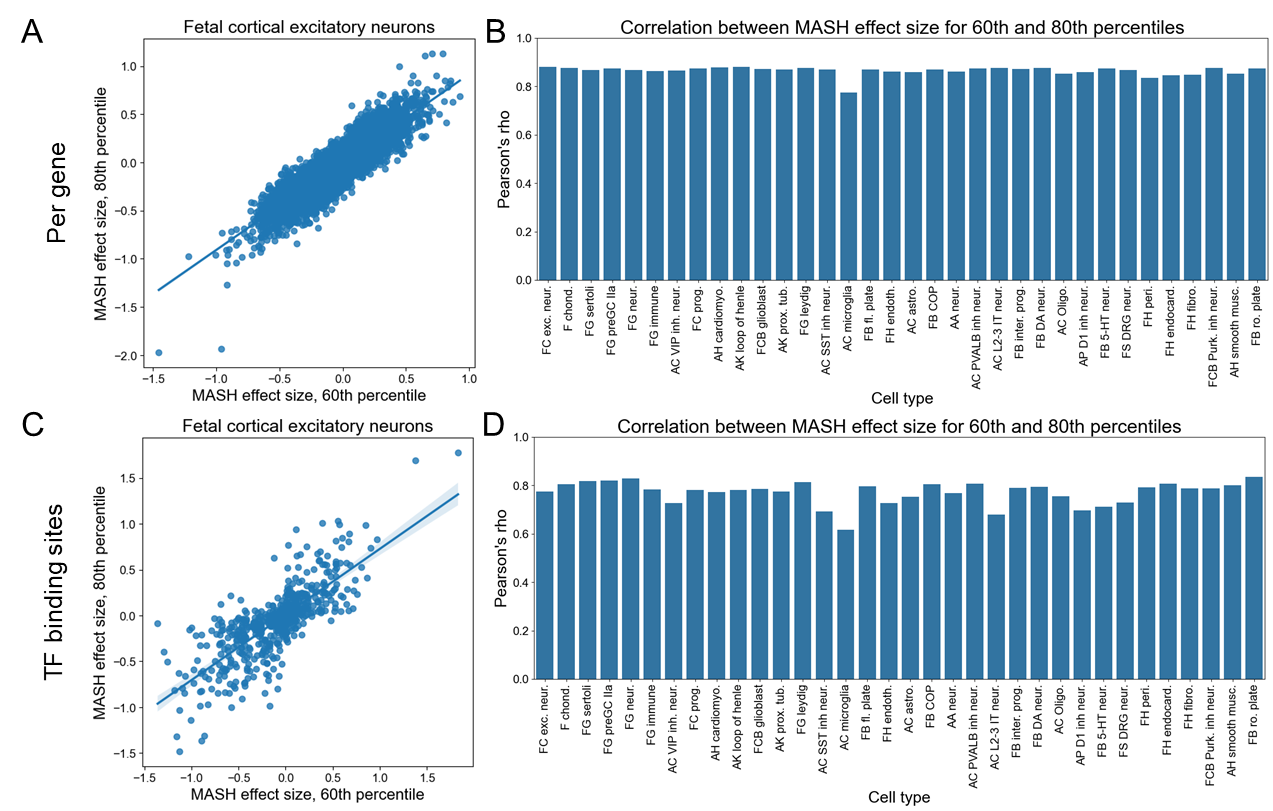

**Supplemental Text 6, Figure 2: Sensitivity analyses for genes and transcription factor binding sites. A)** Scatterplot of the per gene MASH effect size using the 60^th^ percentile of the polymorphic as a cutoff compared to the 80^th^ percentile of the polymorphic distribution as a cutoff for fetal cortical excitatory neurons. The line represents the line of best fit and 95% confidence interval. Each point represents a gene. **B)** The Pearson correlation coefficients for the comparison in (A) all cell types used in this study. **C)** Same as in (A) but each point represents a transcription factor binding sites. **D)** Same as in (B) but for transcription factor binding sites.

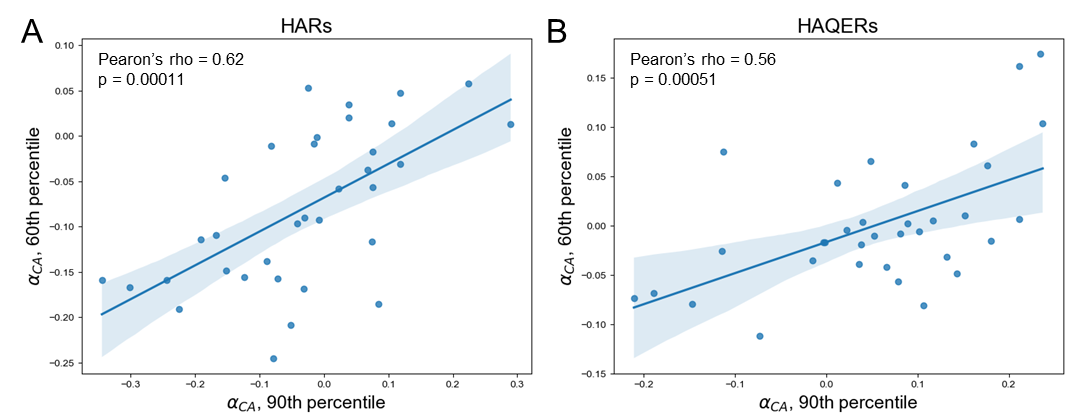

**Supplemental Text 6, Figure 3: Sensitivity analyses for HARs and HAQERs. A)** Scatterplot of the α_Cons_ using the 60^th^ percentile of the polymorphic as a cutoff compared to the 80^th^ percentile of the polymorphic distribution for HARs. Each point represents a cell type. **B)** The same as in (A) but for HAQERs.

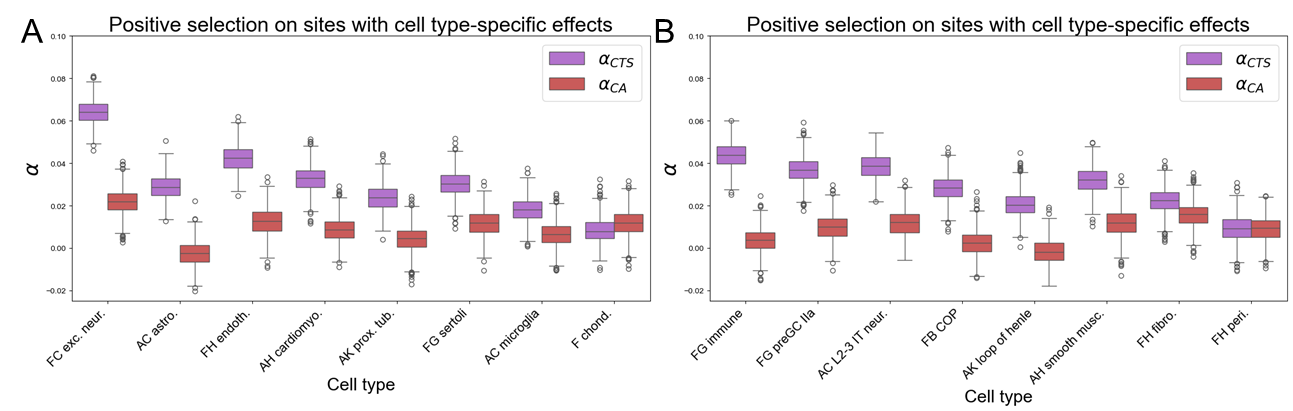

**Supplemental Text 6, Figure 4: Sensitivity analyses for selection on substitutions with cell type-specific effects. A)** Box plot comparing α_CTS_ and α_CA_ across cell types based on 1,000 bootstraps. The cutoff for computing α is the 80^th^ percentile of the polymorphic score distribution (compare with Fig. 6C). **B)** Same as in (A) but for the second group of cell type (compare with Supp. Fig. 16).

**Supplemental Methods**

**Population genetics simulations**

In simulation studies of the MK test, it is common practice to assign synonymous sites a selection coefficient of 0, most nonsynonymous sites having either a weakly or strongly deleterious (negative) selection coefficient, and a small fraction of nonsynonymous sites having a positive selection coefficient, with the actual values for those coefficients selected based on previous work^2^.

We attempted to perform simulations in a similar manner. The primary obstacle in doing so is mapping PhyloP scores to selection coefficients. Although the relationship between these two quantities is unknown, it is very likely that sites with higher PhyloP scores or larger effects on CA have higher absolute selection coefficients. We focused on PhyloP scores for our simulations as we computed those genome-wide and attempted to assign absolute selection coefficients to them in a reasonable manner (based on previous work) that is compatible with SLiM^6^ (though we expect that this would yield highly similar results if based on predicted effects on chromatin accessibility). To do this, we restricted only to sites with greater than 250 species in the alignment and then split sites into 8 bins numbered 1-8: those with PhyloP scores less than 1, from 1-2, 2-3, 3-4, 4-5, 5-6, 6-7.5, and greater than 7.5. The last two bins were chosen based on the somewhat discrete distribution of PhyloP scores, as highly conserved sites generally have scores close to 7.4, 8.5, and 10.5 when using our model file and the 447-way alignment. We then calculated the number of sites in each of these bins which resulted in approximately 80% of sites assigned to bin 1, 11% to bin 2, 4% to bin 3, 1% to bin 4, 0.5% to bin 5, 0.5% to bin 6, 0.8% to bin 7, and 0.7% to bin 8.

To map those bins onto absolute selection coefficients (|s|), we considered anything in bin 1 to be neutral (i.e. |s| = 0)). In previous work, weakly selected sites were assigned absolute selection coefficients from an exponential distribution with parameter |s| = 10/(2*N_e_) and strongly selected sites were assigned with parameter |s| = 500/(2*N_e_) (where N_e_ is the effective population size) so we used those values as a guide when assigning absolute selection coefficients to the remaining bins^2^. We used the following absolute selection coefficients as the parameter for an exponential distribution for bins 2-5, in order these were: 5/(200*N_e_), 5/(20*N_e_), 5/(200*N_e_), 20/(2*N_e_). For the more conserved sites in bins 6 to 8, we drew absolute selection coefficients from a normal distribution with mean 50/(2*N_e_), 125/(2*N_e_), and 500/(2*N_e_) and standard deviation equal to the mean divided by 10, which ensures that the values drawn from these distributions are extremely unlikely to be negative.

Although it is not possible to perfectly capture the complexity of the genome, we attempted to capture the interspersed nature of *cis*-regulatory elements by constructing a 200 kb region consisting of two types of genomic elements. A “mostly neutral” type that covered the first and last 63 kb as well as the 10 kb between each of the “mostly conserved” type that were 500 bp in length and evenly distributed in the central 64 kb. We controlled the proportion of positive selection using the parameter pb such that either 0%, approximately 0.1%, or 1% of non-neutral sites had positive selection coefficients. As a result of this genomic architecture, we expect our simulations to capture the effects of background selection, with stronger background selection in the middle of the 200 kb element. For further details of how these genomic elements were constructed in SLiM^6^, please refer to: https://github.com/astarr97/PosSelect/tree/main/Simulations.

After the simulations had finished running, we then reassigned those selection coefficients back to PhyloP scores based on the properties of the distributions from which the selection coefficients were drawn. For example, absolute selection coefficients greater than 0.017 (which corresponds to 5 standard deviations on either side of the mean of the bin 8 normal distribution) were assigned to bin 8. For further details of the reassignment, please refer to: https://github.com/astarr97/PosSelect/tree/main/Simulations. After assigning absolute selection coefficients back to PhyloP scores, we then compared the PhyloP score distributions for fixed and polymorphic sites and computed α_Cons_ as described in the main text, using the 95th percentile of the polymorphic PhyloP distribution as the cutoff.

**Testing agreement with previous results**

To test for positive selection on viral-interacting proteins (VIPs), we restricted to nonsynonymous substitutions in the proteins from Enard et al. 2016^7^ and then computed the asymptotic estimate of α_Cons_ as described in the main text, using the 60th percentile of the polymorphic PhyloP distribution as the cutoff. To compare the strength of positive selection on residues that are more or less solvent-exposed, we used the genome-wide predictions of amino acid side-chain accessibility (pPSE), a metric of amino acid exposure to solvent, from White et al. 2023^25^. We first used ProtVar^26^ to map the genomic positions of the nonsynonymous substitutions to Uniprot^27^ protein IDs and positions. We then joined the PhyloP information with the pPSE estimates using the uniprot protein positions and computed α_Cons_ as described in the main text, using the 60th percentile of the polymorphic PhyloP distribution as the cutoff.

**Exploring positive selection on rapidly evolving and conserved sites in VIPs**

We restricted to sites in VIPs and tested for positive selection using the 60th percentile of the polymorphic PhyloP distribution as the cutoff for all VIP results. First, we restricted only to sites with PhyloP greater than -2 and computed α_Cons_ as well as a p-value using a one-sided Fisher’s exact test. We next restricted to only sites with PhyloP less than 2, flipped the sign of the PhyloP scores, and tested for positive selection on sites with initially negative PhyloP scores.

**Evaluating the effects of background selection on our results**

We first restricted to genes whose substitution (fixed or polymorphic) sites spanned a minimum of 150 kilobases. We then tiled the region of the genome assigned to that gene with 25 kilobase windows and computed α and a p-value with Fisher’s exact test for all windows with at least 20 fixed and 10 polymorphic sites. We then repeated this computation for each window after shuffling which window each site was assigned to. We performed this procedure both for PhyloP scores and CA for each of the 34 cell types separately. We then computed the -log_10_(Fisher’s exact p-value) multiplied by the sign of α as a measure of the signed evidence for positive selection at the gene level and for each window, and then took the mean of that value across windows for each gene. We then computed the Pearson correlation between the per gene mean across windows and the value for each gene using the unshuffled and shuffled data separately.

**Exploring positives selection on nonsynonymous substitutions in neuron projection genes**

To explore which genes were driving this result, we used GOrilla^28^ to test enrichment of haploinsufficient, nonsynonymous-depleted genes with at least one fixed or polymorphic nonsynonymous substitution relative to all haploinsufficient genes with at least one fixed or polymorphic nonsynonymous substitution. We downloaded the set of genes in the neuron projection category of the cellular component ontology from GOrilla and used that as the set of neuron projection genes for all further analyses. We restricted to that set of neuron projection genes and then used the MK test^1^ to test for positive selection on nonsynonymous sites in that set of genes as well as all haploinsufficient, nonsynonymous-depleted genes. We repeated this analysis with very nonsynonymous-depleted genes. We then restricted to genes with Benjamini-Hochberg^29^ FDR < 0.05 when comparing human expression to chimp expression in DLPFC L2/3 IT neurons and used the binomial test with background probability equal to the proportion of genes with FDR < 0.05 that have lower expression in human to test for a bias toward higher or lower expression in human in that gene set. We repeated this analysis for day 100 human-chimp hybrid cortical organoids using genes with FDR < 0.05 and genes with FDR < 0.1. We took all genes that peak in expression in fetal trimester 2 or trimester 3 and genes that peak at any point during postnatal development^22^ in any neuronal type and intersected that set of genes with the neuron projection genes. We then used Fisher’s exact test to test for enrichment for genes that peak during fetal development. We then restricted to the set of genes that peak during fetal development and ran the MK test.

**Exploring the sensitivity of results to choice of the polymorphic distribution cutoff**

To explore the effect of using a more stringent percentile cutoff on statistical power, we held the odds ratio constant at 2 or 1.5 with 1000 polymorphic or 1000 fixed sites across different percentile cutoffs for the polymorphic distribution. For example, if the cutoff was 0.6, the 2x2 table would be [[600, 488], [400, 492]] for an odds ratio of 1.5. In all cases, we compared the polymorphic distribution cutoff used in the main text (60th percentile for all except HARs and HAQERs for which we used the 90th percentile) to a different cutoff (80th percentile for all except HARs and HAQERs, 60th percentile for HARs and HAQERs). We used seaborn regplot with default parameters to create scatterplots and lines of best fit and used the implementation of the Pearson correlation from scipy^30^ v1.14.1.
